# Interdisciplinary analyses of Bronze Age communities from Western Hungary reveal complex population histories

**DOI:** 10.1101/2022.02.03.478968

**Authors:** Dániel Gerber, Bea Szeifert, Orsolya Székely, Balázs Egyed, Balázs Gyuris, Julia I. Giblin, Anikó Horváth, Kitti Köhler, Gabriella Kulcsár, Ágnes Kustár, István Major, Mihály Molnár, László Palcsu, Vajk Szeverényi, Szilvia Fábián, Balázs Gusztáv Mende, Mária Bondár, Eszter Ari, Viktória Kiss, Anna Szécsényi-Nagy

**Author notes:** Correspondence to: Eszter Ari, Viktória Kiss, Anna Szécsényi-Nagy. These authors jointly supervised this work (Eszter Ari, Viktória Kiss, Anna Szécsényi-Nagy).

## Abstract

In this study we report 21 ancient shotgun genomes from present-day Western Hungary, from previously understudied Late Copper Age Baden, and Bronze Age Somogyvár-Vinkovci, Kisapostag, and Encrusted Pottery archaeological cultures (3530 – 1620 cal BCE). Our results indicate the presence of high steppe ancestry in the Somogyvár-Vinkovci culture. They were then replaced by the Kisapostag group, who exhibit an outstandingly high (up to ∼47%) Mesolithic hunter-gatherer ancestry, despite this component being thought to be highly diluted by the time of the Early Bronze Age. The Kisapostag population contributed the genetic basis for the succeeding community of the Encrusted pottery culture. We also found an elevated hunter-gatherer component in a local Baden culture associated individual, but no connections were proven to the Bronze Age individuals. The hunter-gatherer ancestry in Kisapostag is likely derived from two main sources, one from a Funnelbeaker or Globular Amphora culture related population and one from a previously unrecognised source in Eastern Europe. We show that this ancestry not only appeared in various groups in Bronze Age Central Europe, but also made contributions to Baltic populations. The social structure of Kisapostag and Encrusted pottery cultures is patrilocal, similarly to most contemporaneous groups. Furthermore, we developed new methods and method standards for computational analyses of ancient DNA, implemented to our newly developed and freely available bioinformatic package. By analysing clinical traits, we found carriers of aneuploidy and inheritable genetic diseases. Finally, based on genetic and anthropological data, we present here the first female facial reconstruction from the Bronze Age Carpathian Basin.

## Introduction

Several studies have addressed major population historical events in prehistoric Europe regarding pre-Neolithic hunter-gatherers (HG) (Fu et al., 2016; Mittnik et al., 2018; Rivollat et al., 2020), their assimilation to early European farmers during the Neolithic era (Haak et al., 2015; Lipson et al., 2017; Mathieson et al., 2018; Mittnik et al., 2018), and the appearance, expansion and admixture of steppe ancestry between the Eneolithic / Late Copper Age and the dawn of Early Bronze Age (Allentoft et al., 2015; Haak et al., 2015; Olalde et al., 2018; Papac et al., 2021). Despite these large scale genetic studies are inevitable to understand the roots of the European gene pool, there are currently only a few studies that uncover regional interaction or social stratification using biological relatedness analyses (Freilich et al., 2021; Schroeder et al., 2019; Žegarac et al., 2021), especially in Hungary during the Copper (∼4500-2700 BCE) and Bronze (∼2700-800 BCE) Ages. A number of cultural transformations occurred in the Carpathian Basin, often attributed to population changes and genetic influxes. The observed shifts in ancestry compositions sparsely covered intensive or direct european HG introgressions into early european farmer (EEF) or steppe ancestry groups (Lipson et al., 2017), (besides one such case from today’s Romania (González-Fortes et al., 2017)) despite the known HG presence in the region at the beginning of the Neolithic (Gamba et al., 2014; Mathieson et al., 2018). In contrast, ancient populations from other parts of Europe, such as Scandinavia (Günther et al., 2018; Malmström et al., 2019; Mittnik et al., 2018; Skoglund et al., 2014), today’s Poland (Fernandes et al., 2018) or Iberia (Olalde et al., 2019) show much higher and much later introgression of HG ancestry. Later on, at the beginning of the 3rd millennium BCE, the appearance of steppe related ancestry shaped the regional genetic landscape extensively, founding the modern day European genetic makeup (Allentoft et al., 2015; Brandt et al., 2015; Gamba et al., 2014; Haak et al., 2015; Lazaridis et al., 2014; Linderholm et al., 2020; Mathieson et al., 2018; Olalde et al., 2018).

Besides monitoring population events, archaeogenetics opens a new window to study health qualities of ancient populations that may lead to a better understanding of the background of recent genetic related diseases. Studies aimed at uncovering variants under selective pressure in *Homo sapiens* populations, such as Lactase persistence (LCT) or Human Leukocyte Antigen (HLA) genes and pigmentation markers are beginning to thrive (Childebayeva et al., 2022; Evershed et al., 2022; Lazaridis et al., 2022; Mathieson et al., 2015). However, variants for rare genetic diseases or aneuploidies are sparsely checked on ancient datasets, except for a few cases, such as the study of the Suontaka grave (Moilanen et al., 2022).

Our study aimed to make a transect analysis on a single site from understudied archaeological assemblages. This was combined with population genetic analyses, isotope analyses, and phenotype (pigmentation) and clinical variant analyses. Moreover we present a series of bioinformatic tools for biological relatedness, ploidy and variant analyses implemented in a new bioinformatic package for ancient DNA analysis. We analysed archaeological finds from the Balatonkeresztúr-Réti-dűlő site in Western Hungary (Transdanubia), where - among others - Bronze Age features and human remains were found during roadwork in 2003. Three Bronze Age archaeological horizons were distinguished based on ^14^C dates: the Somogyvár-Vinkovci culture (∼2500-2200 BCE, n=1), Kisapostag culture (∼2200–1900 BCE, n=11) and the Encrusted pottery culture (∼1900–1450 BCE, n=8) that are referred to as Bk-I, II and III phases in this study, respectively (Table 1, Supplementary Information section 1). All three cultural horizons have only a limited number of inhumation remains: this study presents the first validated Somogyvár-Vinkovci culture associated individual from Hungary, while the Kisapostag and Encrusted pottery cultures have been mainly characterised by cremation burials so far. The archaeological origin of the Kisapostag culture is enigmatic and multiple theories have been proposed to explain its possible connections. One theory is that the pottery decoration technique observed in Kisapostag originated either from Corded Ware in the Middle Dnieper region (Ukraine) or epi-Corded Ware groups (from the northern Carpathians), *e.g.* Chłopice-Veselé (Slovakia). The latter option is also supported by inhumation practises and burial positions (Bándi, 1984; Bóna, 1975, 1961; Hajdu, 2010; Kiss, 2020; Szabó, 2010). However, local development of communities with eastern (Makó–Kosihy–Čaka) or southern (Somogyvár–Vinkovci) origins, as well as western and southwestern connections (with the Litzenkeramik or Guntramsdorf-Drassburg group in eastern Austria, Slovenia, western Croatia) have also been raised in the archaeological literature (Bóna, 1992; Črešnar, 2010; Kiss, 2015). Additionally, Bell Beaker influence has been proposed based on the craniometry data (so-called *Glockenbecher* or brachycranic skull type (K. Zoffmann, 2008, 2007; Mozsolics, 1941)). Nevertheless, the archaeological connection between Kisapostag and Encrusted Pottery cultures are strong and well established (Kiss, 2012). In order to provide an additional data point to our analysis of population ancestry of the region, a Late Copper Age individual from the Baden culture (∼3600-2800 BCE), excavated ∼30 km away from Balatonkeresztúr was added to the dataset. Our data highlight not only detailed population events in a microregion, but also reveal hidden processes that formed the genetic landscape of East-Central Europe at the beginning of the Bronze Age.

**Table 1:** Summary of the investigated samples. MtDNA and ChrY denote mitochondrial haplogroup and Y chromosome haplogroup, respectively. In column “Biological relatedness” 1^st^ and 2^nd^ mean the degree of relations. For the feature, grave ID and details on newly reported ^14^C dates see the Supplementary Table 1.

| Group | ID | Grave group | cal BCE date (95.4% CI) | Age | Sex | MtDNA | ChrY | Biological relatedness |
| --- | --- | --- | --- | --- | --- | --- | --- | --- |
| Baden | BAD002 |  | 3530-3370 | 8-9 | M | K1a4a1 | I-M170 |  |
| Bk-I:<br>Somogyvár -<br>Vinkovci culture | S9 |  | 2560-2290 | 35-40 | M | K1a3a | R1a-V2670 |  |
| Bk-II:<br>Kisapostag or<br>Early Encrusted<br>Pottery culture | S1 | A | 2120-1880 | 40+ | M | V | I2a-L1229 | 2<sup>nd</sup> to S2 |
|  | S2 | A | 2120-1880 | 30-35 | M | U5a2b1a | I2a-M436 | 2<sup>nd</sup> to S1 |
|  | S4 | B |  | 17-19 | M | H10a1 | I2a-L1229 | 1<sup>st</sup> to S8 |
|  | S5 | A |  | 16-18 | M | T1a4 | I2a-L1229 | 1<sup>st</sup> to S6 & S11 |
|  | S6 | A | 2030-1770 | 17-18 | M | T1a4 | I2a-L1229 | 1<sup>st</sup> to S5 & S11 |
|  | S7 | A | 2120-1880 | 35-50 | F | V |  |  |
|  | S8 | B |  | 30-40 | M | T2b | I2a-L1229 | 1<sup>st</sup> to S4 |
|  | S10 |  | 2140-1940 | 7-8 | M | K1a4a1g | I2a-L1229 |  |
|  | S11 | B | 2200-1980 | 34-43 | M | T2b | I2a-L1229 | 1<sup>st</sup> to S5 & S6 |
|  | S13 | B | 2120-1890 | 35-45 | F | J2b1 |  |  |
|  | S45 |  | 2200-1980 | 45-55 | M | U5a1g | I2a-L1229 |  |
| Bk-III:<br>Transdanubian<br>Encrusted Pottery<br>culture | S14 | Mass<br>Grave<br>B-938 |  | 7-8 | F | H10a1 |  |  |
|  | S15 |  |  | 21-23 | M | U4b1b1 | I2a-L1229 | 2<sup>nd</sup> to S17 |
|  | S16 |  | 1890-1640 | 35-44 | M | T2g2 | I2a-L1229 |  |
|  | S17 |  | 1870-1540 | 26-35 | M | U5b1b1+@16192 | I2a-L1229 | 1<sup>st</sup> to S19; 2<sup>nd</sup> to S15 |
|  | S18 |  |  | 3-4 | M | U4a2 | R1b-Z2103 |  |
|  | S19 |  |  | 9-10 | M | T2b | I2a-L1229 | 1st to S17 |
|  | S20 |  |  | 1.5-2 | M | K1a+195 | R1b-Z2103 | 1st to S21 |
|  | S21 |  |  | 1.5-2 | F | K1a+195 |  | 1st to S20 |

## Results

We shotgun sequenced genomes of 21 individuals yielding between 0.008x and 2.1x average genomic coverage. We also sequenced reads of a capture set consisting 3000 nuclear SNPs (single nucleotide polymorphisms, see Methods), and whole mitochondrial DNAs (mtDNAs) of all individuals. The shotgun and the capture sequenced samples ultimately resulted in an average ∼144k SNPs/individual using the 1240k SNP panel for genotype calling (Mathieson et al., 2015), see Materials and Methods and Supplementary Tables 4 and 7. We utilised STR (Short Tandem Repeat) analysis of the Y chromosome to ascertain direct paternal connections (Supplementary Table 3). The Y-STR profiles resulted in comparable haplogroup predictions with the NGS Y-SNP data, and were authenticated via repeated reactions (see Methods). Furthermore, by using all known biological and archaeological details, we reconstructed the face of individual S13 (from phase Bk-II), see Supplementary Information section 4. The bioarchaeological analyses included ^14^C dating and ^87^Sr/^86^Sr isotope analyses, the latter is routinely used to trace individual mobility (Alt et al., 2014).

### Archaeological and anthropological evaluation of samples

We included only one Infans II (late childhood) individual (BAD002) from the site of Balatonlelle belonging to the early phase of the Baden culture (3530-3370 cal. BCE). A majority of the samples came from the site of Balatonkeresztúr Réti-dűlő. We sampled and sequenced 20 individuals, labelled S1 to S45, skipping numbers that did not belong to the Bronze Age horizon or weren’t suitable for genetic testing. One male individual (S9) that was associated with Bk-I by ^14^C data, has a very long (ultradolichocran) skull type, which differentiates him from most individuals in Bk-II and Bk-III, who have very short (brachycranic) skull types (Köhler, 2014) (Table 1). The male dominance (∼78%) in Bk-II and Bk-III suggests distinctive funeral treatment for males and females. Bk-II phase is represented by an Infans II (7-8 year old male), 3 juveniles (16-19 year old males) and 7 adults (30+ year old). They are spatially distributed into grave groups of A and B with two additional, separate inhumations (Table 1, Supplementary Information Fig. S.1.2.1). Most of the burials contained no grave goods except for small copper jewellery in S10 and S13, and some shell fragments in S45. Radiocarbon dates place these inhumations to ∼2200-1770 cal BCE (95.4% CI), however, with Bayesian analysis using the OxCal software (Bronk Ramsey, 2009) the timespan of the Bk-II burials can be reduced to ∼2050-1940 cal BCE with a 84.4% CI, whereas only two graves (individuals S10 and S11) were possibly slightly earlier (Supplementary Information section 1.8). The lack of infants at Bk-II is similar to other archaeological sites as this phenomenon is common in most periods. It can be associated with different skeletal taphonomy for infants or differential burial treatment as compared to the burial of adults (Alt et al., 2014). The reason for the absence of young adults (∼20-30 year olds) is unknown. Bk-III is represented by a single mass grave (∼1870-1620 cal BCE) with skeletal remains of 8 people of various ages. This is an unusual burial treatment as most bodies were cremated or buried as single inhumations. For detailed descriptions of the sites and burials, see Supplementary Information section 1.

### Uniparental genetics and relatedness analyses

Analysis of uniparental (maternal and paternal) lineages provide rough estimates of genetic composition and are essential to assess biological relatedness and social structure for the studied population. Additionally, we performed phylogenetic analysis using MrBayes software (Ronquist and Huelsenbeck, 2003) on mitochondrial DNA to see the phylogeographic affinities of the studied individuals. Accordingly, the valid phylogenetic trees show diverse connections in both Bk-II and Bk-III to various modern and ancient populations, but the overlap between the two horizons are limited. For detailed mtDNA analyses, see Supplementary Information section 2.1.

Contrary to the diverse mtDNA makeup, most male individuals in Bk-II and Bk-III belong to the Y chromosome haplogroup I2a-L1229, except for two individuals that belong to haplogroup R1b-Z2103 (Table 1). Similar phylogeographic analysis to the mitochondrial DNA can be performed on the paternal lineages as well using STR markers. Network analysis (Supplementary Information section 2.2) narrowed down regional Y-chromosomal affinities to the North European plain and indicated continuity between Bk-II and Bk-III. Uniparental makeup shows a patrilocal social structure that is similar to previously reported Bronze Age findings (Mittnik et al., 2019; Schroeder et al., 2019; Žegarac et al., 2021). Results are highly similar to previous observations on Encrusted Pottery culture’s population at the Jagodnjak site, Croatia (Freilich et al., 2021). Inference on biological relatedness was based on READ (Monroy Kuhn et al., 2018) and a newly developed method called Modified Pairwise Mismatch Rate (*MPMR*, Supplementary Information section 2.3). The relatedness network (Fig.1.b) of Bk-II approximately follows the distribution of individuals in A-B grave groups (Fig.1.a), which were likely established along family relationships and chronology. Individuals buried in the Bk-III mass grave only show a few blood relations (Fig.1.c): a half-brother or uncle-nephew, a father-son and a dizygotic twin, to our knowledge the latter is the oldest detection of such relatedness. None of the distant inhumations (S10, S45) show biological relationship to any other individuals up to second degree. For further details, see Supplementary Information section 2 and Supplementary Tables 1-3.

**Fig. 1.**
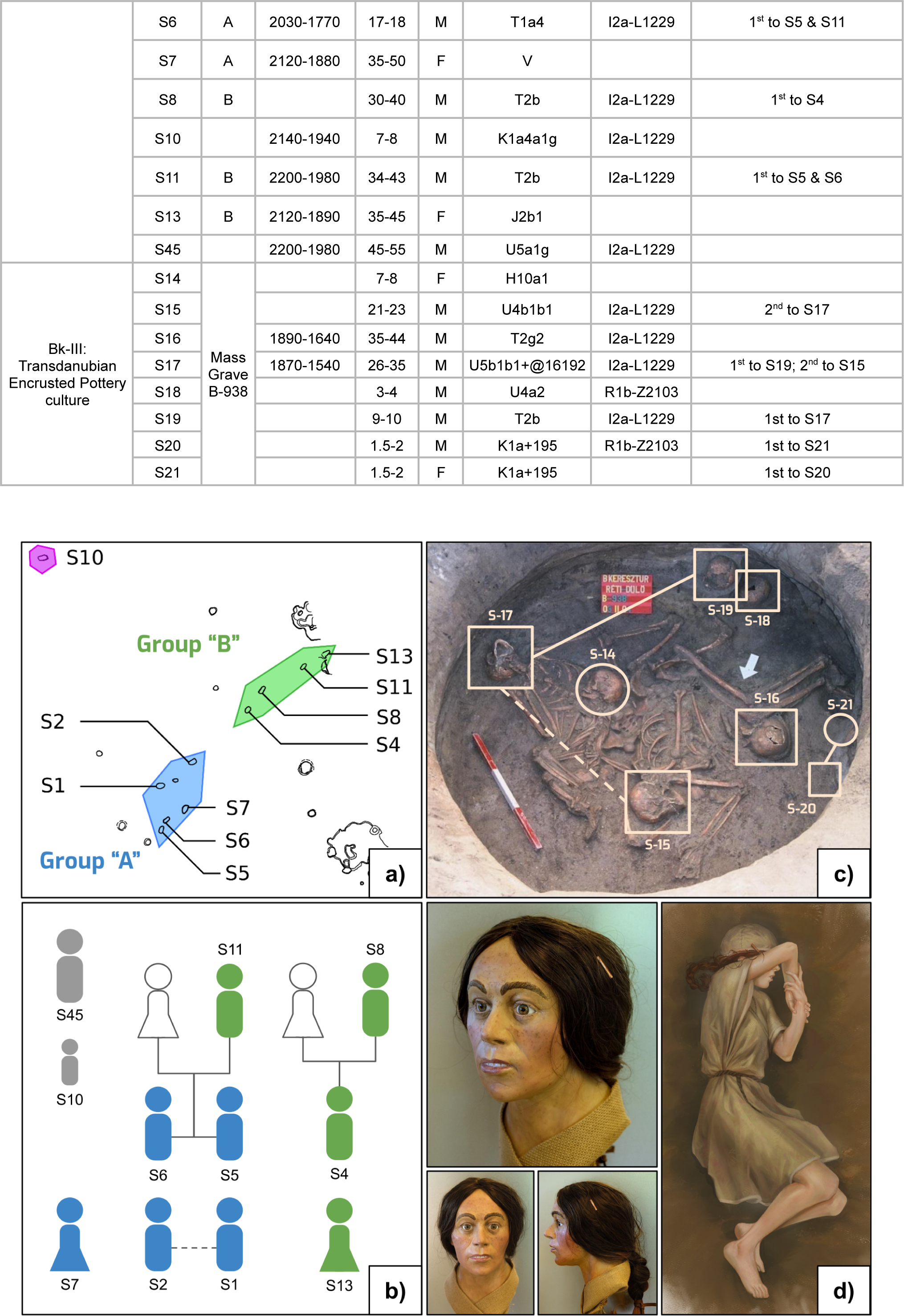
Biological relatedness of Bk-II and Bk-III, reconstruction of individual S13. a) Distribution of graves of Bk-II: individuals S1, S2, S5, S6 and S7 belong to grave group ‘A’ highlighted in blue, whereas individuals S4, S8, S11 and S13 belong to grave group ‘B’ highlighted in green. Individuals S10 and S45 (not shown) placed separately from grave groups. b) Relatedness network of Bk-II individuals, where colours denote the corresponding grave groups in figure ‘a’. The dashed line between individuals S1 and S2 represent an undirected second degree relationship. c) Relatedness of Bk-III projected onto the photo of the mass grave (obj. B-938). Squares denote males, circles females, solid lines first degree and dashed lines second degree relationships. d) Forensic facial and burial reconstruction of individual S13.

### Genetic disorders and pigmentation

Investigating genetic disorders in archaeological datasets can be useful to improve our knowledge of the history of health and medicine, and also highlights the overall genetic health of past populations. Genetic disorders, if accompanied with severe phenotypic anomalies, could also explain unusual burial practices, for example as it was described in cases of dwarfism in the Byzantine era (Slon et al., 2013), or in the case of the Suontaka burial (Moilanen et al., 2022). Therefore, we analysed the ploidy of the autosomes not only for genetic sex determination, but to recover possible aneuploidies resulting in serious health related traits, such as Turner or Down syndrome (O’Connor, 2008). For inferring pigmentation of the studied individuals, we used the HIrisPlex (Chaitanya et al., 2018) system supplemented with further variants obtained from SNPedia database (Cariaso and Lennon, 2012). Finally, we created an in-house disease panel described further below and in Supplementary Information section 3.

#### Aneuploidies

Abnormal numbers of chromosomes results in a few well known diseases which we tested for thoroughly. We developed a new method called Z-score Adjusted Coverage (*ZAC*) for ploidy estimates by using a set of reference genomes. This can estimate ploidy for samples as low as 0.008x average genomic coverage, enabling genetic sex determination and aneuploidy assessment for all new samples (Supplementary Information section 3.1). As a result, we found one individual, S10 - the only child burial in Bk-II - with an XYY gonosomal genotype, described as Jacob’s syndrome. This syndrome is relatively frequent (∼0.1%) in today’s populations. In most of the cases it remains silent but occasionally comes with a wide scale of symptoms, mainly behavioural disorders (Bardsley et al., 2013).

#### Mitochondrial DNA diseases

We examined the clinical significances of the polymorphisms that can be found in mtDNA by using the *mitopathotool* software on the *AmtDB* database (Ehler et al., 2019), and found that individual S1 (40+ years old male from Bk-II) had one of the defining mutations (T14484C, 48x coverage) of Leber’s hereditary optic neuropathy (LHON) causing complete vision loss in ∼50% of males between the ages of 20 to 40, and in rare circumstances by other neuropathies (Tońska et al., 2010).

#### Nuclear variants with clinical significance

We also examined the nuclear genomes to find regions with clinical significance. Since a complete panel for determining disease susceptibility only exists in commercial DNA kits with non-available descriptions similar to the 1240k panel, we created a new SNP calling panel focused on various conditions including amyotrophic lateral sclerosis, Alzheimer disease, autism, Crohn’s disease, diabetes, lactose intolerance, mental disorders, Parkinson disease, schizophrenia and ulcerative colitis. For this study we used a ∼5k set of clinically significant SNPs, which were marked as “pathogenic” or “likely pathogenic” in the ClinVar database (Landrum et al., 2018), by ignoring deletion, duplication and copy number variants, as well as SNPs with questionable (signed as “reported”, “conflicting reports”, *etc.*) contributions to diseases. The exact method of calling variants can be found in Supplementary Information section 3.2.2. Both the tool and the ∼5k set are built in the PAPline package. We also created a bioinformatic tool that rolls up variant information from input data, which is available in the PAPline package (see the *PAPline* section for details). After running the panel, we excluded low coverage transitional variants from the final evaluation due to the possible presence of DNA damage. We only made exceptions when skeletal features supported the presence of the low quality variant or when more than one sample possessed the same allele. Nevertheless, we listed all alternate variant hits in Supplementary Table 6. We are aware that low coverage data is not sufficient for firm conclusions, however, the aim was more of a technical description of such analyses. Here we summarise only a few mentionable results of the run, but for the detailed discussion see Supplementary section 3.2.

Lig4 syndrome is a transitional mutation (rs104894421) induced disease with skeletal abnormalities (Altmann and Gennery, 2016), for which individuals S15 and JAG93 from the Jagodnjak site of the Encrusted pottery culture (Freilich et al., 2021) both provided a single read hit. We excluded the Jagodnjak group from our analyses for the lack of UDG treatment, meaning that both hits could be false positives; however, individual S15 possesses the distinguishing skeletal features of this disease, increasing the possibility of the actual presence of this allele in the Encrusted pottery population. Another ambiguous, but possible hit is rs121434442 in individual S6. This SNP is the causative factor for hereditary spastic paraplegia (Shribman et al., 2019) a disease mostly recognised by muscle stiffness in the lower limbs causing movement restrictions. Individual S11, father of S6, shows signs of a limb condition that may be linked to this disease. Finally, autism 15 susceptibility signature transversional variant (rs7794745) (Koeda et al., 2015) was present in individuals S6 and S45. Severe bruxism on the upper front teeth of S45 (Supplementary figure S.3.2.2.4) suggests compulsive behaviour that occurs frequently among people with autism spectrum disorder (Koeda et al., 2015; Muthu and Prathibha, 2008). While this condition itself could also be linked to some profession related abrasion, the physical features combined with genetic data and the distinguished burial treatment (Supplementary figure S.1.5.11) support the possibility of the actual onset of symptoms.

#### Pigmentation

According to our results based on a final set of 58 SNPs, the pigmentation patterns are highly different between horizons, as Bk-I mostly possesses variants for light pigmentation, blue eyes and blonde hair, while Bk-II is more similar to populations of Neolithic Europe of darker colouration (Childebayeva et al., 2022; Lazaridis et al., 2022) (Fig.1.c), although some variants for lighter pigmentation exist within this group too. Members of Bk-III on the other hand show a wide range from dark to light pigmentation tones and even the presence of variants for red hair (Supplementary Table 5, Supplementary Information section 3.2.1).

### Whole genome composition and genetic ancestry

#### Balatonkeresztúr site samples

To get a general overview of the autosomal composition of the individuals, we performed Principal Component Analysis (PCA) with *smartpca* software (Patterson et al., 2017) based on 590k nuclear SNPs (Reich, 2021) and ADMIXTURE (Alexander et al., 2009) analyses based on the 1240k SNP set (Reich, 2021). According to PCA (Fig.2.a) Bk-I is clearly separated from Bk-II and Bk-III, where Bk-II has a strong shift towards European HGs overlapping with only a fraction of known ancient samples (Reich, 2021) and Bk-III. *Admixture* analyses (Fig.2.b) for assessing genetic components revealed ∼17% HG, ∼40% EEF, and ∼43% steppe ancestry for Bk-I, similar to average Bronze Age Europeans (Allentoft et al., 2015; Brandt et al., 2015; Gamba et al., 2014; Haak et al., 2015; Olalde et al., 2018) (Supplementary Table 9, 12-16; Supplementary Information sections 5.2, 5.5.2, and 5.6). According to *qpAdm* (Patterson et al., 2012), Bk-I is most likely the ∼1:2 mixture of a Vučedol culture associated individual (Croatia_EBA_Vucedol_3, ∼38±4%), and a mostly steppe characteristic source. This steppe source can be best modelled as a Srubnaya/Alakul culture related population (Russia_Srubnaya_Alakul.SG, ∼62±4%), in line with archaeological observations (Kulcsár, 2009). However, this high proportion of steppe ancestry is likely derived from a previously unsampled group in Eastern Europe, maybe in the vicinity of the Baltics (for details, see Supplementary Information section 5.6.2.1). Bk-II comprises a unique makeup of ∼42% HG, ∼41% EEF, and ∼17% steppe ancestries. *qpAdm* analysis revealed most plausible sources as a Sweden_FBC (Funnelbeaker culture) related population and Ukraine_EBA with almost equal contributions (Supplementary Information section 5.6.2.2), however, both populations are likely only an approximation for the actual ancestry of Bk-II, which we discuss further below. Bk-III shows a slight shift in ancestry composition from Bk-II with ∼29% HG, ∼46% EEF, and ∼25% steppe ancestries. *qpAdm* analyses uncovered that the main ancestry component for Bk-III is Bk-II (∼60±8%), while “dilution” of Bk-II to Bk-III is mostly driven by contact with various local populations, genetically best represented by later Transdanubian Hungary_LBA or Serbia_Mokrin_EBA_Maros (Maros culture) groups (Supplementary Information section 5.6.2.3, Supplementary Table 15).

**Fig. 2.**
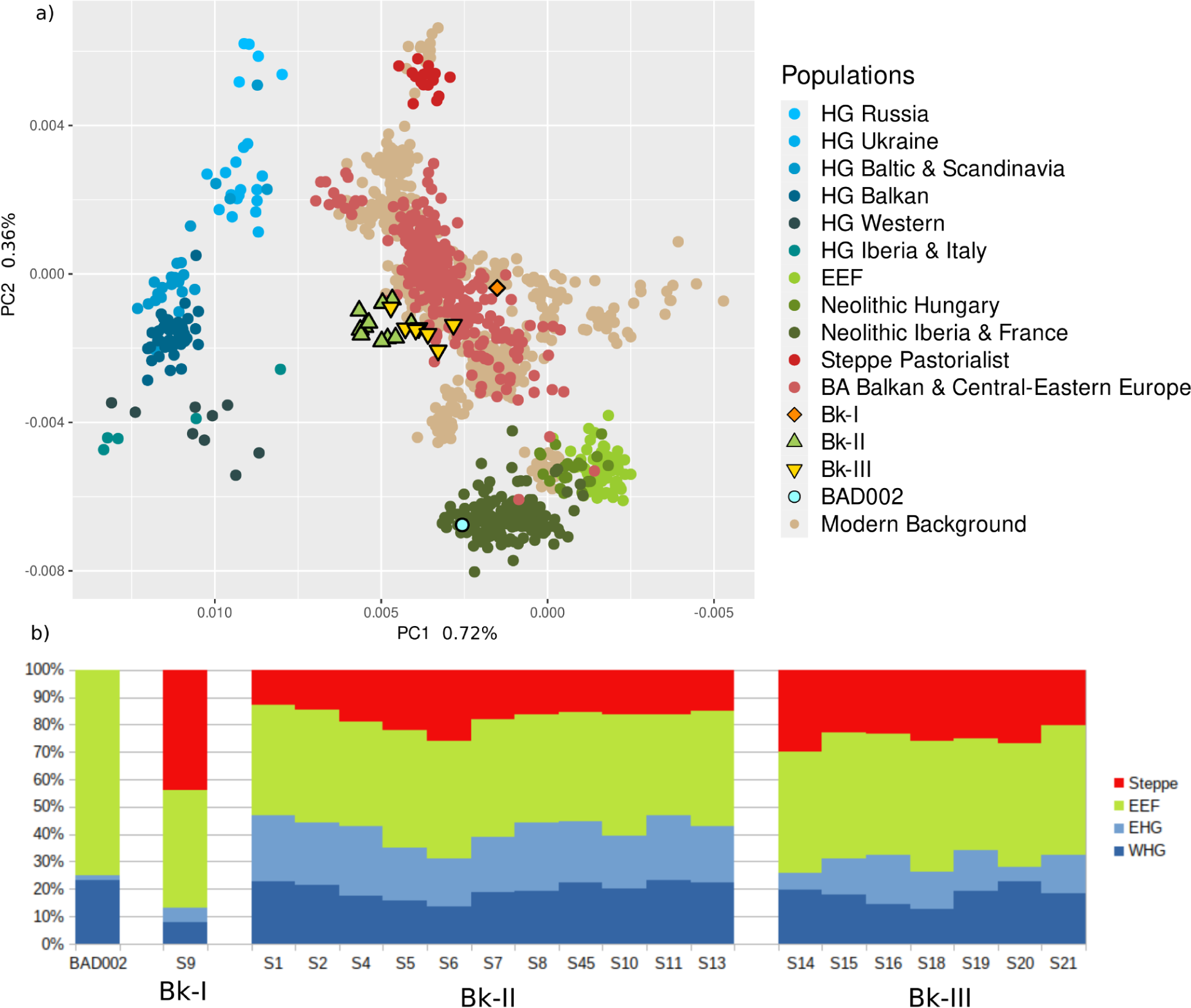
Basic genetic composition of the investigated samples. a) Principal Component Analysis based on 590k SNPs calculated by the smartpca software (Patterson et al., 2017), where Bk-II (marked with green triangles) clearly separated from all other ancient Central-Eastern European populations. b) The admixture proportions of the BAD002, Bk-I, Bk-II, and Bk-III samples, where the percentage of steppe ancestry is shown with red, early European farmer (EEF) with light green, Eastern hunter-gatherer (EHG) with light blue, and Western hunter-gatherer (WHG) with dark blue colour (supervised Admixture analyses).

#### Genetic outliers from previous studies and the origin of HG ancestry in Bk-II

We aimed to investigate further the exact composition of HG ancestry in Bk-II. *qpAdm* analysis of the basic composition resulted in EEF ∼40±2%, EHG ∼39±3%, WHG (Western HG) ∼13±2.7%, Caucasus HG ∼8±2%, *p*=0.0917 in par with *Admixture* analysis, pointing towards a rather EHG characteristic composition. Next, we performed an *f4* test in the form of *f4*(test HG, Serbia_IronGates_Mesolithic, Bk-II, Mbuti.DG) (Patterson et al., 2012), to see which HG population relates the best with the Bk-II samples. The aim of this analysis was to detect different HG ancestry contributions besides Iron Gates HG, which has knowingly contributed to Neolithic European populations in the study region and witnessed an intermediate composition between the WHG and EHG (Mathieson et al., 2018). Contrary to the *Admixture* and *qpAdm* results, this test revealed that Bk-II individuals have excess in HG ancestry mainly from WHG groups or other mixed characteristic HGs (Croatia_mesolithic, Poland_BKG_o2.SG (Brześć Kujawski Group outlier) or KO1 (Körös culture outlier HG)), but only marginal relations with the EHG (Lithuania_Mesolithic) populations (for detailed results, see Supplementary Information section 5.3). Surprisingly, none of these HG populations with mixed characteristics (and neither Iron Gates) have enough EHG component to explain the ancestry of the Bk-II samples. The *f4* test also revealed that the Bk-II and Bk-III populations differed significantly from other EHG characteristic populations, such as HG-s from today’s Russia, Ukraine, Baltics (younger phase) or Scandinavia, although we can see some weak connections to older (down to sixth millennium BCE) Lithuanian HG-s. These results may reflect the population turnover in the sixth millennium BCE in the Baltics (Mittnik et al., 2018), suggesting that this EHG ancestry is related to the Lithuania_Mesolithic. On the other hand, *qpAdm* provides negative weights for this component when we try to model Bk-II as a combination of WHG (Loschbour_WHG), EHG (Lithuania_Mesolithic), EEF (Turkey_N) and Yamnaya (Russia_EBA_Samara_Yamnaya), suggesting that Lithuania_Mesolithic is not a good proxy for the actual EHG component.

To infer the timing of HG admixture, we used the *DATES* (Chintalapati et al., 2022) analysis. This test revealed that the HG ancestry in Bk-II resulted from three independent admixture events: one from Iron Gates HG at the beginning of the Neolithic (similar to other populations at that time), one from a WHG characteristic source around the turn of the fourth and third millennium BCE, and an EHG characteristic source around the second half of the third millennium BCE (for details, see Supplementary Information section 5.4). Summarising these results, we conclude that the EHG characteristic source of the Bk-II individuals does not exist in the current database.

We were interested in whether other populations carried this peculiar HG ancestry, to see which region it might originate from. To achieve this, we did a literature search to select individuals with high levels of HG ancestry, who were genetic outliers in their cultural or geographical or temporal context, in order to assess whether they are related to the Bk-II group. Selection was based on previous observations and HG ancestry differences within groups using the results of the *Admixture* analysis (Supplementary Information section 5.2.2). Then, to reveal similar patterns of HG ancestry, we ran *f3* statistics in the form of *f3*(test HG, test population, Mbuti.DG) on all of the groups (obtained from AADR (Reich, 2021) database, listed in Supplementary Table 8). Subsequent euclidean distance based clustering of *f3* values revealed a number of outliers and even whole populations belonging to the same subcluster as Bk-II (Fig.3, Supplementary Information section 5.5). Accordingly, the earliest signs of such HG ancestry appeared among various Neolithic groups from Western Europe (in line with characteristically high WHG ancestry among Megalithic, Globular Amphora or Funnelbeaker cultures’ population) and from Eastern Europe (Bulgaria and Ukraine). Individuals with this ancestry predating Bk-II by only a few generations appeared in Czechia, Northern Hungary, Eastern Germany and Western Poland, indicating that the Kisapostag associated population came to Transdanubia via a Northern route, in line with the observations of (Freilich et al., 2021).

**Fig. 3.**
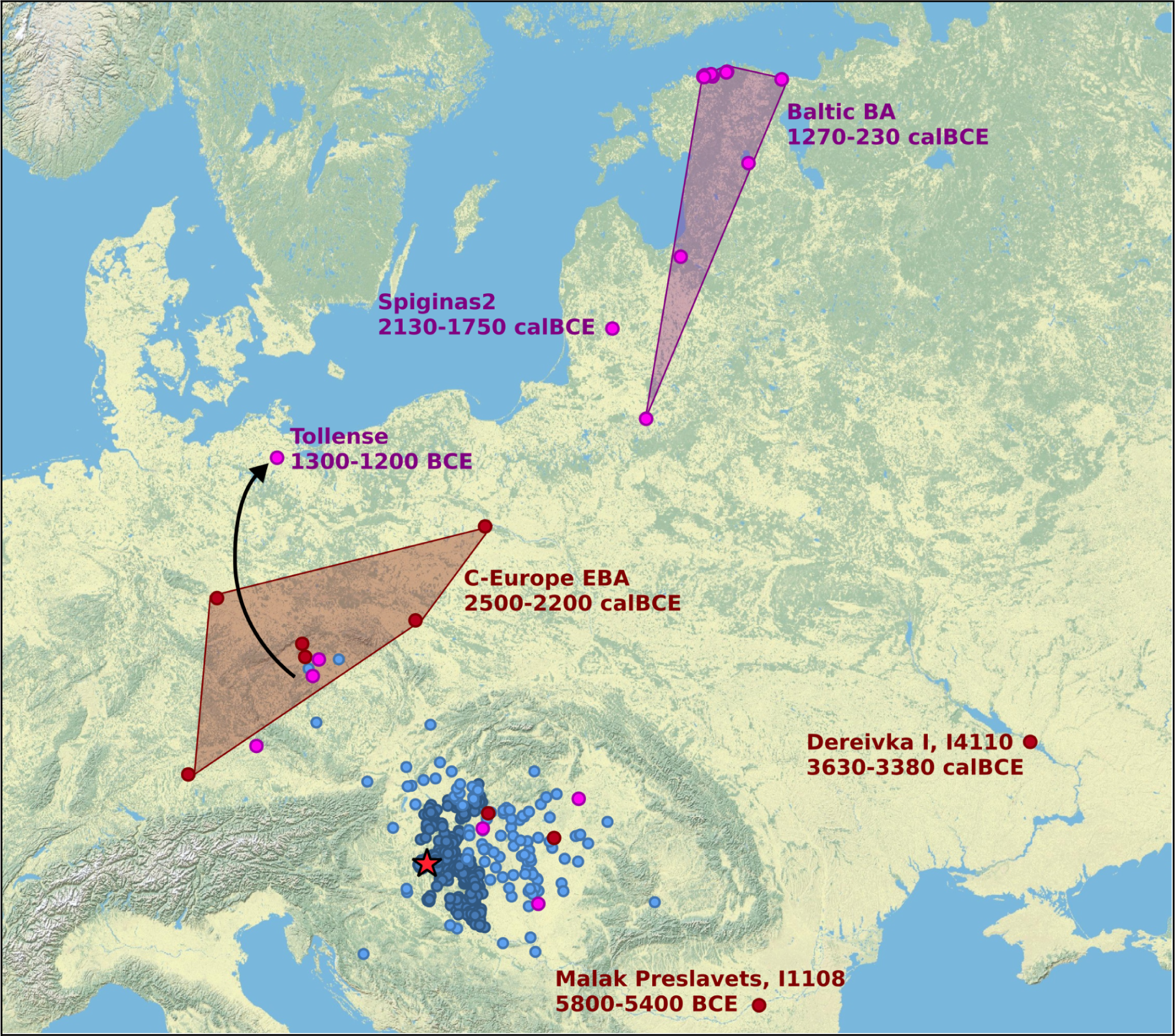
Map of East-Central Europe with sites and genetic parallels of Kisapostag/Encrusted pottery culture. The map shows the site of Balatonkeresztúr (red star), Kisapostag and Encrusted pottery culture archaeological sites (dark blue circles), their archaeological connections based on pottery and metal finds (light blue circles) after Kiss^36^. Red and purple circles represent individuals that are connected to Bk-II individuals by HG ancestry. Also, red circles are preceding, while pinks are succeeding or contemporaneous to the Bk-II horizon. For the detailed list see Supplementary Information *Table 5*.5.1 and for method description, see Supplementary Information *section 5.5*.

Many contemporaneous populations to Bk-II and Bk-III from the British Isles to today’s Poland, down to today’s Serbia have outliers with a Bk-II-like genomic composition, mostly overlapping with known Kisapostag and Encrusted pottery contact regions (Fig. 3). Interestingly, at the end of the second millennium BCE many Baltic groups appear to be highly similar to Bk-II, indicating long term success of this ancestry outside the Carpathian Basin. Notably, in the vicinity of Prague many pre- and post-Bk-II outliers appear along with the archaeological presence of the Kisapostag culture, including the Tollense group, which also originates from the region of Bohemia according to isotopic evidence (Price et al., 2019), suggesting a local reservoir of the population. While the appearance of Bk-II ancestry in the Baltics could be connected to this reservoir, especially in the light of the mobility of Tollense group, the ^14^C date of Lithuania_LN_o around 2000 BCE suggest that the population was likely prevalent in nearby unsampled regions of Eastern Europe.

Taking into consideration all of the genetic parallels, their dates and geographic locations, one plausible scenario is that the EHG characteristic core of Bk-II (which ultimately could be best modelled as Ukraine_EBA by composition) moved northward from the region of today’s Eastern Romania, Moldavia or Western Ukraine, subsequently mixed with Funnel Beaker culture (FBC) or Globular Amphora culture (GAC) related populations and then split into two groups: one taking a route to Transdanubia, and the other moving further North. These results are highly in par with (Mittnik et al., 2018), who suggested population replacement at the end of the second millennium BCE in the Baltic region from a nearby, unsampled region by a population of considerably higher steppe, EEF and WHG ancestry than the prevailing ones; however, further data are needed from Eastern Europe to affirm this hypothesis.

#### Isotope analyses

We sampled molars from the burials and measured the ratio of ^87^Sr/^86^Sr isotopes to evaluate whether individuals were born in the area of their burial. The results (Fig. 4) show that burials from Balatonkeresztúr have strontium isotope values that match local plants and water, which suggests that none of the studied individuals are first generation occupants. It is, however, interesting how the M3 molar values for individuals S15, S16 and S17 (all from the mass grave) differ from the others. While these values are not out of the estimated local isotope range for the area, they could indicate movement within the region during adolescence when those tissues are forming. This movement could have occurred at the same time for these individuals, as the stronger the divergence from the majority, the younger the individual was at the time of death. It is particularly interesting how individual S15 shows the highest divergence from the others, as this individual had severe complications for walking due to hip dysplasia (see also Supplementary Information section 3.2.2). Moreover, since M3 values show divergence, but their M1 do not, these individuals likely grew up in the vicinity of the site, spent some years away from it, and then returned to the same place where they died and were buried.

**Fig. 4.**
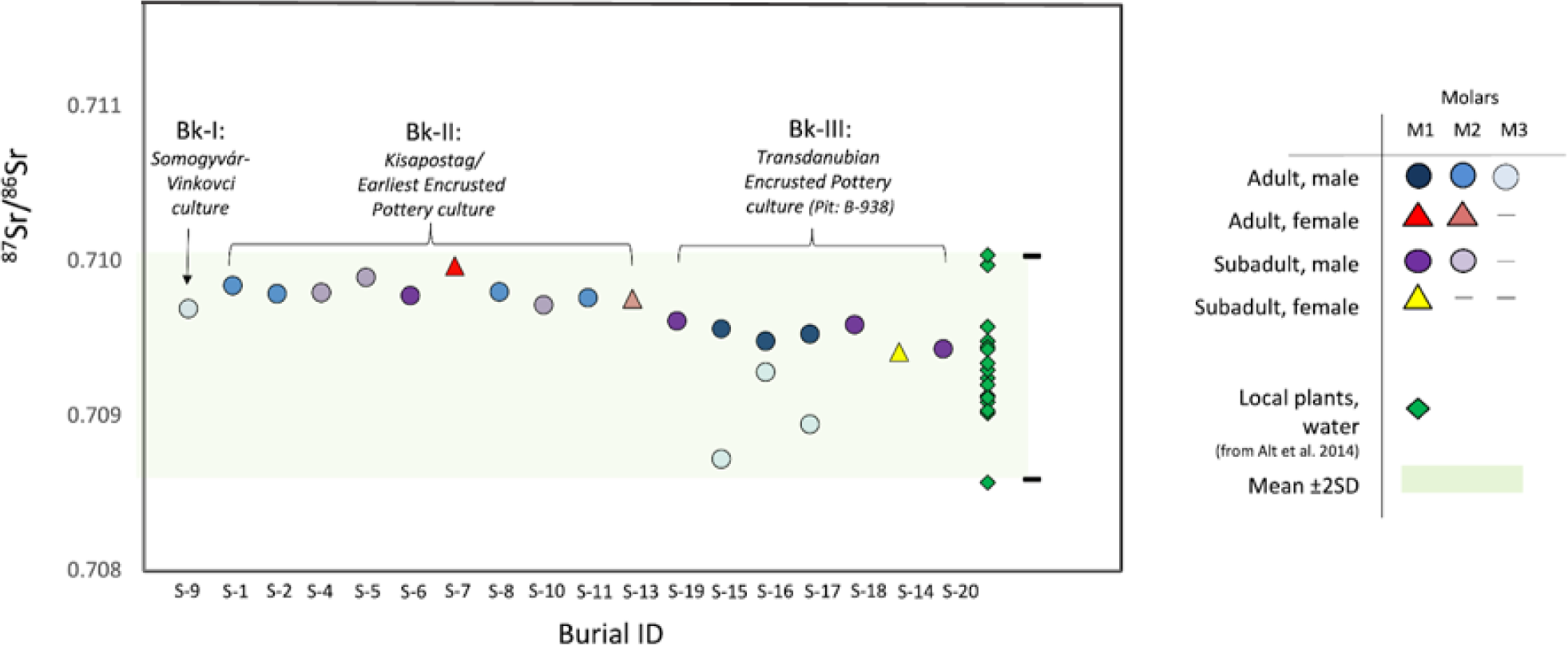
^87^Sr/^86^Sr isotope data from the Balatonkeresztúr site. Samples were taken from dental enamel (first, second and third molars) to evaluate whether individuals were born in the area, or grew up in a geologically distinct region. All of the samples are consistent with previously published plant and water ^87^Sr/^86^Sr ratio (green diamonds) data collected from the southern portion of Lake Balaton (Alt et al., 2014). For further data see Supplementary Information *section 1.9*.

#### A Late Copper Age outlier individual from Balatonlelle site

We included in this study a Late Copper Age individual, BAD002, from the Balatonlelle site because of his high HG genomic ancestry component. The mitogenome of BAD002 (K1a4a1) shows affinity to the Iberian Bell Beaker culture associated individuals (Supplementary Information Fig. S.2.1.1). His Y chromosomal haplogroup belongs to I-M170. Compared to known Neolithic and Copper Age populations in the Carpathian Basin (Lipson et al., 2017), BAD002 has a higher HG component (∼34%), and he also lacked steppe related ancestry. Therefore, on the genomic PCA, BAD002 relates with Iberian and French Neolithic individuals. According to our ancestry estimates, France_MontAime_MLN.SG describes best the BAD002 individual but other Western European sources, such as Spain_EN are also plausible (for details, see Supplementary Information section 5.6.1). The pigmentation pattern of BAD002 shows resemblance to average Neolithic Europeans. The foreign cultural traits of the boy’s jewellery is in line with his outlier genetic composition in the study region (Bondár and Szécsényi-Nagy, 2020), therefore we conclude that this individual exemplifies large-scale migration in the Copper Age, providing research questions for future studies. Finally, further tests (outgroup *f3*-statistics and *qpAdm*) excluded the contribution of BAD002 to Bk-II (Supplementary Information section 5).

### PAPline

We introduce a newly developed, freely available bioinformatic package, named *PAPline* (Performing Archaeogenetic Pipeline), written in *linux bash, R,* and *python v3.8.10* programming languages. One can use this package primarily to analyse next generation sequencing data of archaeogenomic samples, supplemented by tools, including ploidy test, MPMR relatedness analysis and clinical variant test. The standalone tools and the core workflow of *PAPline* is available at https://github.com/ArchGenIn/papline. In the future *PAPline* is aimed to be compared to the EAGER (Peltzer et al., 2016) and the Paleomix (Schubert et al., 2014) pipelines, for a detailed description visit the github page.

## Discussion

The Carpathian Basin was inhabited by the Baden population at the end of the Copper Age. Their genetic composition was represented by an EEF and – compared to the previous Neolithic populations of the region (Lipson et al., 2017) – a slightly increased HG genetic component. Here, we demonstrated that in the early phase of this culture, a Western European group appeared in Transdanubia, diversifying our previous knowledge about the region’s Late Copper Age.

The Carpathian Basin experienced the influx of steppe-related genetic ancestry during the Late Copper Age (Mathieson et al., 2018; Olalde et al., 2018). This transformation was already detectable in the Bronze Age genetic picture of the Balatonkeresztúr-Réti-dűlő, where we could examine multiple populations. The earliest Bronze Age horizon Bk-I (representative of the Somogyvár-Vinkovci culture) is best described by the mixture of local (Vučedol) and a high steppe ancestry population from Eastern Europe, however, since this group has data for only one individual, future studies on this archaeological culture and region may diversify the composition of the prevailing population. Bk-I was replaced by the Kisapostag culture associated group of Bk-II, likely around the 23-22^th^ centuries BCE. According to our results, the Bk-II population had an outstandingly high HG genetic ancestry level compared to other Bronze Age groups of the region. This can be traced back to two main sources, one to a WHG, and one to an EHG characteristic population, best modelled as FBC/GAC and Ukraine_EBA, however, likely both are only proximate to the actual source, which are yet to be described. The Y chromosome haplogroup I2a-L1229 can be linked to the FBC/GAC component, although this subgroup only appears first in Bk-II and related groups. The calculated admixture dates suggest the presence of a highly EHG characteristic population in Eastern Europe as late as the beginning of the Bronze Age. This EHG component shows the closest resemblance to Lithuanian Mesolithic individuals, but the best proxy for this population is probably missing from the published database, opening research questions for future studies.

Following the formation of the population represented here by Bk-II, it contributed to various populations in Central-Eastern Europe, whose genetic legacy persisted mostly in the region of today’s Hungary and Czechia at least until the end of the Bronze Age, and even to the end of the first millennium BCE in the Baltic region. This study does not disclaim any of the archaeological theories regarding the origin of the Kisapostag culture (Bóna, 1992, 1975; Črešnar, 2010; K. Zoffmann, 2008; Kiss, 2015), as the EHG core of the Kisapostag associated group fits really well with the Middle Dnieper origin, while further adaptation of cultural elements during their arrival and during their occupation in Transdanubia is plausible. The latter idea is further supported by the ^87^Sr/^86^Sr isotope ratio data which shows local isotope ratios for both sexes in both Bk-II and Bk-III. These results place back the time of their arrival a few generations, meaning that both local and southern cultural traits could explain the culture’s archaeological heterogeneity.

Bk-III was the direct descendant of Bk-II, forming cultural (Encrusted pottery) and genetic continuity for hundreds of years at the studied site. Observable dilution of HG ancestry in Bk-III compared to Bk-II can be connected to continuous female-biased admixture with nearby communities according to our and previous genetic (Freilich et al., 2021) and archaeological (Bóna, 1975; Vicze, 2011) evidence. The shift in pigmentation patterns is likely the result of this admixture.

In both periods, the homogeneity of paternal lineages suggest a patrilocal residence system, similarly to previously described social organisations (Freilich et al., 2021; Schroeder et al., 2019). However, ^87^Sr/^86^Sr isotope data shows local values for both sexes, which along with similar genomic makeup of females and males suggest exogamy most probably between villages of the same population. The overlap between outlier parallels of Bk-II/III and archaeological contact regions is also noteworthy, as it suggests smaller scale migrations of Kisapostag/Encrusted pottery individuals or groups along trading networks, such as mobility possibly connected to wandering merchants.

Notably, none but one (mtDNA haplogroup U5a1g) of the uniparental lineages are the same at the haplogroup level with the individuals from the Croatian Encrusted Pottery culture Jagodnjak site, despite high similarities in cultural traits, social structure and genomic composition of the communities (Freilich et al., 2021). This supports a regional patrilocal, clan-like superfamily structure of Kisapostag and Encrusted Pottery groups. This finding is particularly interesting in light of a strikingly different social structure observed among the 2100-1800 BCE Maros culture individuals from Mokrin, Serbia (Žegarac et al., 2021), that show extensive amounts of admixture related to the Kisapostag/Encrusted pottery culture.

The relatively limited presence of female and children burials in both Bk-II and Bk-III periods may suggest distinctive treatment or another (here undiscovered) burial group for women and children at the same site. However, in other cemeteries of the culture, *e.g.* Ordacsehi and Bonyhád in Hungary, males, females, and children were buried close to each other, suggesting high variance in burial practices (Hajdu, 2010; Somogyi, 2004; Szabó, 2010).

Although low genomic coverages did not allow fine SNP recovery, we did find evidence for malignant variants within all of the tested groups, and undoubtedly showed the presence of LHON and Jacob’s syndrome within Bk-II. While it only remains a possibility, the presence of autism risk factor in the CNTNAP2 gene, signs for severe bruxism, and the distinctive burial treatment of individual S45, suggest the actual onset of symptoms. Additionally, the disease panel we created and made freely available, can be extended and used in future studies, providing insight into past population health qualities.

Considering the unstructured age and biological relatedness distribution in the mass grave of Bk-III compared to Bk-II, the coetaneous death of eight people at least, the absence of traumatic or ritual events on bones, and the non-cremated nature of the burial, all signals a sudden tragic event in the Middle Bronze Age period (Encrusted Pottery population), most likely an epidemic, as first suggested based on the anthropological analyses (Köhler, 2006). Careful burial positions also suggest that the deceased were buried by their own community. Interestingly, comparative ^87^Sr/^86^Sr isotope analyses on the first and third molar of the individuals in the BK-III mass grave indicate that subadult males – including a severely disabled individual (S15) with hip dysplasia – left their community for a while and then returned to their birthplace prior to their death, raising further questions for future studies on prehistoric lifeways and social organisation.

## Materials and Methods

### Isotope analyses

Radiocarbon dating was performed at the HEKAL AMS C-14 facility of the Institute for Nuclear Research in Debrecen, Hungary (see Supplementary Information section 1.8). ^87^Sr/^86^Sr isotope measurements were performed in the ICER Centre, Institute for Nuclear Research Debrecen, Hungary and at Quinnipiac and Yale University, Connecticut, USA (see Supplementary Information section 1.9).

### Ancient DNA laboratory work

Petrous bones and teeth were taken from skulls for genetic investigation (Supplementary Table 1). Laboratory work was performed in a dedicated ancient DNA laboratory facility (Institute of Archaeogenomics, Research Centre for the Humanities, Eötvös Loránd Research Network, Budapest, Hungary). Each step was carried out in separate rooms under sterile conditions, during work protective clothing was used. Irradiated UV-C light, DNA-ExitusPlus™ (AppliChem) and/or bleach were applied for cleaning after and between work stages, and also, blank controls were utilised at all times.

Sample surfaces were cleaned by sandblasting and mechanically milled to powder. DNA extraction was performed according to (Dabney et al., 2013) with minor changes according to (Lipson et al., 2017). DNA extraction success was verified by PCR using mtDNA primer pairs (F16209-R06348; F16045-R06240). Half-UDG treated libraries were used according to (Rohland et al., 2015) with minor changes. Unique double internal barcode combinations were used for each library (Supplementary Table 1). Libraries were amplified with TwistAmp Basic (Twist DX Ltd) and purified with AMPure XP beads (Agilent). Then, concentration measurements were taken on Qubit 2.0 fluorometer, fragment sizes were checked on Agilent 4200 TapeStation System (Agilent High Sensitivity D1000 ScreenTape Assay).

Hybridisation capture method for mtDNA and 3k nuclear SNP was used besides whole genome shotgun, as described by (Csáky et al., 2020; Haak et al., 2015; Lipson et al., 2017). Bait production was based on (Fu et al., 2016) and (Csáky et al., 2020), the oligos as a pool was ordered from CustomArray Inc. Both for shotgun and capture libraries, universal iP5 and unique iP7 indexes were used. Sequencing was done on Illumina MiSeq and NovaSeq platforms with custom setup and 150, 200 and 300 cycles, respectively.

Additionally, we investigated Y chromosome STR profiles (17 markers) with AmpFLSTR® Yfiler® PCR Amplification Kit (Applied Biosystems), having one blank and one positive control at each reaction preparation. The workflow followed the recommended protocol except the PCR cycles were increased from 30 to 34 and reactions were halved in volume. Two repeats were done where at least 4 markers yielded results. Data analyses were carried out in GeneMapper® ID Software v3.2.1 (Applied Biosystems), results are summarised in Supplementary Table 3.

### Bioinformatic analyses

Illumina sequencing paired-end reads were processed by the *PAPline* https://github.com/ArchGenIn/papline. We used the GRCH37.p13 reference sequence to call the pseudohaploid genomes. For relatedness inferences we applied the *READ* software (Monroy Kuhn et al., 2018) and a custom script (named *MPMR*, see Supplementary Information section 2.3 and Supplementary Table 2). MtDNA analyses included phylogenetic analyses using the *MrBayes* v3.2.6 (Ronquist and Huelsenbeck, 2003) and the *BEAST* v1.10.4 (Suchard et al., 2018) software and diversity tests using the *Popgenome* (Pfeifer et al., 2014) *R* package, see Supplementary Information section 2.1. For Y chromosome haplogroup determination the *Yleaf* v1 (Ralf et al., 2018) software was applied. We used the *Network* v10.1.0.0 and *Network publisher* v2.1.2.5 (Bandelt et al., 1999; “Network Software,” 2008) programs for analysing the network of STR data, see Supplementary Information section 2.2. We discarded individuals S2, S4, S5, S6, S17 and S20 from the population genetic analyses due to low genomic coverages and/or being first degree relative of other samples. The Principal Component Analysis was made by the Eigensoft smartpca software (Patterson et al., 2017) using the Human Origins Panel SNP set (Patterson et al., 2012), for other analyses the 1240k array SNP set (Mathieson et al., 2015) was used for variant calling, for results, see Supplementary Table 7. For investigating ancestry estimates we used supervised admixture analysis calculated by the *ADMIXTURE* v1.3.0 software (Alexander et al., 2009). *f*-statistics and qpAdm were performed using the *admixr* v0.9.1 (Petr et al., 2019) and the *admixtools* v2.0.0 (Patterson et al., 2012) *R* packages. The timing of the admixture events were inferred by using the *DATES* software (Chintalapati et al., 2022).

## Data availability

All studied data are cited in the article and/or Supplementary Information and tables. New sequencing data are deposited in the European Nucleotide Archive (ENA) under accession number PRJEB49524.

## Acknowledgement

This study was funded by the Hungarian Academy of Sciences through the Momentum Mobility research project (LP2015-3/2015). This paper and A.S-N. was supported by the János Bolyai Research Scholarship of the Hungarian Academy of Sciences. E.A. ‘s work was supported by the National Research, Development and Innovation Office, Hungary (NKFIH) grant (PD-19/131839). M.B. ‘s work and the analyses of the BAD002 sample was supported by a NKFIH grant under project code K-18/128413. We would like to greatly thank restorer Zsuzsanna Herceg and digital artist Fanni Gerber for the artwork of individual S13, and Zsolt Réti for the digital reconstruction of the mass grave (obj. B-938).

## Author contribution

D.G, V.K., A.Sz-N. conceived and designed the experiments. D.G. processed the sequencing data, created the *PAPline*, and performed downstream bioinformatic analyses. B.Sz. did all molecular laboratory work. O.Sz. created the mtDNA database for phylogenetic analyses. B.E. obtained Y chromosome STR data. B.Gy. and E.A. optimised genetic analyses. J.I.G., A.H., L.P., M.M. and I.M. performed Sr isotope analysis. G.K., Sz.F., V. K. and M.B. evaluated the archaeological context. B.G.M., Á.K. and K.K. did the anthropological examination of the remains. Á.K. made the facial reconstruction. V.Sz., I.M. and M.M. performed ^14^C calibrations and modelling. B.G.M. sampled the remains. E.A., V.K. and A.Sz-N. jointly supervised the research and wrote the paper with D.G. All authors provide critical feedback for this study and contribute to the final manuscript.

## Ethics declarations

The authors declare that they had requested and got permission for the destructive bioarchaeological analyses of the archaeological material in the study from the stakeholders, excavator and processor archaeologists.

