## Supplementary Information for "Interdisciplinary analyses of Bronze Age communities from Western Hungary reveal complex population histories"

### Table of content

**1) Archaeological, isotope and anthropological data** 1
1.1) Summary of the archaeological context and preliminary genetic analyses of BAD002 1
Fig. S.1.1.1 4
1.2) Overall description of Balatonkeresztúr site 4
Fig. S.1.2.1 4
1.3) Overall description of the studied era 4
1.4) Somogyvár-Vinkovci culture 5
Fig. S.1.4.1 7
1.5) Kisapostag/Earliest Encrusted Pottery culture 7
Fig. S.1.5.1 8
Fig. S.1.5.2 8
Fig. S.1.5.3 8
Fig. S.1.5.4 9
Fig. S.1.5.5 9
Fig. S.1.5.6 10
Fig. S.1.5.7 11
Fig. S.1.5.8 12
Fig. S.1.5.9 13
Fig. S.1.5.10 14
Fig. S.1.5.11 16
1.6) Transdanubian Encrusted Pottery culture 18
Fig. S.1.6.1 19
Fig. S.1.6.2 19
1.7) Anthropological and paleopathological analyses 20
1.8) Radiocarbon dates of the Balatonkeresztúr site 20
Fig. S.1.8.1 21
Fig. S.1.8.2 21
Modelled radiocarbon dates of samples from the Balatonkeresztúr site, a multiplot for the Kisapostag period (BK-II) radiocarbon series. 21
Fig. S.1.8.3 21
Fig. S.1.8.4 21

Fig. S.1.8.5 23
Fig. S.1.8.6 24
Fig. S.1.8.7 24
1.9) Sr isotope data of Balatonkeresztúr site 24
SI Table 1 28
**2) Uniparental genetics and relatedness 28**
2.1) The mitochondrial DNA haplogroups and their phylogenetic evaluation 28
Fig. S.2.1.1 29
Fig. S.2.1.2 29
Fig. S.2.1.3 29
Fig. S.2.1.4 29
Fig. S.2.1.5 29
Fig. S.2.1.6 29
Fig. S.2.1.7 29
Fig. S.2.1.8 30
Fig. S.2.1.9 30
Fig. S.2.1.10 30
Fig. S.2.1.11 30
2.2) Y chromosomal haplogroups and STR network analyses 30
Fig. S.2.2.1 36
Fig. S.2.2.2 36
Fig. S.2.2.3 36
Fig. S.2.2.4 36
Fig. S.2.2.5 36
Fig. S.2.2.6 36
2.3) Relatedness analysis 36
**3) Phenotype assessment 41**
3.1) Genetic sex and aneuploidy 41
3.2) Variant discovery 41
3.2.1) SNPs of pigmentation 41
3.2.2) SNPs of clinical significance 42
3.2.2.1) Lig4 syndrome 42
3.2.2.2) Diabetes and related disease susceptibility 42
3.2.2.3) Hereditary spastic paraplegia 42
3.2.2.4) Autism 15 (AUTS15) susceptibility 42
Fig. S.3.2.2.4. 42
**4) Facial reconstruction 42**
SI Table 2 45
Fig. S.4.1 47
Fig. S.4.2 47
Fig. S.4.3 47
Fig. S.4.4 47
Fig. S.4.5 47
Fig. S.4.6 47
Fig. S.4.7 47

Fig. S.4.8 47
Fig. S.4.9 48
**5) Whole genome analyses 48**
5.1) PCA 48
5.2.1) Admixture 48
Fig. S.5.2.1 52
5.2.2) Admixture based outlier detection 52
5.3) f4-statistics 52
Fig. S.5.3.1 53
Fig. S.5.3.2 53
Fig. S.5.3.3 53
Fig. S.5.3.4 53
Fig. S.5.3.5 53
5.4) DATES analyses 53
SI Table 5.4 58
5.5) f3-statistics 58
Fig. S.5.5.1 60
Fig. S.5.5.2 60
SI Table 5.5.1 60
5.6) qpAdm 61
5.6.1) BAD002 63
5.6.2) Balatonkeresztúr site 64
5.6.2.1) Bk-I 64
5.6.2.2) Bk-II 65
Fig. S.5.6.2.2 65
5.6.2.3) Bk-III 65
**6) References 66**

### 1) Archaeological, isotope and anthropological data

by Mária Bondár, Szilvia Fábián, Viktória Kiss, Kitti Köhler, Gabriella Kulcsár, Ágnes Kustár

#### 1.1) Summary of the archaeological context and preliminary genetic analyses of BAD002

The Late Copper Age Baden culture inhabited the major part of the Carpathian Basin from 3500 BCE until 2800/2700 BCE. The burials of this population reflect extremely diverse mortuary practices. The graves include both inhumation and cremation burials. Very often, a grave contained several burials, but skull burials and symbolic graves (the latter often empty or containing but a few artefacts, for example a wagon model) are also known. Some graves contained both human and animal burials. A mainly sedentary lifestyle was characteristic for this cultural complex with smaller and larger settlements. During the centuries of this culture, major changes occurred in the life of Eurasian people; many innovations (e.g. the secondary products revolution, wagons and wheels) were made that had lasting effects on the history of humanity. These novelties spread fast and wide and further deepened economic and social disparities within communities that were reflected in burials (Bondár, 2018, 2012).

Skull cults were prevalent among Mesolithic and Neolithic groups in Europe and the Middle East as well. In the Late Copper Age Baden culture this practice was rare; however, Feature 415 at the Balatonlelle-Rádpuszta site includes a 7-8 years old male individual surrounded by four skulls and one fragmented skull of a child and a young adult male (Bondár, 2020). BAD002 corresponds to This site is located approximately 30 km away from the Balatonkeresztúr-Réti-dűlő site and the child skeleton, referred to as BAD002, was included in this study for comparison purposes (Figure S.1.1.1).

Two of the skulls were presumably close relatives of BAD002 according to mtDNA results. The skulls lack mandibles and were considered secondary because they were placed into the burial of BAD002, contemporary with the funeral act. Since previous findings of skull cults appear throughout Europe, the burial practice does not help pinpoint a specific cultural origin; however, the presence of copper and black jet beads (Bondár et al., 2021), as well as the mitochondrial haplogroup U5b3 from two of the skulls provide linkages to western European populations (Bondár and Szécsényi-Nagy, 2020). Preliminary results revealed that BAD002 had an outstandingly high Hunter Gatherer (HG) component compared to known Neolithic and Copper Age groups from the Carpathian Basin, therefore we included it in this study.

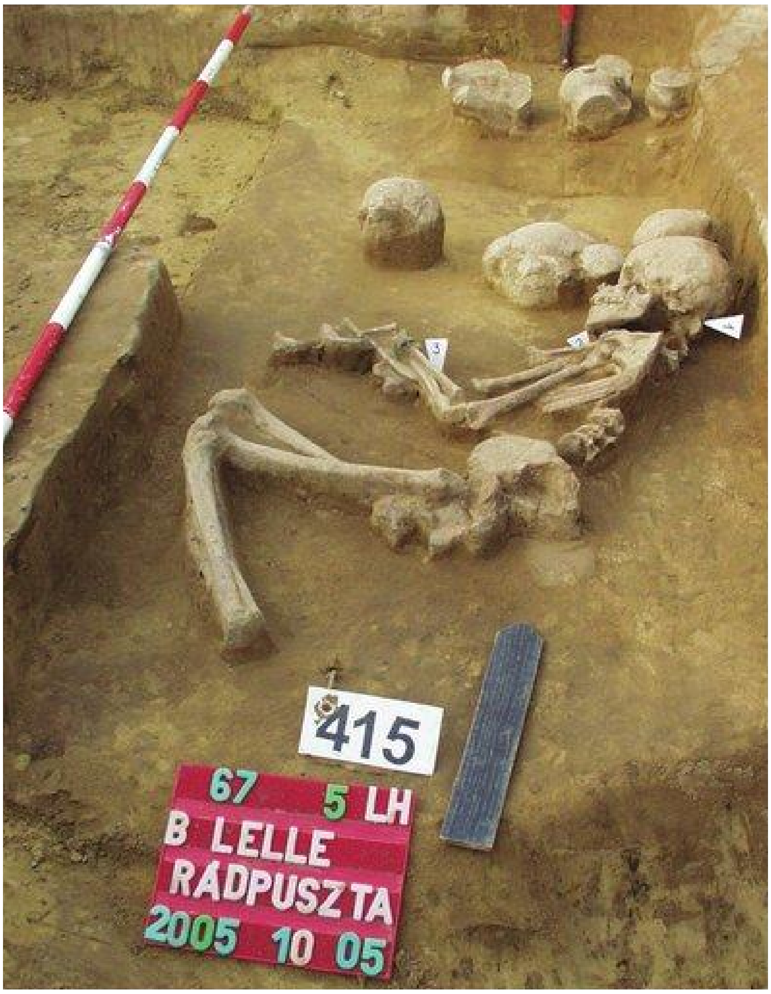


*Fig. S.1.1.1*

*Balatonlelle-Rádpuszta, the complete skeleton of BAD002 in Feature 415 surrounded by pottery and skulls of children and an adult individual. According to Bondár & Szécsényi-Nagy 2020 (Bondár and Szécsényi-Nagy, 2020), two of the skulls possess the same mitochondrial haplogroup as BAD002, suggesting that at least some of the skulls belong to close relatives of this young male individual. Remains are dated to 3530-3370 cal BCE (95.4% CI).*

#### 1.2) Overall description of Balatonkeresztúr site

The Balatonkeresztúr-Réti-dűlő site (Fig. S.1.2.1) is located 3 km from the southern shore of Lake Balaton. In 2003–2004 an area of 45,000 m$^2$ was investigated preceding the construction of the M7 motorway. As a result, the traces of occupations associated with nine archaeological cultures spreading over eight chronological periods were identified from the analysis of 2,976 archaeological features (Fábián and Serlegi, 2009; Honti et al., 2006). This includes: the Middle and Late Copper Age (Balaton-Lasinja, Furchenstich, Boleráz and Baden cultures), Early Bronze Age (Somogyvár-Vinkovci and Kisapostag/earliest Encrusted Pottery cultures), Middle Bronze Age (Transdanubian Encrusted Pottery culture), Late Iron Age (La Téne D), Migration period (Langobards), Árpádian Age (12-13$^{th}$ century) and Late Middle Ages (13-15$^{th}$ century (Honti et al., 2004)). Human remains were discovered from settlement features without diagnostic finds, dated by radiocarbon sampling: one pit (Somogyvár-Vinkovci) contained bones of one individual, while another pit (Transdanubian Encrusted Pottery) contained the remains of 8 human bodies. East of the settlement of the Kisapostag culture a small cemetery with 11 skeletals was discovered. Most of the burials contained no grave goods, except for two with small bronze ornaments (Honti et al., 2004).

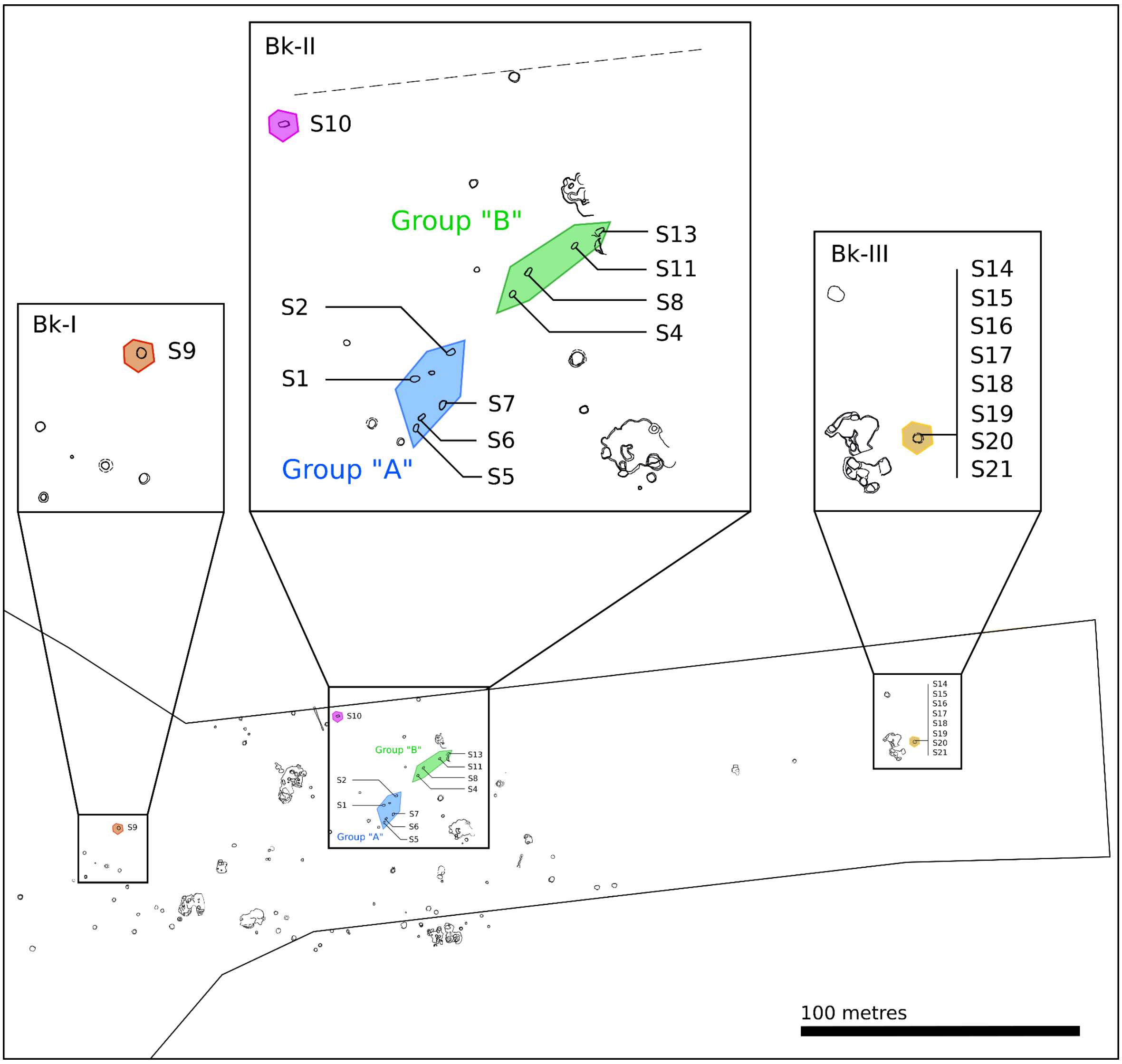


*Fig. S.1.2.1*

*Map of Balatonkeresztúr-Réti-dűlő site, where graves and grave groups are shown in separate enlarged boxes. The only child grave is situated near to the edge of the excavation site, whereas S45 is situated hundreds of metres away from the rest of the burials (not shown), for possible reasons see the 'Discussion' section in the main text of the manuscript. The mass grave of Bk-III is also situated far from the core of Bronze Age objects (uncoloured) at the lower point of the site.*

#### 1.3) Overall description of the studied era

The first thousand years of the Bronze Age in the Carpathian Basin – between 2600/2500-1500 BCE – is an important period in connection with the spread of bronze metallurgy, new pottery styles, development of new settlements and emerging social inequalities in Central Europe. During the Early Bronze Age in Hungary (2500-2000/1900 BCE; parallel with the onset and older phase of the Central European Early Bronze Age, Reinecke Br A0 and A1) arsenical bronze axes and daggers, as well as ornaments made of pure copper were common. After 2000/1900 BCE tin bronze weapons and tools appeared in all spheres of everyday life (Dani, 2013; Dani et al., 2016; Fischl et al., 2013). Two different burial practises, cremation and inhumation can be observed during the Early Bronze Age (Kiss, 2020a; Kulcsár, 2009). After the beginning of the Middle Bronze Age in western Hungary around 2000/1900 BCE (parallel with the beginning of Central European developed Early Bronze Age, Reinecke Br A2) cremation burials were applied nearly exclusively. For example, at the Bonyhád-Biogas factory site, only 6.5% of the burials were inhumations from the earliest period of the cemetery (Hajdu et al., 2016). Considering the popularity of cremation as a mortuary practice during the Bronze Age of this region (and it's destructive taphonomic impacts to human bone), when inhumations are present in the archaeological record they present an important source of material to conduct physical anthropological analyses, as well as stable isotope and ancient DNA studies.

#### 1.4) Somogyvár-Vinkovci culture

In the first part of the Early Bronze Age in western Hungary, hilltop/hillfort settlements as well as smaller and larger villages of this culture can be distinguished. Only a very small number of burials are known from this period, which include urn cremations and several inhumations. In addition to these burials, a few inhumations from settlement pits were also discovered (Kulcsár, 2009). At Balatonkeresztúr-Réti-dűlő only one burial (individual S9) can be associated with this archaeological culture according to its $^{14}C$ date (3929±30 BP, 2560-2290 cal BCE, 95.4% CI). Remains of this individual (Fig. S.1.4.1) were discovered in a settlement pit, lying in a special position, on his back and with the knees and head on the left side, oriented N/NE-S/SW (Preda-Balanica et al., 2020). Within the pit, pottery shards and animal bones were found, indicating that it likely functioned as a waste pit. Bones were preserved in worse condition than other human remains discovered at the site, suggesting bad life conditions. The skeletal remains of this individual consist of a moderately preserved skull and well preserved bone material. Determined as a male of 35-40 years with a calculated height of 154.8 cm, his anthropological features, mainly his skull shape (ultradolichokran curvoccipital) resemble the preceding Copper Age population (Köhler, 2011, 2009; Köhler et al., 2017). No particular pathological condition could be observed, except some cribra orbitalia (porotic lesions on the upper wall of the eye-orbit), which may have resulted from the increased erythrocyte number that signal a wide variety of pathological conditions from malnutrition to malaria (Marciniak et al., 2021).

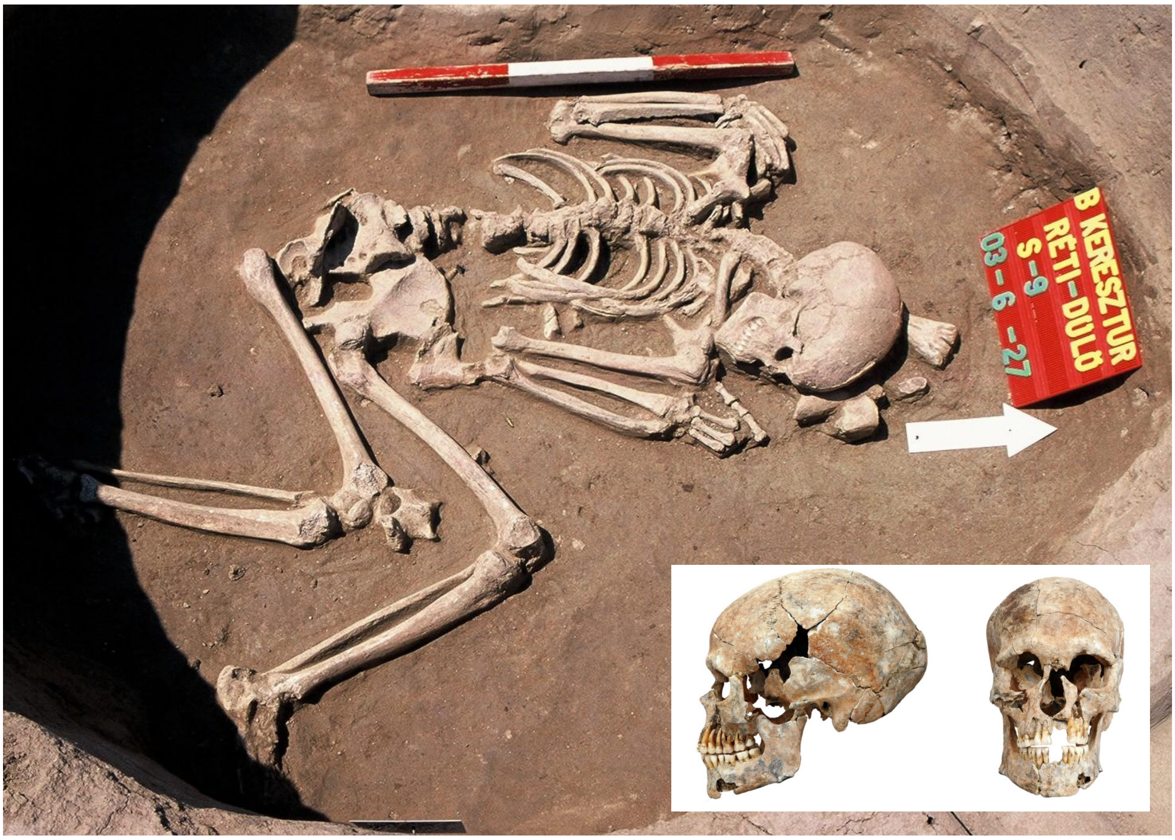


*Fig. S.1.4.1*

*Grave of individual S9 surrounded by animal remains, such as cattle metatarsus under his skull. His burial position shows resemblance to people of high steppe ancestry (Preda-Balanica et al., 2020), which was later confirmed by aDNA analysis. Photo by Szilvia Fábián.*

#### 1.5) Kisapostag/Earliest Encrusted Pottery culture

The two-tiered settlement structure mentioned above, with scattered dwellings at smaller and larger villages, and sometimes hilltop/hillfort sites and settlements surrounded by ditches, is characteristic for the communities of the Kisapostag/earliest Encrusted Pottery. Open settlements, like the ones discovered in Kaposvár and Balatonkeresztúr, consist of a few houses, storage and refuse pits. Metal finds are usually made of fahlore copper, rich in arsenic, antimony and silver impurities; tin bronzes can be observed in a few cases only at the end of the Early Bronze Age (Kiss, 2015, 2012; Somogyi, 2004). Before the 1990s, it was assumed that cremation (urn burials and scattered cremations) was the only burial custom associated with the Kisapostag/earliest Transdanubian Encrusted Pottery culture. In recent decades, around 90 inhumation burials of this culture have also been documented (Kiss, 2020a). Anthropological and bioarchaeological analyses of these inhumation burials remain limited (Hajdu et al., 2016; Kiss, 2020a; Somogyi, 2004).

From the Balatonkeresztúr site, there are 12 inhumation burials associated with this culture according to archaeological features and $^{14}$C dates (Supplementary Table 1). The graves, with bodies placed in flexed position, laying on their side, were found in two groups, east from the settlement of the same period. Group A includes individuals: S1, S2, S3, S5, S6, S7, N=6;Group B includes individuals: S4, S8, S11, S13, N=4. Two single inhumations located outside of the burial groups (individuals S10 and S45) were also identified Individual S3 was not sampled due to the complete destruction of the poorly preserved remains during excavation. Group A consists of three juveniles and three adults, five males and a female (individual S7), while in Group B we find one juvenile male, and three adults, two males and one female (individual S13). One inhumation, found a bit further from the two burial groups, belongs to a young male individual (S10), while another older male was found further west (S45). Three graves (individuals S10, S12 (cremated), and S13) contained metal ornaments as grave goods. Many samples were only preserved as scattered and heavily destroyed remains, and most skeletons show slight or modest distortion. The overall pathological makeup of the population does not show much evidence for disease and the average adult biological age is high. Detailed anthropological analyses along with archaeological artefacts associated with each burial can be found in the Supplementary Figures S.1.5.1-S.1.5.11.

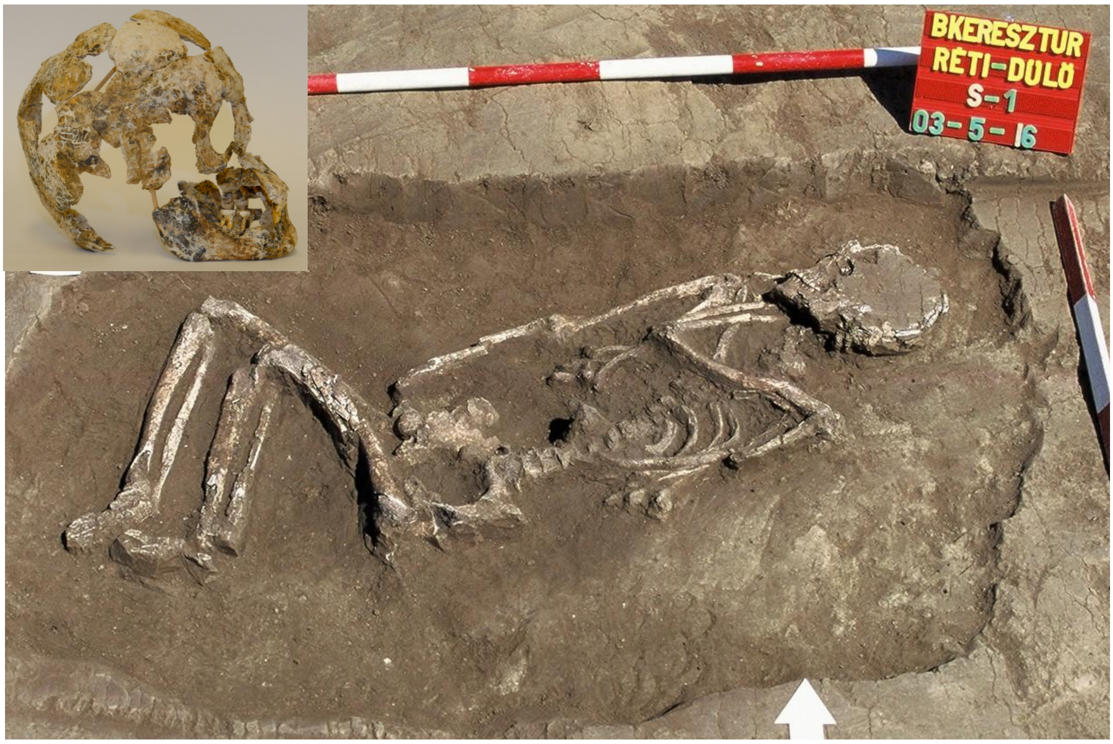


*Fig. S.1.5.1*

*Grave of individual S1, Group A. The body was placed on the right side in a flexed position, oriented NE-SW. One piece of flint was found in the grave pit. The remains consisted of fragmented post-cranial bones and a skull, $^{14}C$ dated to 3611 ± 31 BP (2120-1880 cal BCE, 95.4% CI), and belonged to a male of over 40 years. Determination of age at death was based on tooth abrasion: level 5; sexualisation index: - 29. Several teeth remained: three upper and one lower molar were decayed (caries); the lower right first molar is ante mortem missing, indicated by the healed alveolus. Other pathological conditions were not observed. The skull has a brachycranic and likely planoccipital shape, which is an almost ubiquitous feature of this group. Skull reconstruction and photo by Dániel Gerber, grave photo by Szilvia Fábián.*

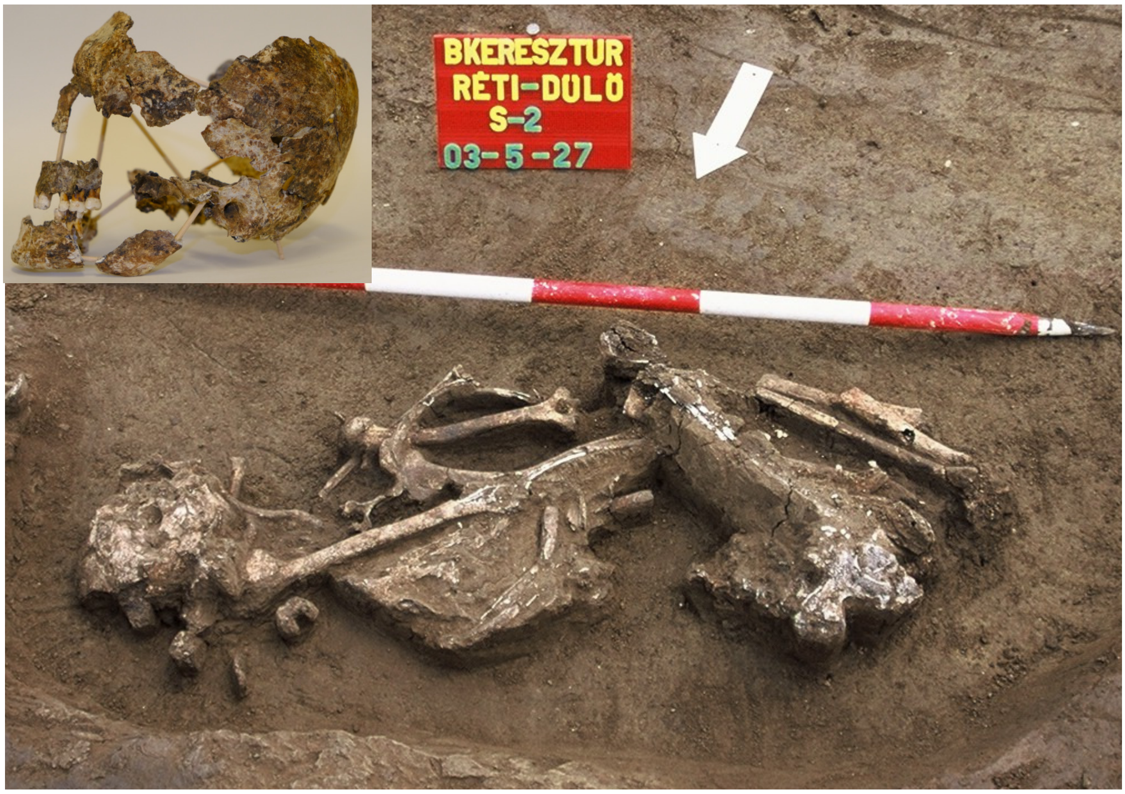


*Fig. S.1.5.2*

*Disturbed grave of individual S2, Group A. The remains and endogenous DNA are poorly preserved. The body was placed on the left side in a flexed position, oriented NE-SW. Several non-diagnostic pieces of Bronze Age pottery were found in the burial pit. The remains consisted of fragmented bones, and is dated to 3609 ± 32 BP (2120-1880 cal BCE, 95.4% CI). S2 is a male of 30-35 years (determination of age at death based on tooth abrasion: level 3; sexualisation index: -0,50). No molar decay was observed on the remaining 12 teeth. His skull type is brachycranic and likely planoccipital. A rare anatomical variation of perforated fossa olecrani can be observed on the left humerus. Skull reconstruction and photo by Dániel Gerber, grave photo by Szilvia Fábián.*

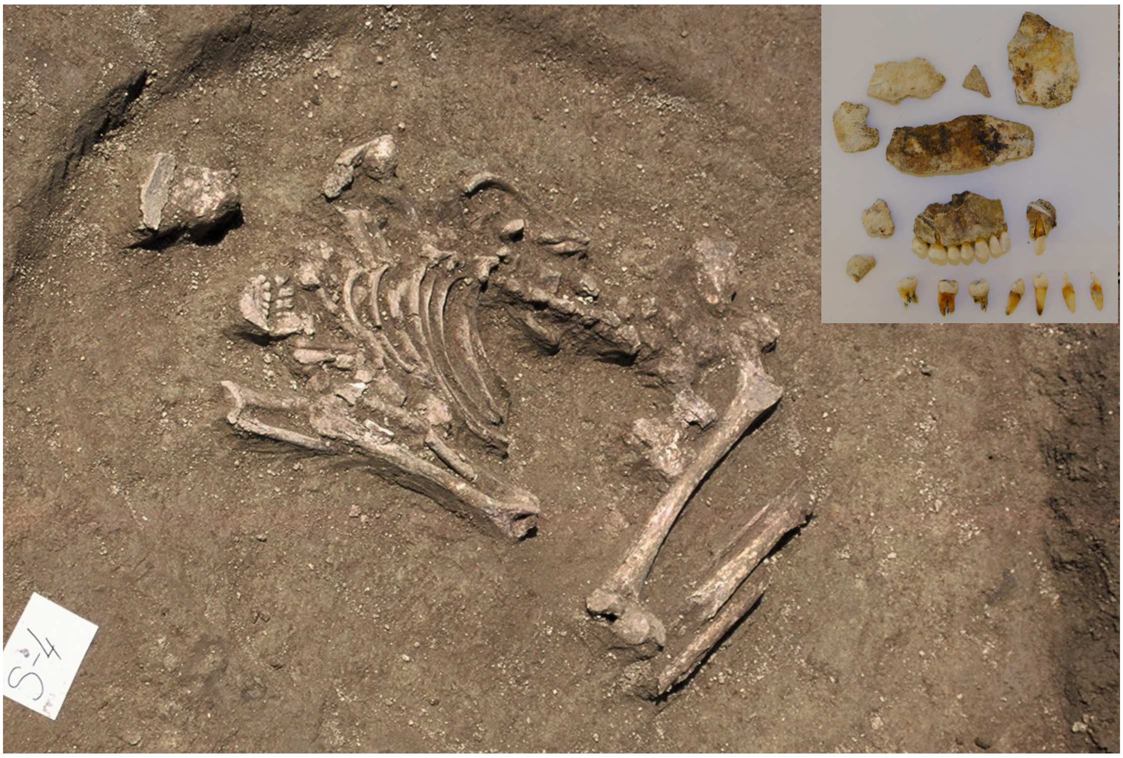


*Fig. S.1.5.3*

*Grave of individual S4, Group B. The skeletal remains and the DNA were preserved poorly. The deceased was placed on its right side in a flexed position, oriented slightly NE-SW. Human remains only consisted of a small amount of fragmentary bone material and teeth. The remains belong to a 17 to19 year-old male individual (determination of age at death based on muscular adhesion surfaces, and tooth abrasion: level 2). Sixteen teeth were recovered, with no visible signs of pathological conditions. One boar incisor and fragmentary pottery sherds were associated with the grave. $^{14}$C dating was not performed (the remains can be dated by the genetically determined father, individual S8, as it is shown in the 'Results' section in the main text). Photo by Szilvia Fábián.*

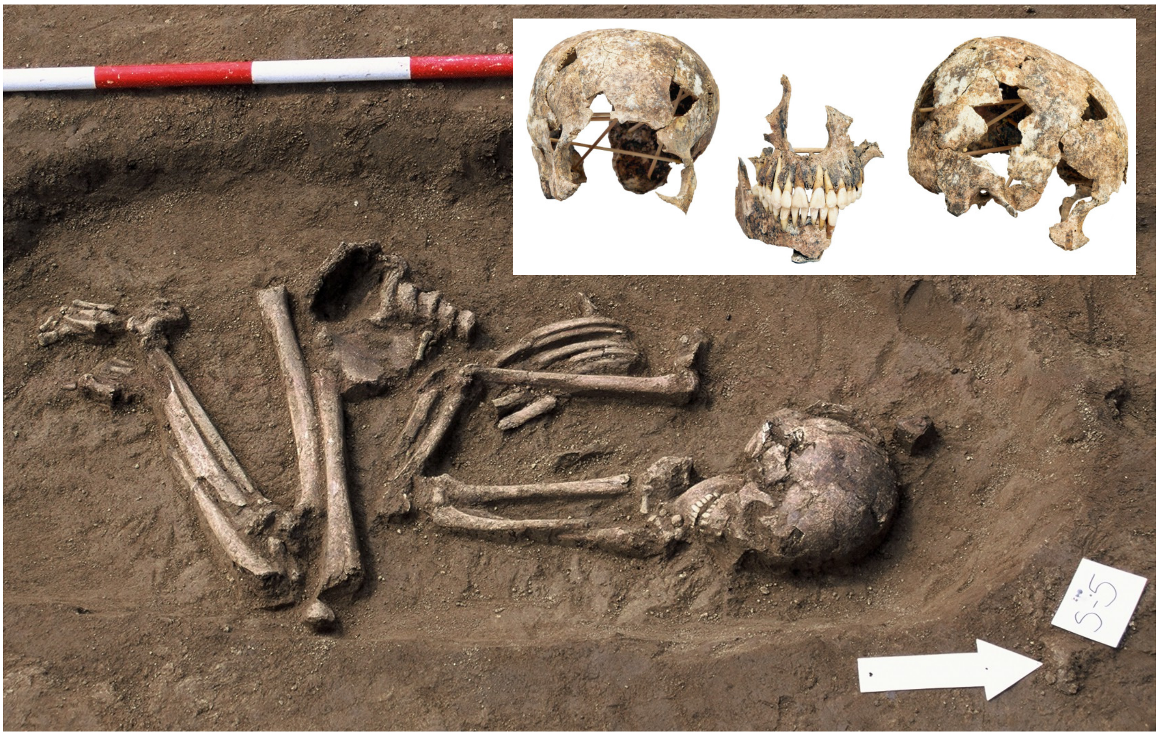


*Fig. S.1.5.4*

*Grave of individual S5, Group A. Moderately preserved remains of a 16 to18 year-old male individual in a flexed position, oriented N-S. Several non-diagnostic Bronze Age pottery sherds were found in the grave. Determination of age at death was based on muscular adhesion surfaces, and tooth abrasion (level 2-3). Thirty-two teeth were recovered. Skull type is brachycranic and likely planoccipital. Besides a severely decayed molar (caries on the left upper first molar) no other pathological conditions were observed. $^{14}C$ dating was not performed (the remains can be dated by the genetically determined father, individual S11, as it is shown in the 'Results' section in the main text of the manuscript). Skull reconstruction by Dániel Gerber, grave photo by Szilvia Fábián.*

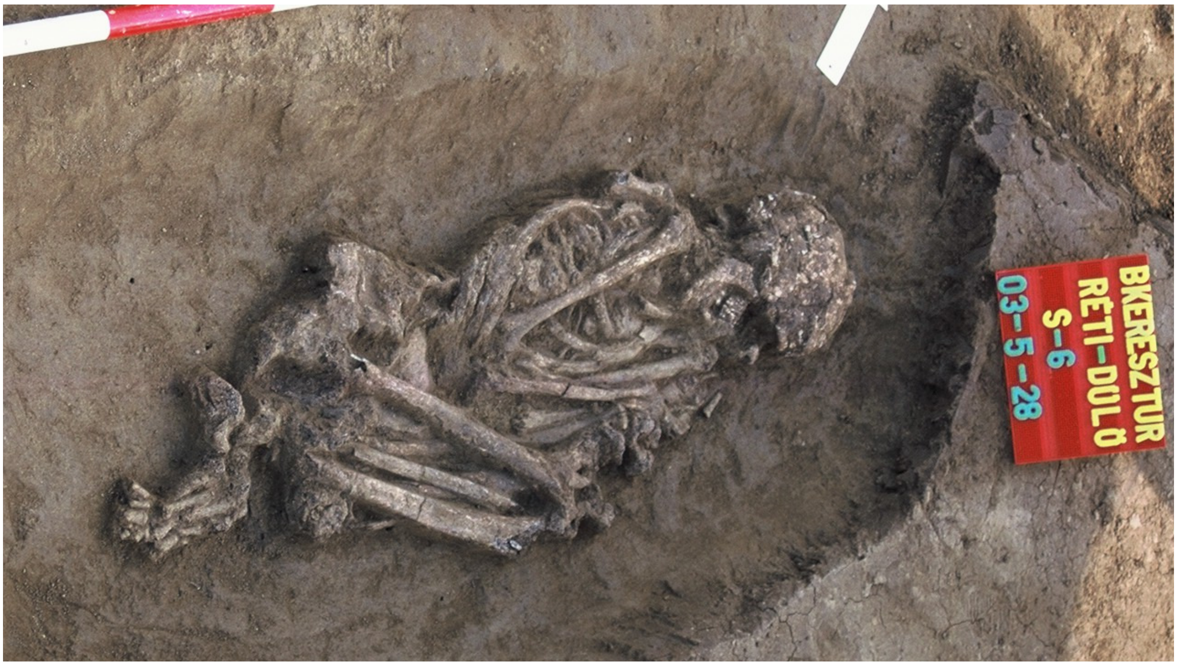


*Fig. S.1.5.5*

*Grave of individual S6,Group B. The skeletal remains and the DNA were preserved poorly. The body was placed on the left side in a heavily flexed position, oriented NE-SW. The moderately preserved bone fragments were $^{14}C$ dated to 3571 ± 31 BP (2030-1770 cal BCE, 95.4% CI), and belonged to a 17 to18 year-old male individual (sexualisation index: +0,71; determination of age at death based on muscular adhesion surfaces, and tooth abrasion: level 2). Twenty four teeth were present; besides one decayed molar (caries present on the right lower first molar) no other pathological conditions were observed on the skeletal remains. Photo by Szilvia Fábián.*

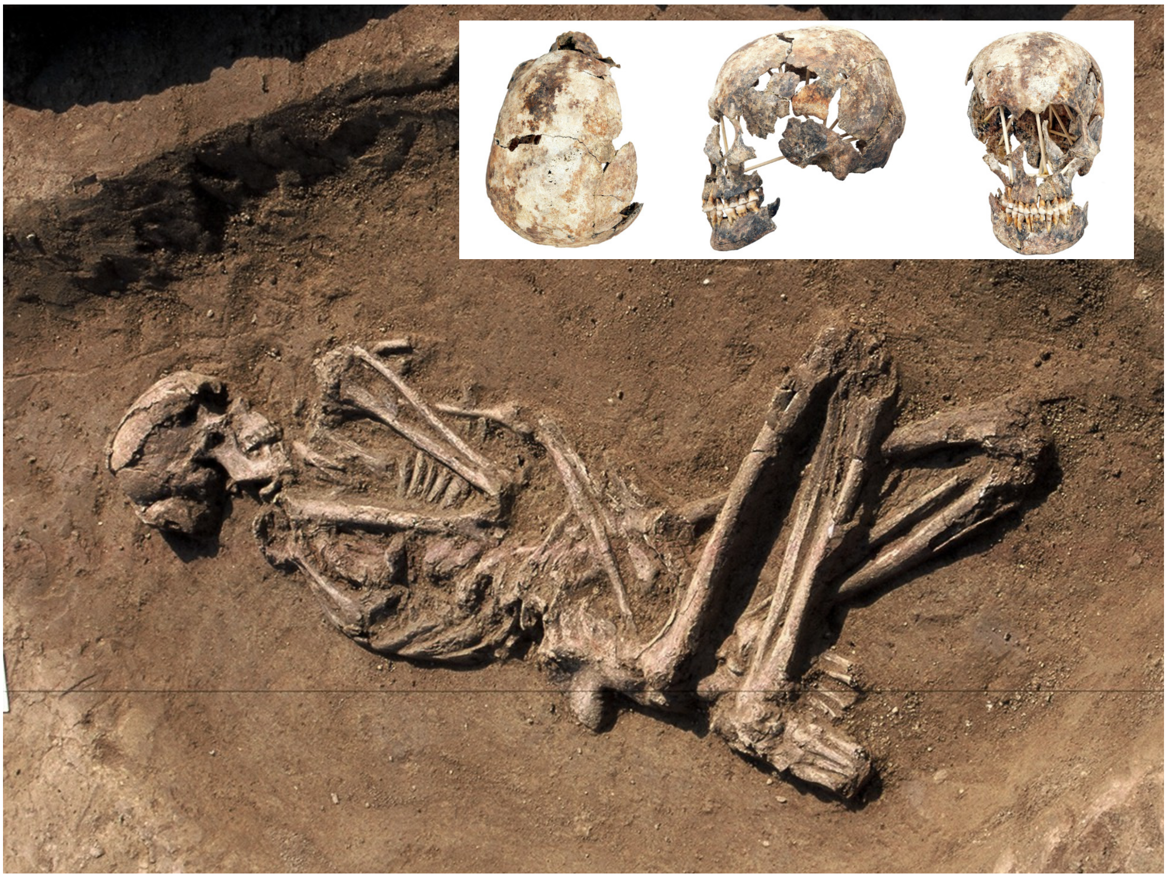

*Fig. S.1.5.6*

*Grave of individual S7, Group A. Moderately preserved remains of a 35 to 50 year-old female (sexualisation index: -0,18; determination of age at death based on sutures ossification, and tootхth abrasion), $^{14}C$ dated to 3611 ± 32 BP (2120-1880 cal BCE, 95.4% CI). The skull was of brachycranic type with a long facial structure. Twenty eight teeth were present (with no caries); however, some teeth were ante mortem missing (right lower second premolar and second molar, and left lower second premolar). Extended porotic lesions were found in the skull (os frontale et parietale), in addition to some enthesopathy, i.e. bone spikes caused by heavy labour on the right calcaneus (left is not examined because of its fragmentary condition). No grave goods were recovered from the grave besides some non-diagnostic Bronze Age pottery sherds. Skull reconstruction by Dániel Gerber, grave photo by Szilvia Fábián.*

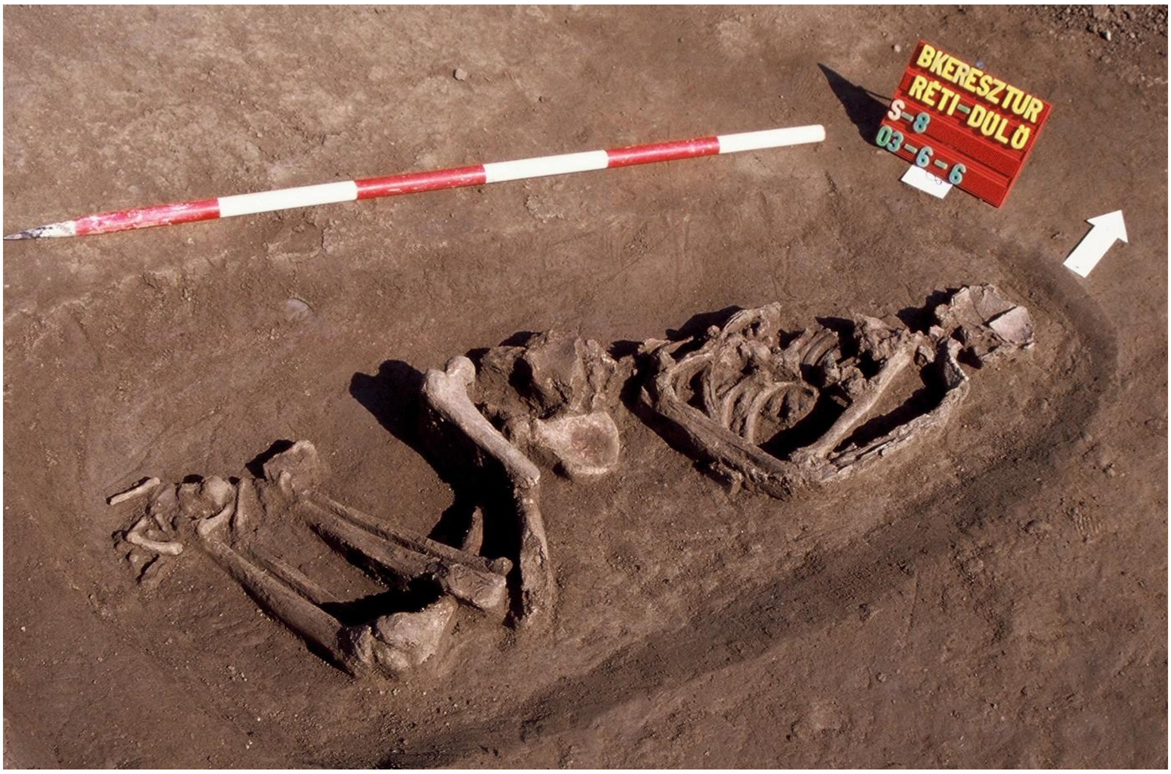


*Fig. S.1.5.7*

*Grave of individual S8, Group B. The body was placed on the left side in a flexed position, oriented NE-SW. The poorly preserved bones belonged to a 30 to 40 year-old male (sexualisation index: +0,80; determination of age at death based on abrasion of teeth (level 4), sutures ossification, and the surface of facies auricularis) with a calculated height of 168.2 cm. Just like individual S2, the body had an anatomical variation of perforated fossa olecrani on the right humerus, likely as a result of biological relationship (individuals S2 and S8 were 2nd degree relatives). Only four teeth remained, showing no signs of decay (caries). On the diaphysis of his right tibia, periostitis is observed, while both calcanei show signs of enthesopathy. Photo by Szilvia Fábián.*

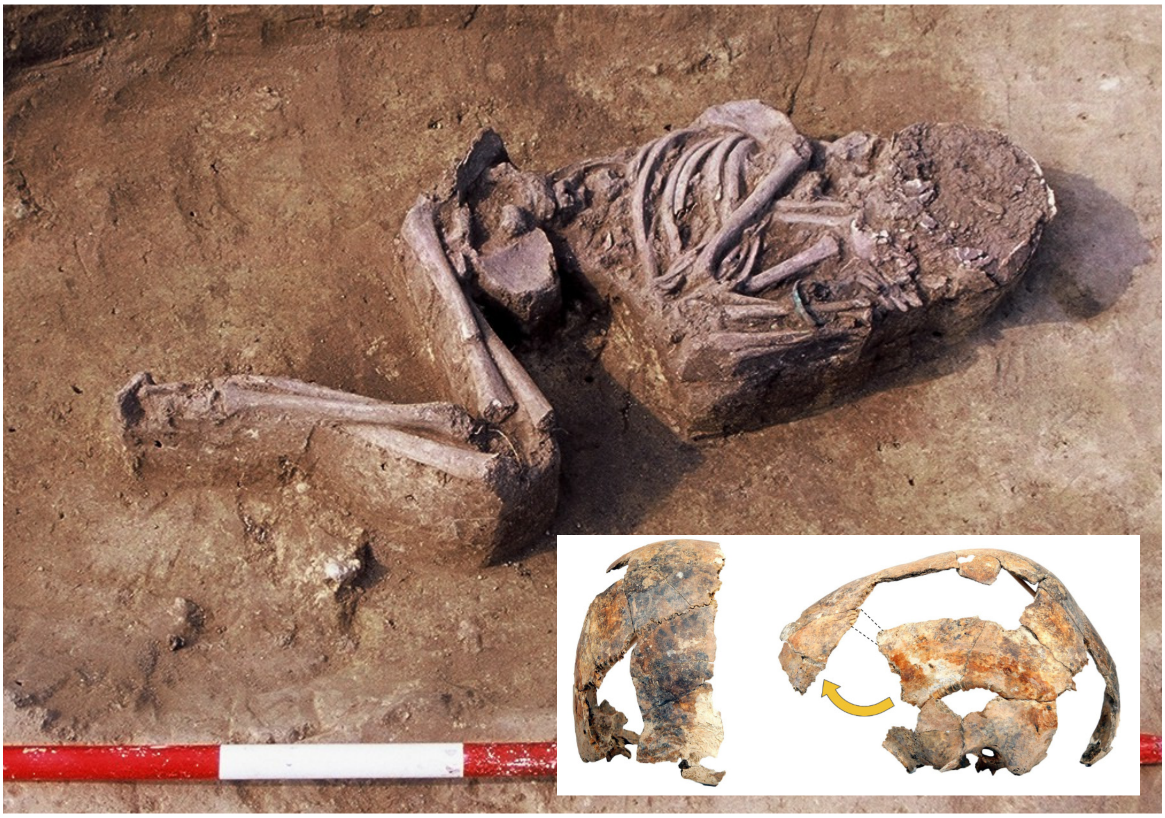

*Fig. S.1.5.8*

*Grave of individual S10, the only child grave from Bk-II. Moderately preserved remains of a 7 to 8 year-old male individual (determination of age at death based on teeth, and size of long bones). $^{14}C$ dated to 3661 ± 30 BP (2140-1940 cal BCE, 95.4% CI). The body was placed in a flexed position, oriented NE-SW. The bones do not show any sign of pathological conditions. A copper or bronze bracelet on the arm shows higher social status for this individual. Skull reconstruction by Dániel Gerber, grave photo by Szilvia Fábián.*

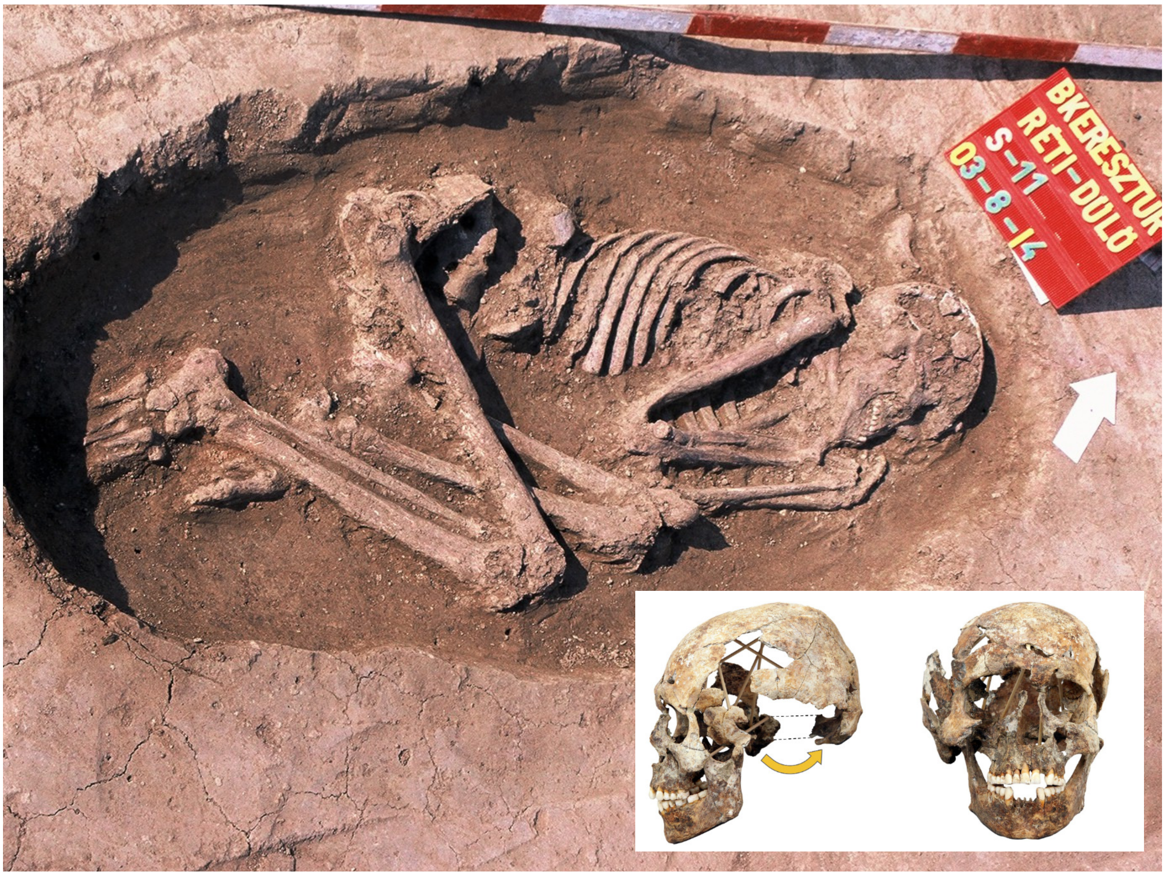


*Fig. S.1.5.9*

*Grave of individual S11, Group B. Moderately preserved remains of a 34 to 43 year-old male (sexualisation index: +1,16, determination of age at death based on the trajectory of proximal epiphysis of the humerus and femur and abrasion of the remaining 26 teeth: level 3). $^{14}C$ dated to 3705 ± 30 BP (2200-1980 cal BCE, 95.4% CI). The body was placed in a flexed position and oriented to the NE-SW. The dolichocranic skull was highly deformed under soil pressure. The bones are distorted, similarly to many other individuals from this archaeological culture. The calculated height of the body is 171.9 cm, and the individual had a severe walking condition associated with his left leg owing to an early life injury or a developmental disorder. The grave contained some non-diagnostic Bronze Age pottery sherds. Skull reconstruction by Dániel Gerber, grave photo by Szilvia Fábián.*

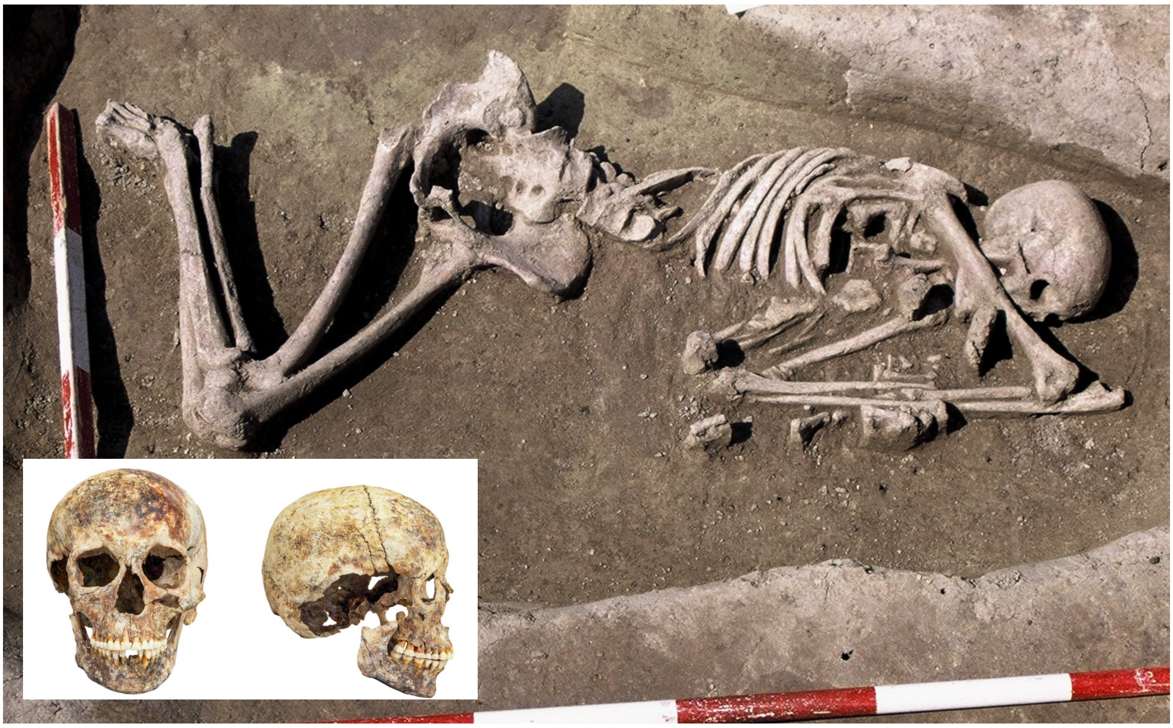

*Fig. S.1.5.10*

*Grave of individual S13, Group B. Well preserved remains of a 35 to 45 year-old female (sexualisation index: -0,81, determination of age at death is based on sutures ossification, the surface of facies symphyses and tooth abrasion: level 4-5). $^{14}C$ dated to 3618 ± 30 BP (2120-1890 cal BCE, 95.4% CI). The body was placed on the left side in a flexed position, oriented N-S. The remains had a strange body placement compared to other individuals, implying special treatment. The skull is of brachycranic type, the calculated height of the body is 158.7 cm, and the individual had a rare anatomical variation of extra sutural bones and maxillary prognathy. Twenty nine teeth remained, with caries observable only on a lower left second molar. A lower left third molar had fallen out ante mortem. The only pathological condition was one missing and one decayed tooth. Analyses on the pelvis bone suggest no or few birth giving events during her lifetime. The individual had several fragments of metal beads made of copper sheet at the left side of the head, along with a small burnt animal bone fragment, as well traces of green patina on the right side of the skull. The remains and skull from this burial are preserved the best among all individuals from the site enabling facial reconstruction, grave reconstruction, and a detailed anthropological description using genetic data, also see the 'Facial reconstruction' section. Skull reconstruction by Dániel Gerber, grave photo by Szilvia Fábián.*

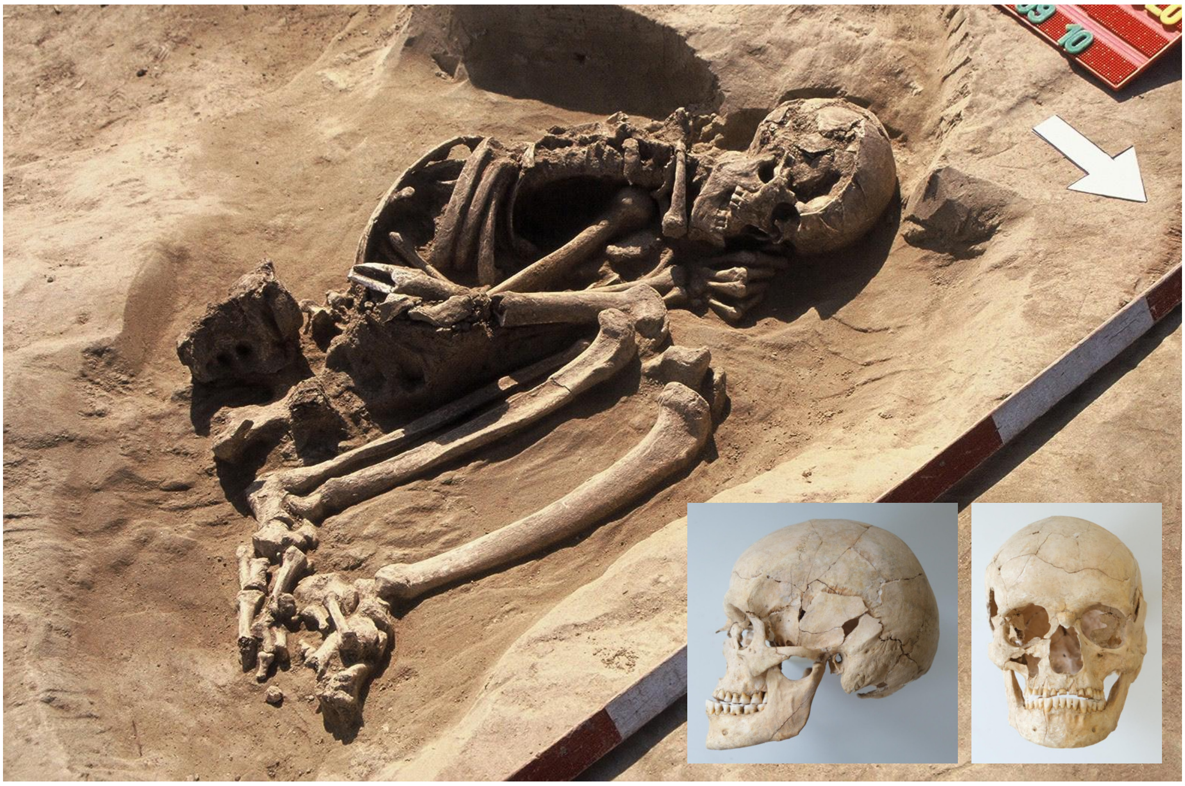

*Fig. S.1.5.11*

*Grave of individual S45. Moderately preserved remains of a 45 to 55 year-old male, oriented to W-E, next to a typical Kisapostag-associated pit, where broken pottery and a high amount of shell fragments were also recovered. This burial was located a significant distance from the other Kisapostag-associated graves. The individual had no grave goods, although shell fragments could be observed right next to the bones, and was buried in a heavily flexed position facing to the North. The remains were $^{14}C$ dated to 3702 ± 28 BP (2200-1980 cal BCE, 95.4% CI).Significant upper tooth abrasion can be observed with slight mandibular prognathism, likely in association with some sort of profession or behavioural anomaly that caused bruxism, for further information, see SI section 3.2.2.4. The skull is curvoccipital and mesochycranic, the mandibles are slightly protruding maybe in relation with the bruxism. Skull reconstruction and photo by Dániel Gerber, grave photo by Szilvia Fábián.*

#### 1.6) Transdanubian Encrusted Pottery culture

This archaeological culture, distributed in a large area of Western Hungary, is characterised by cremation burials with a large number of ornamentations highlighted by white encrustations on the dark (brown or black) surface of the pottery. The origin of this pottery was connected to the encrusted decoration of the Vučedol culture (Bóna, 1975, 1961). However, later studies emphasised the continuous development from the Kisapostag/Earliest Encrusted Pottery tradition (Hajdu et al., 2016; Kiss, 2012; Torma, 1978). Cremation burials of the older period can be dated between the $19^{th}$ and $17^{th}$ centuries BCE, while younger periods date between the $17^{th}$ and the $15^{th}$ centuries BCE (Kiss et al., 2019). Rescue excavations of the last decades resulted in the discovery of the large villages of this culture. These data suggest both smaller and larger, permanently occupied settlements with a relatively small population that repeatedly shifted their location. Besides single-layered sites, hilltop and (fewer) fortified settlements were also established. Local production of metallurgy is proven by a small number of moulds found in these settlements, as well as the specific bronze jewellery depositions of the Tolnanémedi hoard horizon (Kiss, 2020b, 2009). Urn burials and scattered cremation burials both occur, while inhumation is hardly ever documented (Kiss, 2012). However, these irregular settlement burials and several inhumations in 'urnfields' of the culture are important for the study of the anthropological and genetic makeup of the Middle Bronze Age population in western Hungary. During 2003 a round pit was excavated at the Balatonkeresztúr site (obj. B-938) in which eight human skeletons of three adults and five children were found without evidence of grave goods (Figure S.1.6.1). The feature was identified as a household refuse pit utilised for mortuary purposes dating to the Late Copper Age or the Early Bronze Age (Fábián, 2006; Köhler, 2006). Radiocarbon samples from the pit measured by the VERA AMS laboratory in Vienna, Austria yielded a date of 3455±35 BP (1890-1640 cal BCE, 95.4% CI); the HEKAL AMS laboratory in Debrecen, Hungary yielded a date of 3387±30 BP (1870-1540 cal BCE, 95.4% CI) which falls into the period of the Hungarian MBA. This multiple pit-burial, however, could be considered as an important funerary assemblage in the light of the recently heightened interest in irregular prehistoric burials (Earle et al., 2014; Müller-Scheeßel, 2013). The physical anthropological examinations of the Balatonkeresztúr remains prove the presence of the *planoccipital brachycranic taurid* type crania (Köhler, 2006), a physical trait which could be linked to the Bell Beaker group (K. Zoffmann, 2000). This outcome is significant considering that cremation was practised widely by Transdanubian populations of the period. Prior to the examination, virtually nothing was known about the physical anthropological makeup of these communities. Individuals S14 to S21 belong to this group: circles with solid lines represent first degree relationships, dashed lines represent second degree relationships (Figure S.1.6.2).

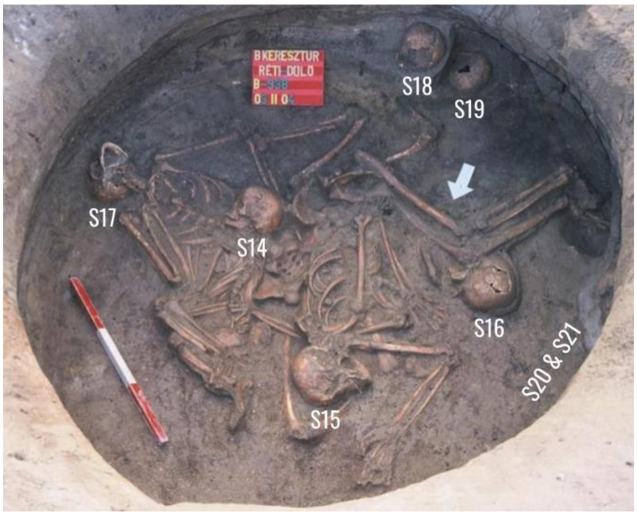


*Fig. S.1.6.1*

*Mass Grave B-938 of Bk-III. Photo by Szilvia Fábián.*

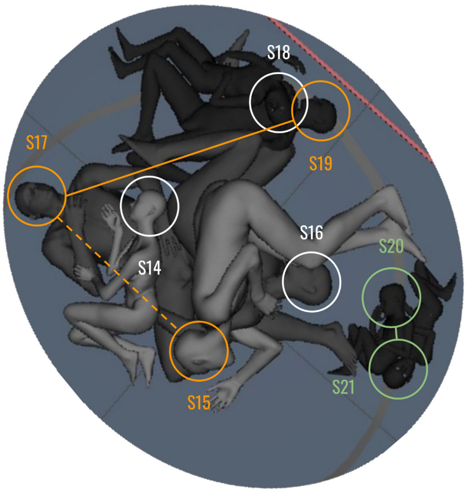


*Fig. S.1.6.2*

*Digital reconstruction of individuals from the mass grave of Bk-III. Connected individuals are related (solid line for first degree, dashed line for second degree).*

#### 1.7) Anthropological and paleopathological analyses

By Kitti Köhler, Balázs Gusztáv Mende

The condition of the studied anthropological material is medium, sometimes poor. The facial part could be reconstructed only in a few cases, while the postcranial remains are relatively well preserved. The finds are stored in the Rippl-Rónai Museum in Kaposvár. The estimation of age-at-death and the sex determination were based on the methods commonly used in physical anthropology (Acsádi and Nemeskéri, 1970; Bernert et al., 2007; Fazekas and Kósa, 1978; Ferembach et al., 1979; Işcan et al., 1985, 1984; Stloukal and Hanáková, 1978; Ubelaker, 1989). The paleopathological lesions were investigated macroscopically (Aufderheide and Rodríguez-Martín, 1998; Manchester, 1983; Ortner, 2003). The Bk-I EBA pit burial and Bk-II inhumation burials are first published here, while the physical anthropological analyses of S14 to S21 individuals from the Bk-III pit burial were already published (Köhler, 2006), and examinations are here completed by a detailed archaeological interpretation and results of isotope and aDNA testing. The results are summarised in Supplementary Table 1.

#### 1.8) Radiocarbon dates of the Balatonkeresztúr site

by Vajk Szeverényi, István Major & Mihály Molnár

Radiocarbon dating was performed for all Bronze Age horizons of the site. Where $^{14}C$ data are missing, DNA based biological relatedness supports the corresponding dating, see SI section 2.3. Overall 11 samples have been radiocarbon dated from the site: one sample was taken from the earliest Bronze Age burial connected to the period of Somogyvár-Vinkovci type material (S9), eight samples from Early Bronze Age graves with Kisapostag type material (four from Grave Group A (S1, S2, S6 and S7), two from Grave Group B (S11 and S13), and two from isolated graves (S10 and S45)), while two individuals were sampled from the Middle Bronze Age mass grave (S16 and S17).

All radiocarbon measurements were carried out at the HEKAL AMS C-14 facility of the Institute for Nuclear Research, Debrecen (Molnár et al., 2013b, 2013a), except for individual S16, which was measured at the VERA AMS facility in Vienna. The dates were calibrated with the ‘OxCal’ v4.4 software (Bronk Ramsey, 2009) using the IntCal20 Northern Hemisphere radiocarbon calibration curve (Reimer et al., 2020), see the individually calibrated dates at Supplementary Table 1. Individual S9 is dated to ca. 2560-2290 cal BCE (all calibrated dates are presented at two-sigma range, unless indicated otherwise). The dates from the graves of the BK-II phase range between 2200 and 1770 cal BCE, while BK-III dates fall between 1890 and 1540 cal BCE.

Bayesian analysis using the ‘OxCal’ software was implemented to attempt to obtain more precise age estimates. For the first model, independently identified typological phases were introduced as prior information. As a result of the analysis (Figure S.1.8.1), the ranges of the Somogyvár (2480-2200 cal BCE) and Encrusted Pottery dates changed somewhat (1890-1680 cal BCE and 1880-1610 cal BCE), moving the more probable part of the date ranges towards the Kisapostag dates. The latter did not change considerably as a result of the Bayesian analysis (2200-1780 cal BCE).

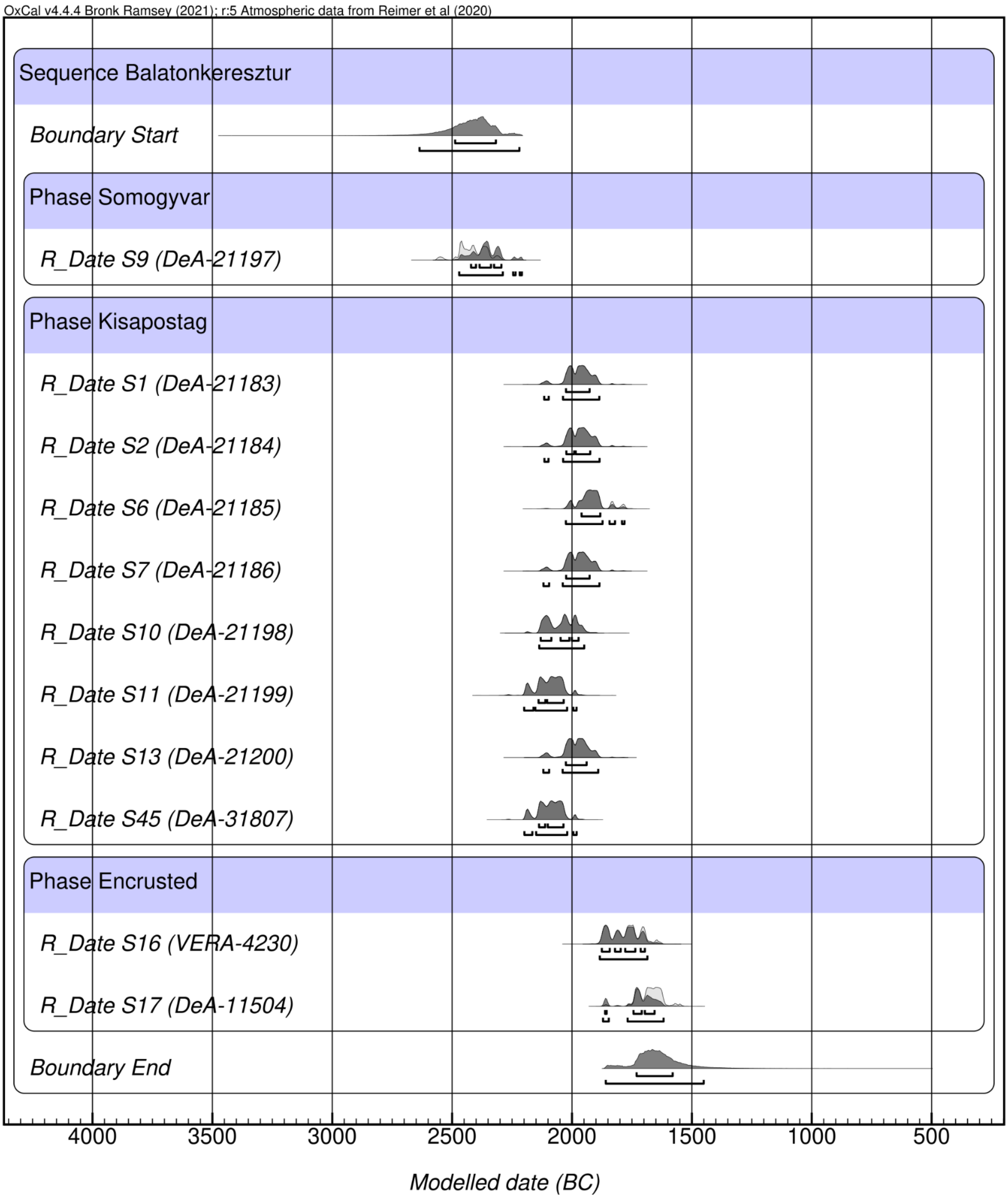


*Fig. S.1.8.1*

*Modelled radiocarbon dates of samples from the Balatonkeresztúr site, a multiplot for all three Bronze Age period (Bk-I-III) radiocarbon series.*

Only the Kisapostag dates were included in the second model as the contexts belong to a single phase. Grave Groups A and B and the isolated graves show no archaeological signs of chronological difference, thus were treated as the same phase. As a result (Figure S.1.8.2), the timespan of the BK-II phase could be reduced to ca. 2120-1900 cal BCE.

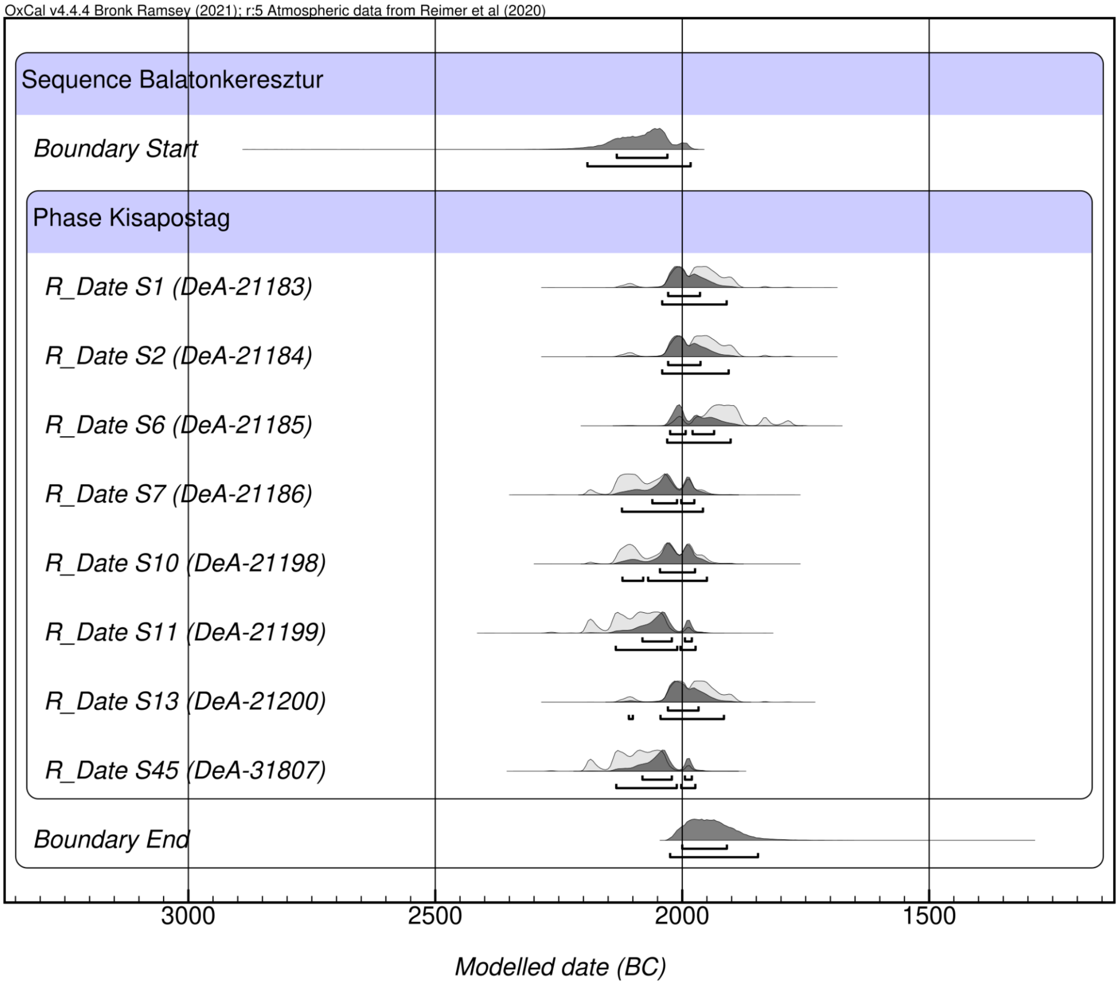


*Fig. S.1.8.2*

*Modelled radiocarbon dates of samples from the Balatonkeresztúr site, a multiplot for the Kisapostag period (BK-II) radiocarbon series.*

We carried out the same analysis, this time including only the graves from Grave Groups A and B, since they were both spatially bounded and genetically related as shown by the results of the archaeogenetic analyses (see SI section 2.3), thus they could be assumed to be temporally close as well. The result (Figure S.1.8.3) reduced the timespan of the use of the two grave groups to ca. 2130-1890 cal BCE.

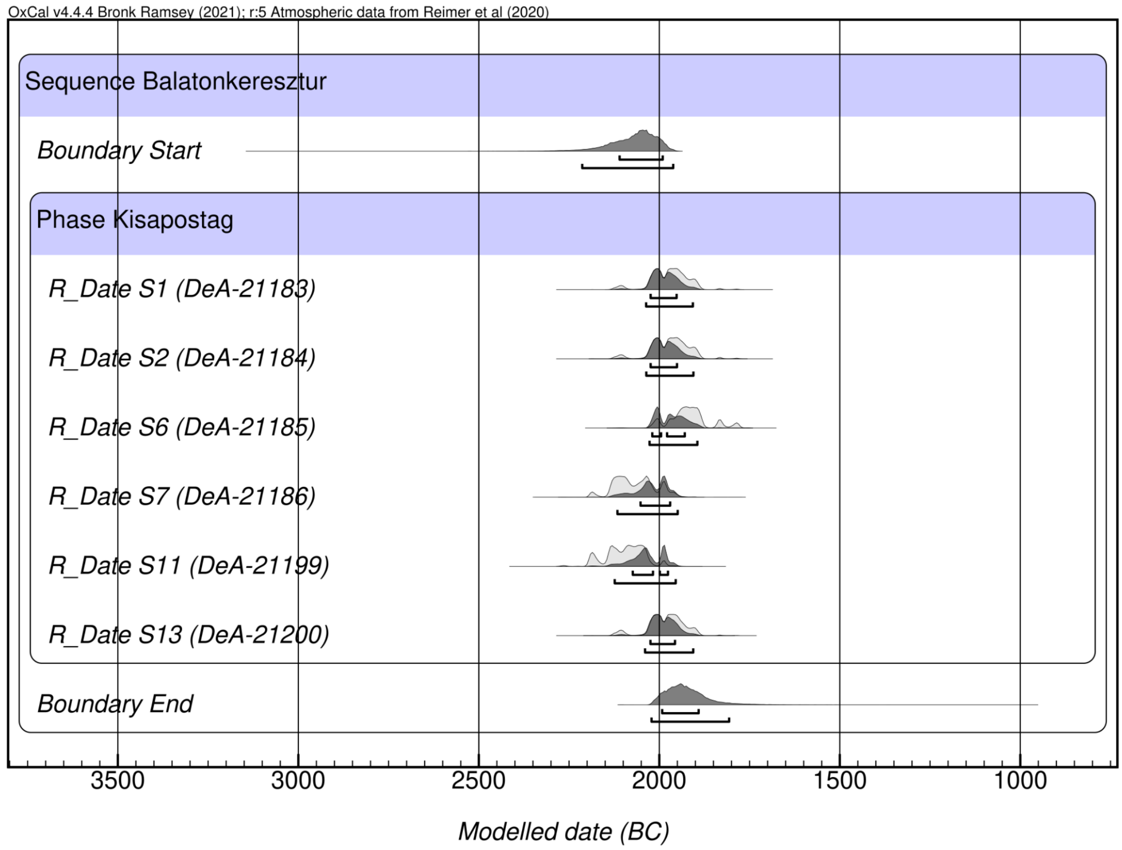


*Fig. S.1.8.3*

*Modelled radiocarbon dates of samples from the Balatonkeresztúr site, a multiplot for Grave Groups A and B.*

In recent years it has been recognized that archaeogenetic analysis and information on relatedness can provide additional chronological information that can be built into the Bayesian modelling of radiocarbon dates (Massy et al., 2022; Sedig et al., 2021). Sedig and colleagues even developed a new software, refinedate, to perform such analyses. Nevertheless, OxCal software is also perfectly capable of including and analysing such information and will be used here, following the work carried out by Massy et al. (2022). There is a first-degree (father-son) relationship between individual S11 from Grave Group B and individual S6 from Grave Group A. Since the father (S11) died at the age of 34-43, and the son at the age of ca. 17-18 years, it may be assumed that their dates of death were rather close, if not even simultaneous. Although we obviously cannot be certain of this, we combined these to date a single event. The combined date for their deaths (Figure S.1.8.4) is 2130-1940 cal BCE (95.4% CI), within this boundary dates between 2050 and 1940 cal BCE have a significantly greater probability (84.4% CI).

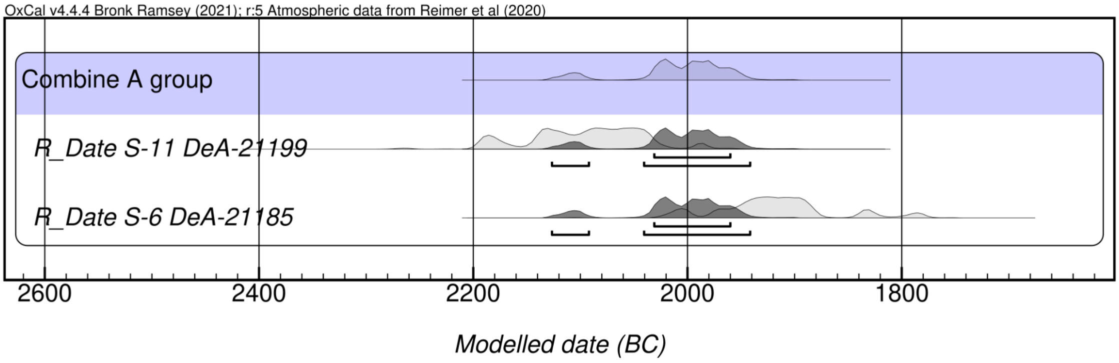


*Fig. S.1.8.4*

*Modelled radiocarbon dates of samples from the Balatonkeresztúr site, a multiplot for Graves 11 and 6*

In the case of a 17 years old son and a 43 years old father it is also possible that the son died earlier, which we may use as prior chronological information. Following (Massy et al., 2022) we can model this scenario as well. Since this prior information seems to reverse the order of these two dates (which is also indicated by OxCal through the poor agreement values), modelling them should actually place significant constraints on their ranges. We may model them in two ways. We may simply place these dates into a sequence where individual S6 precedes individual S11 (Figure S.1.8.5). In this case the two-sigma date for S6 is 2140-1960 cal BCE, where the period of 2040-1960 cal BCE has greater probability (68.9%) within. The date for S11 is 2110-1940 cal BCE.

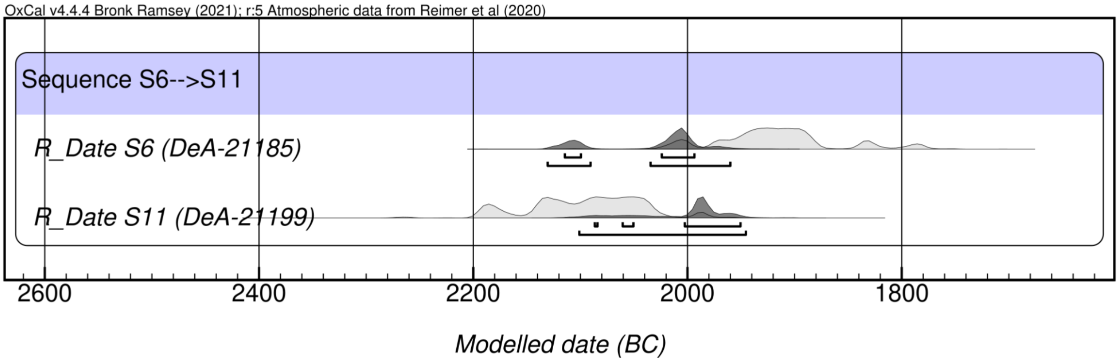


*Fig. S.1.8.5*

*Modelled radiocarbon dates of samples from the Balatonkeresztúr site, a multiplot for Graves 11 and 6, where the latter precedes the former.*

The other way is to place these two dates into a model with two phases, where the data of the son represents the earlier phase (generation), and that of the father the latter (Figure S.1.8.6). Here S6 is dated to 2130-1950 cal BCE, where the range between 2040 and 1950 has greater probability (73.5%) within, and S11 is dated to 2110-1940 cal BCE. Thus, both models indicate roughly the same dates for the deaths of these individuals.

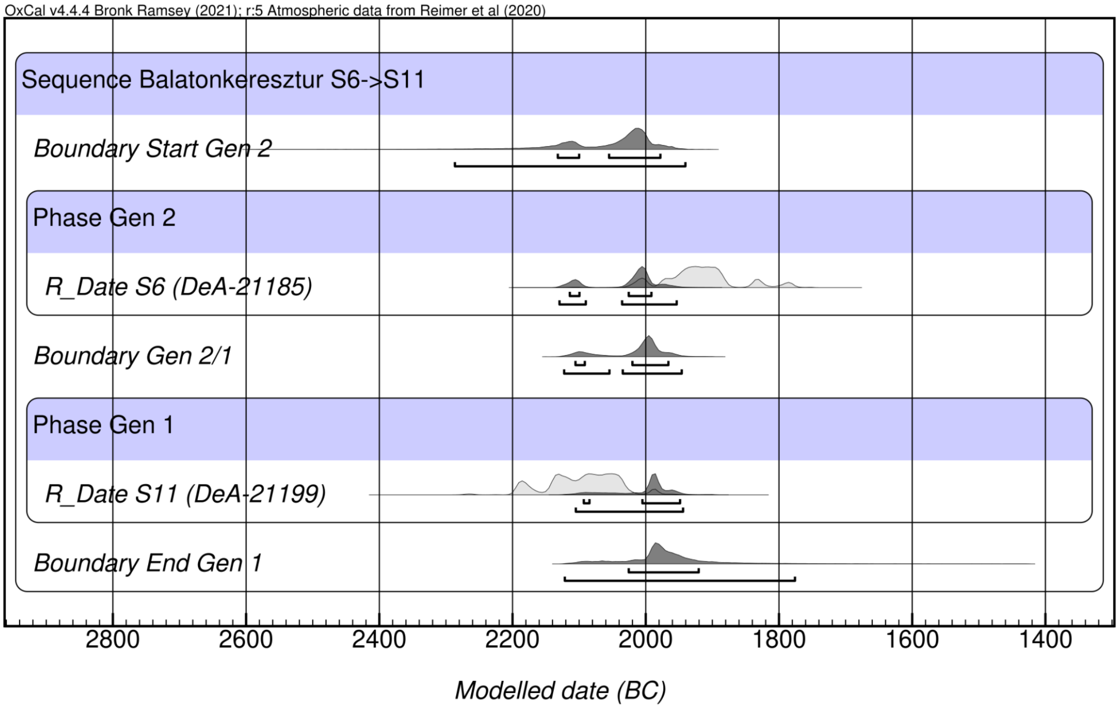


*Fig. S.1.8.6*

*Modelled radiocarbon dates of samples from the Balatonkeresztúr site for individuals S6 and S11.*

Finally, since the Middle Bronze Age samples derive from a mass grave in a pit, where the context and the full articulation of the skeletons indicate that the individuals deceased at the same time or within a short interval, the two dates were combined to date a single event. As a result (Figure S.1.8.7), the combined date for the mass grave is 1870-1620 cal BCE (95,4% CI), within this boundary the timespan between 1770 and 1620 cal BCE has a significantly greater probability (93,1% CI).

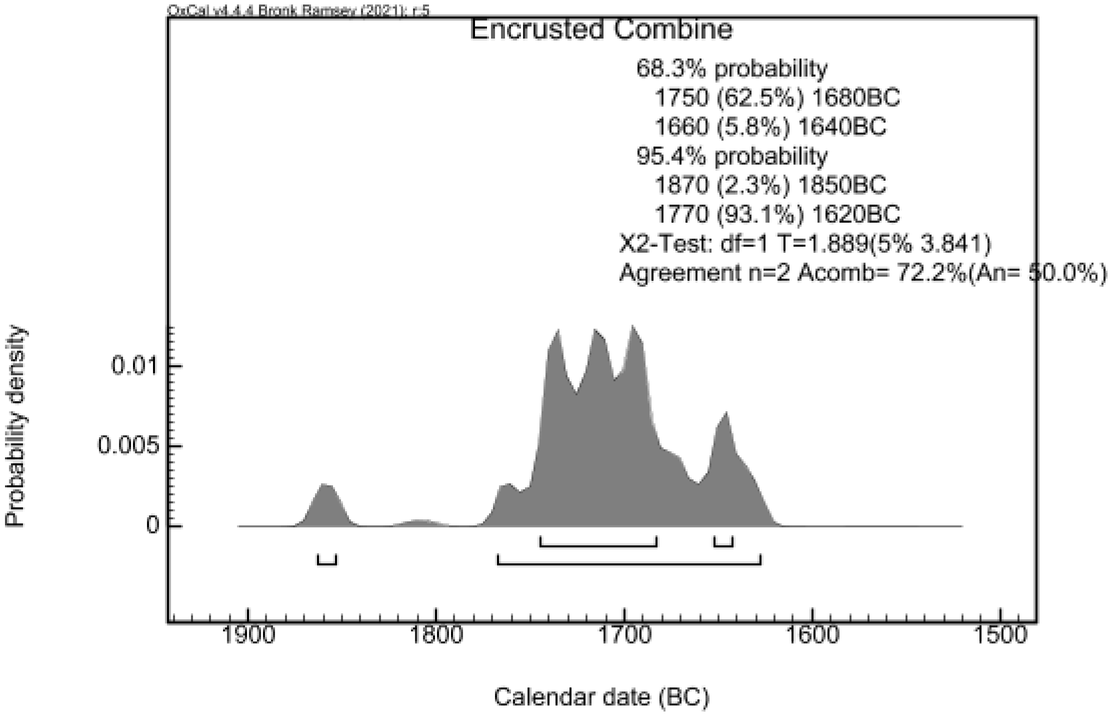


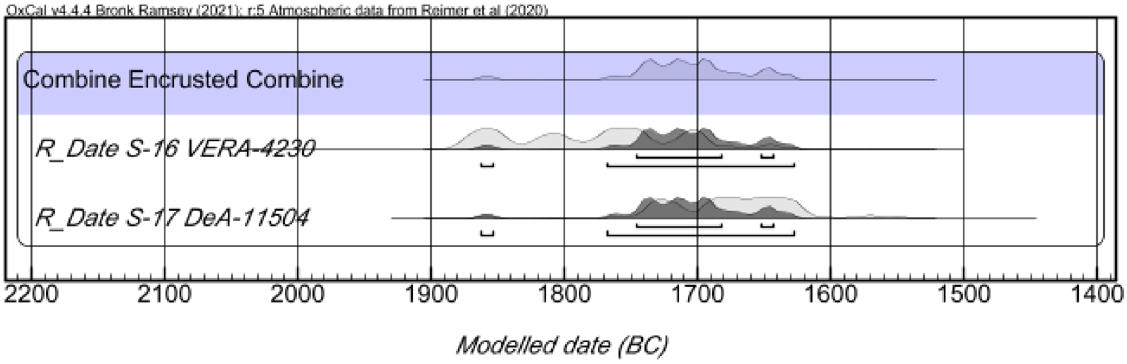


*Fig. S.1.8.7*

*Combined radiocarbon date of samples, from the Encrusted Pottery period (Bk-III) mass grave from the Balatonkeresztúr site.*

#### 1.9) Sr isotope data of Balatonkeresztúr site

by Julia I. Giblin, Anikó Horváth, László Palcsu

Samples from individuals S1-S13 were analysed for strontium isotopes ($^{87}Sr/^{86}Sr$) by the ICER laboratory, Atomki, Debrecen (Hungary), where the bone and enamel samples were physically pre-cleaned using a Dremel 300 drill equipped with a diamond tipped tool. Any remaining dentin on the teeth or cancellous bone on the cortical bone were removed using a diamond tipped Dremel tool attachment. Bones were repeatedly cleaned of dust by ultrasonication in milliQ water, after which they were rinsed in a 1 M acetic acid solution to remove diagenetic carbonate. Then, the samples were rinsed 3 times in milliQ water and dried overnight at 60°C. Five to-20 mg of enamel and 20 mg of bone samples, respectively, were weighed and placed into cleaned, labelled PFA beakers. Enamel samples were digested twice in 1ml of 14M $HNO_3$, bone sample was digested in 1 ml of 14M $HNO_3$

and 100 µl of $HClO_4$ and evaporated to dryness on a hotplate at 120°C. When the sample was fully demineralized, it was taken up in 8M $HNO_3$ and loaded into the pre-cleaned Sr-specific ion exchange resin column to separate Sr from the other matrix elements. Ultrapure water (UPW) (Merck, 18 MΩ·cm) and doubly distilled 14 M nitric acid were used for sample digestion, resin cleaning, and elution. Crown ether-based Sr-Spec Resin (100–150 µm particle size) from Triskem International, France was pre-cleaned with 8M HNO3 and UPW prior to use. Sr was isolated from the matrix components and from Rb to avoid an isobaric overlap of $^{87}Sr^+$ and $^{87}Rb^+$. Additional blank and standard solutions were included to verify the blank and accuracy of the chemical preparation. All sample preparation occurred in a Class 1000 cleanroom. Standard solution was prepared from the NBS987 $SrCO_3$ standard. Strontium isotope ratios were measured using a Neptune Plus MC-ICPMS (multi-collector inductively-coupled-plasma mass spectrometer, Thermo Scientific), equipped with an Aridus-3 (CETAC) desolvating system at the Isotope Climatology and Environmental Research Centre (ICER) in Debrecen, Hungary. The $^{87}Sr/^{86}Sr$ ratio was corrected for instrumental mass discrimination using $^{88}Sr/^{86}Sr$ = 8.375209, as well as by applying an interference correction for $^{87}Rb^+$ and $^{86}Kr^+$ with $^{85}Rb^+$ and $^{83}Kr^+$, respectively. All values were normalised to the reported value of 0.710240 for NIST SRM 987.

Dental enamel samples analysed by J. Giblin (S14-20) were isolated using a diamond drill bit attached to a Dremel tool and then chemically pretreated at the Center for Anthropological Research (CAR) at Quinnipiac University in a 2% bleach solution (NaOCl) to remove organic contaminants (13 hours), followed by a 0.1 M acetic acid ($CH_3CO_2H$) leach for 4 hours to remove non-structural carbonates. Samples were then freeze dried. Approximately five milligrams of each pretreated sample were weighed and digested in a Picotrace class ten clean room at the Yale Metal Geochemistry Center (Department of Geology and Geophysics at Yale University) by using in-house distilled ultra-pure acids. Each sample was dissolved in 1 ml 6.2 M hydrochloric acid (HCl) in 5 ml acid-cleaned Teflon beakers and put on a hot plate at 100°C overnight to digest. Two drops of pure hydrogen peroxide ($H_2O_2$) were added to dissolve potential organic matter and samples were then evaporated dry. Dried samples were then dissolved in 1 ml 2 M nitric acid ($HNO_3$). A split of this sample solution was purified for $^{87}Sr/^{86}Sr$ analysis using an ESI PrepFast-MC-Sr system. The resulting elution of strontium was evaporated on a hot plate and then raised in 1 ml weak nitric acid (5% $HNO_3$ v/v) for analysis on a Thermo Neptune MC-ICP-MS using NIST SRM 987 as a bracketing standard (average $^{87}Sr/^{86}Sr$ ratio of 0.71034±0.00001, 2SD). The data were corrected for mass bias using $^{88}Sr/^{86}Sr$ = 8.375209.

The radiogenic strontium isotope data ($^{87}Sr/^{86}Sr$) from all of the Balatonkeresztúr individuals are displayed in Figure 4 of the main text and SI Table 1. They were consistent with “local” values measured from plants, water and soil from the region to the south of Lake Balaton (Alt et al., 2014). The local estimate in this case was calculated by taking the mean value of the published plant and water samples from (Alt et al., 2014) plus or minus two standard deviations. There were no significant differences between males and females or subadults versus adults. The samples from the Transdanubian Encrusted Pottery culture (Pit: B-938) were slightly less radiogenic and more variable than the earlier time periods. This is particularly apparent for samples from three individuals (S15, S16, and S17) where the first and third molars were sampled. While both values could have come from the region south of Lake Balaton, they exhibit some spread, indicating movement within the region during early adolescence.

*SI Table 1*

*Radiogenic isotope data for Balatonkeresztúr site samples*

| Laboratory ID | Laboratory | Burial/Specimen | Tooth Sampled | $^{87}Sr/^{86}Sr$ |
|---|---|---|---|---|
| I/2584/2 | ICER Laboratory, Atomki | Balatonkeresztúr, S1 | Second molar | 0.709843 |
| I/2584/3 | | Balatonkeresztúr, S2 | Second molar | 0.709790 |
| I/2584/4 | | Balatonkeresztúr, S4 | Second molar | 0.709797 |
| I/2584/5 | | Balatonkeresztúr, S5 | Second molar | 0.709896 |
| I/2584/6 | | Balatonkeresztúr, S6 | First molar | 0.709778 |
| I/2584/7 | | Balatonkeresztúr, S7 | First molar | 0.709969 |
| I/2584/8 | | Balatonkeresztúr, S8 | Second molar | 0.709802 |
| I/2584/9 | | Balatonkeresztúr, S9 | Third molar | 0.709692 |
| I/2584/10 | | Balatonkeresztúr, S10 | Second molar | 0.709716 |
| I/2584/11 | | Balatonkeresztúr, S11 | Second molar | 0.709763 |
| I/2584/12 | | Balatonkeresztúr, S13 | Second molar | 0.709757 |
| CAR0995 | Quinnipiac University, Yale Metal Geochemistry Center | Balatonkeresztúr, S-14 (Pit B-938) | First molar | 0.709413 |
| CAR0991 | | Balatonkeresztúr, S-15 (Pit B-938) | First molar | 0.70956 |
| CAR1016 | | Balatonkeresztúr, S-15 (Pit B-938) | Third molar | 0.70872 |
| CAR0992 | | Balatonkeresztúr, S-16 (Pit B-938) | First molar | 0.70948 |
| CAR1015 | | Balatonkeresztúr, S-16 (Pit B-938) | Third molar | 0.70928 |
| CAR0993 | | Balatonkeresztúr, S-17 (Pit B-938) | First molar | 0.70953 |
| CAR1014 | | Balatonkeresztúr, S-17 (Pit B-938) | Third molar | 0.70895 |
| CAR0994 | | Balatonkeresztúr, S-18 (Pit B-938) | First molar | 0.70959 |
| CAR0990 | | Balatonkeresztúr, S-19 (Pit B-938) | First molar | 0.70961 |
| CAR0996 | | Balatonkeresztúr, S-20 (Pit B-938) | First molar | 0.709432 |

### 2) Uniparental genetics and relatedness

by Eszter Ari, Dániel Gerber, Bea Szeifert, Orsolya Székely

#### 2.1) The mitochondrial DNA haplogroups and their phylogenetic evaluation

Studying maternal lineages in the era of whole genome studies still has its potential for assessments of biological relatedness and preliminary evaluation of population connections. Our aim was not to recover individual histories and delicate connections, as the sparsity of the database in our view could distort such results, but to highlight general patterns that may or may not paralleled the autosomal results.

Here we retrieved mitochondrial DNA (mtDNA) haplogroups for all 20 individuals (Supplementary Table 1), and inferred phylogenies for each subgroup by using all available mtDNA sequences from modern and ancient databases (ENA and NCBI), see below. We used the Bayes method for inferring phylogenies of subhaplogroups (implemented in the 'MrBayes' v3.2.6 software (Ronquist and Huelsenbeck, 2003)). First, FASTA files were created from the BAM files by a custom pipeline implemented in the 'PAPline' using the 'Samtools' v1.6 (Li, 2011; Li et al., 2009) and the 'ANGSD' v0.931 software (Korneliussen et al., 2014) and a custom 'R' v3.4.4 script. The threshold for the coverage was setted to 2x< and 50%< (majority rule) allele occurrence. Sequence alignments using the 'MAFFT' v7.271 software (Katoh and Standley, 2013) were created using all publicly available recent (http://www.ianlogan.co.uk/) and archaic (AADR (Reich, 2021)) whole mtDNA sequences for each macrohaplogroup phylogeny, restricting the analysis to ~1500 samples per inferred subtree. Alignments were checked using the 'SeaView' software (Gouy et al., 2010) by eye and ambiguous alignment regions. Additionally, low coverage (>30% of N sites) samples were discarded. We used the '4by4' nucleotide model, substitution type of 2, site variation model of invgamma [+I +G], 4 chain, 1 million generations, disabled parsimony model and 0,25 burn-in fraction for the phylogenetic inferences by the 'MrBayes' software. The results are shown on Figs. S.2.1.1 -

S.2.1.10. We also restricted visualisation of subtrees where posterior probability (indicated with node numbers on the Bayesian trees) was higher than 0.6. Other phylogenies were considered unreliable or uninformative.

The individual BAD002 branches together with the Bell Beaker culture (BBC) associated individuals from Spain, which is in line with the Western European affinities of his genomic makeup (Figure S.2.1.1). Both in Bk-II and Bk-III maternal lineages various affinities appear, such as connections to the Balkan (haplogroup J2b1, individual S13), Scandinavia (haplogroup T2b, individual S19), Central Europe (haplogroup K1a4a1g, individual S10) or British Isles (haplogroup T1a4, individuals S5 and S6). Differences in geographical affinities clearly represent both changes and overlaps in genomic ancestry between Bk-II to Bk-III. One individual of the U4b1b1 subhaplogroup (ind. S15 from Bk-III) has direct maternal connections to Iron Gates HG groups (Mathieson et al., 2018) and to BDB001 from Germany (Rivollat et al., 2020), which we tested using the 'Beast' v1.10.4 software (Suchard et al., 2018) for divergence dates (Figure S.2.1.10), but this resulted in a rather uninformative tree that only confirmed the BDB001 connection. The maternal lineage of the individual S15 originated from the Mesolithic, thus its connection to Hunter Gatherer (HG) groups is rather too old to affirm inferences, while it shows a rough migration pattern from the Iron Gates to the Baltics. The mtDNA provides a weak signal for population continuity at the sequence level For example, only the H10a1 subhaplogroup shows direct phylogenetic connection between Bk-II and Bk-III, but considering the mitogenome sequence diversity (calculated by the R package 'Popgenome' (Pfeifer et al., 2014)) within periods (Bk-II *d*=0.978; Bk-III *d*=0.964), this is less surprising. Finally, individual S45 shows the only uniparental connection between the Jagodnjak site Encrusted Pottery group by haplogroup U5a1g (Figure S.2.1.10).

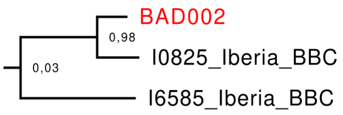


*Fig. S.2.1.1*

*Haplogroup K1a4a1 subtree, calculated by the MrBayes software.*

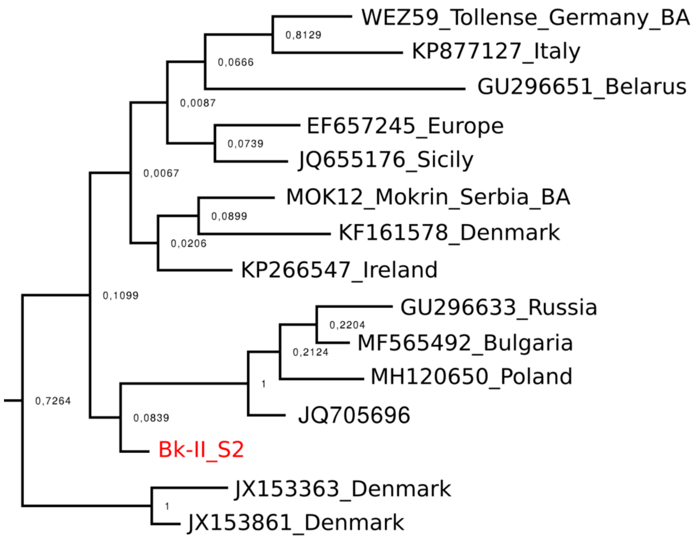


*Fig. S.2.1.2*

*Haplogroup U5a2b1a subtree, calculated by the MrBayes software.*

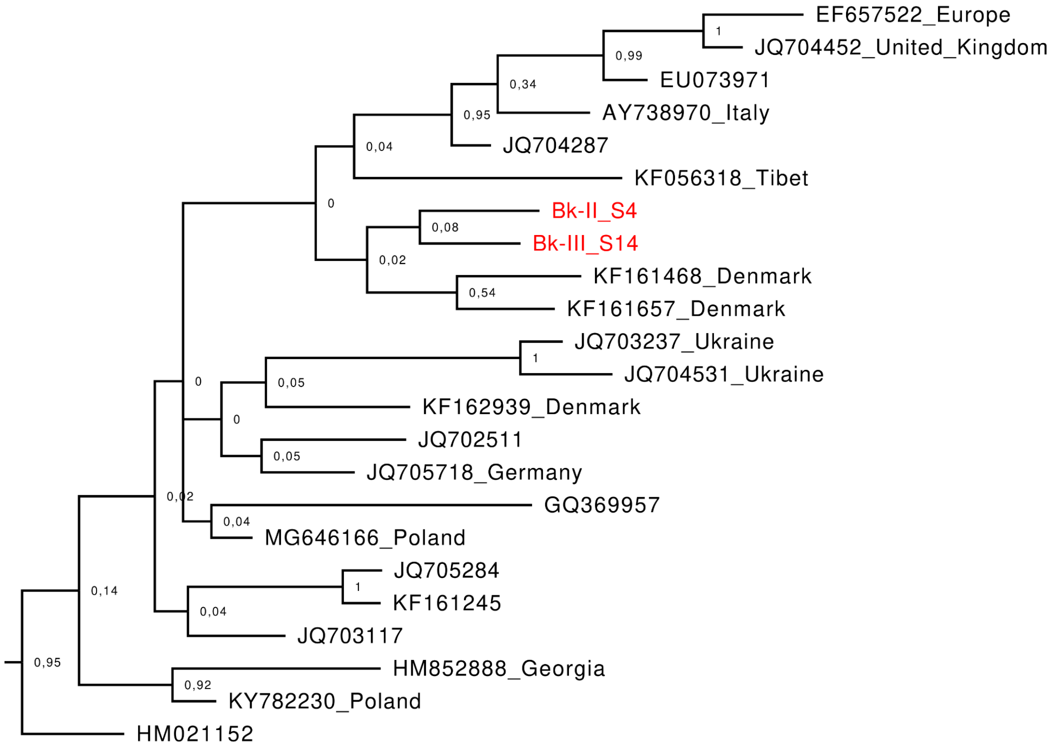


*Fig. S.2.1.3*

*Haplogroup H10a1 subtree, calculated by the MrBayes software.*

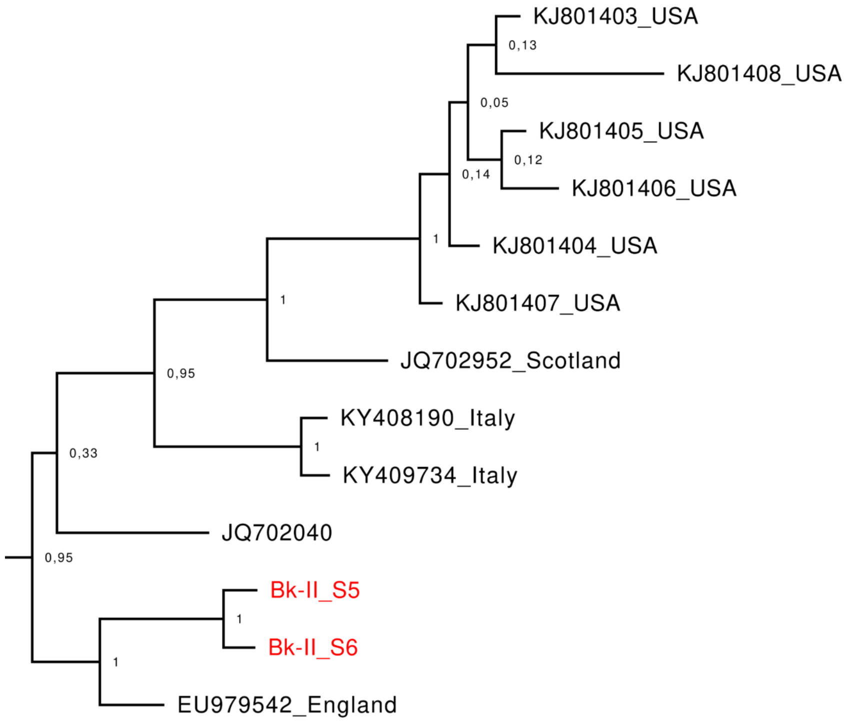


*Fig. S.2.1.4*

*Haplogroup T1a4 subtree, calculated by the MrBayes software.*

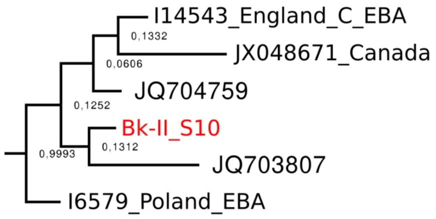


*Fig. S.2.1.5*

*Haplogroup K1a4a1g subtree, calculated by the MrBayes software.*

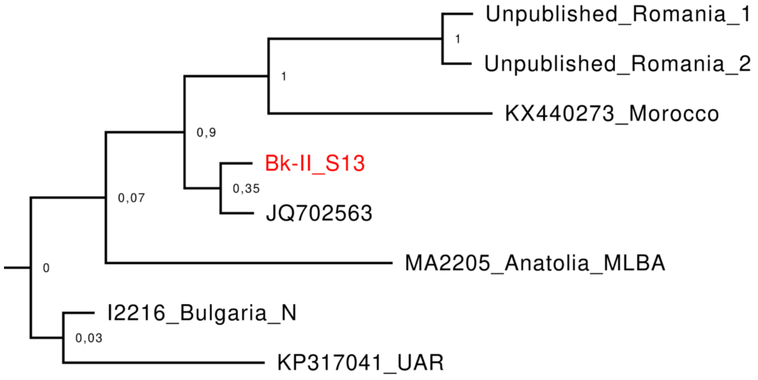


*Fig. S.2.1.6*

*Haplogroup J2b1 subtree, calculated by the MrBayes software.*

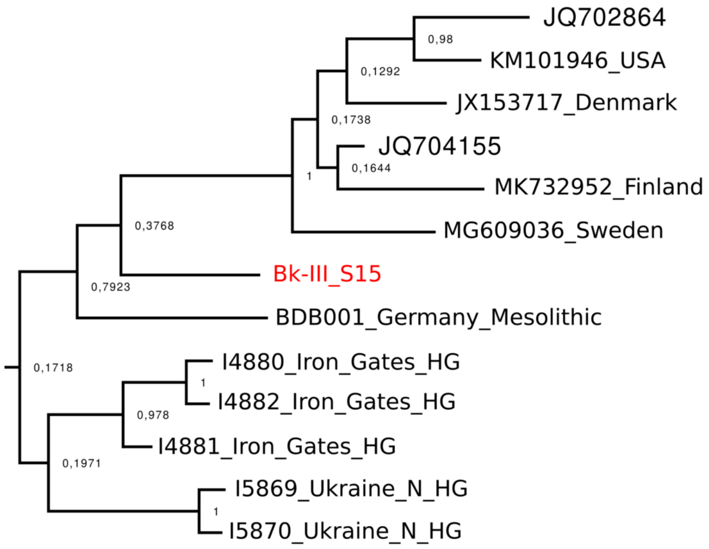


*Fig. S.2.1.7*

*Haplogroup U4b1b1 subtree, calculated by the MrBayes software.*

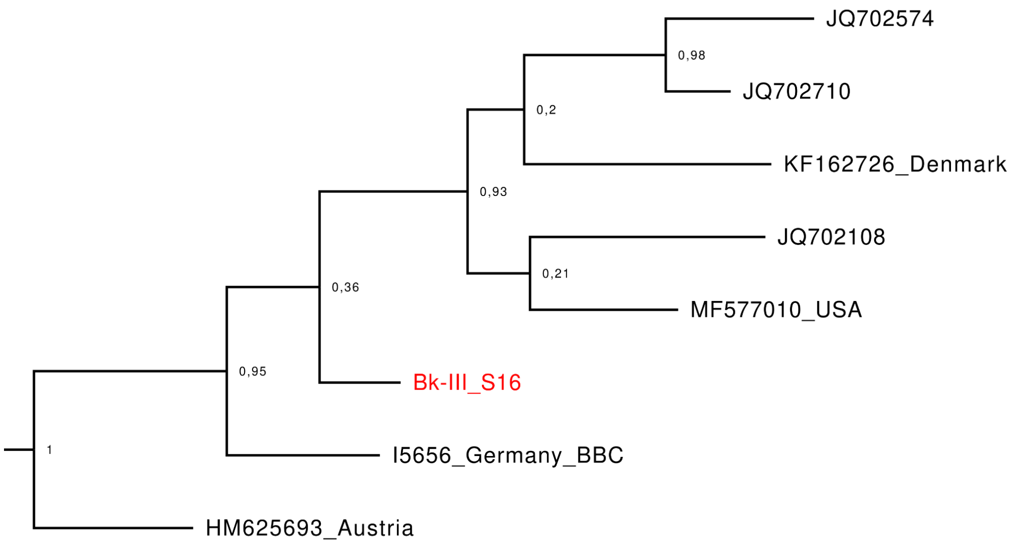


*Fig. S.2.1.8*

*Haplogroup T2g2 subtree, calculated by the MrBayes software.*

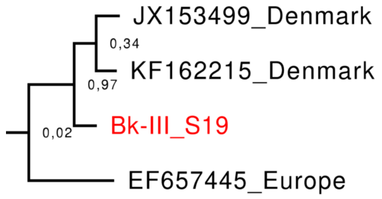


*Fig. S.2.1.9*

*Haplogroup T2b subtree, calculated by the MrBayes software.*

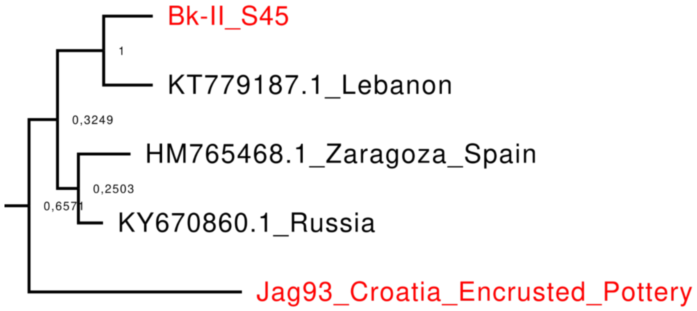


*Fig. S.2.1.10*

*Haplogroup U5a1g subtree, calculated by the MrBayes software.*

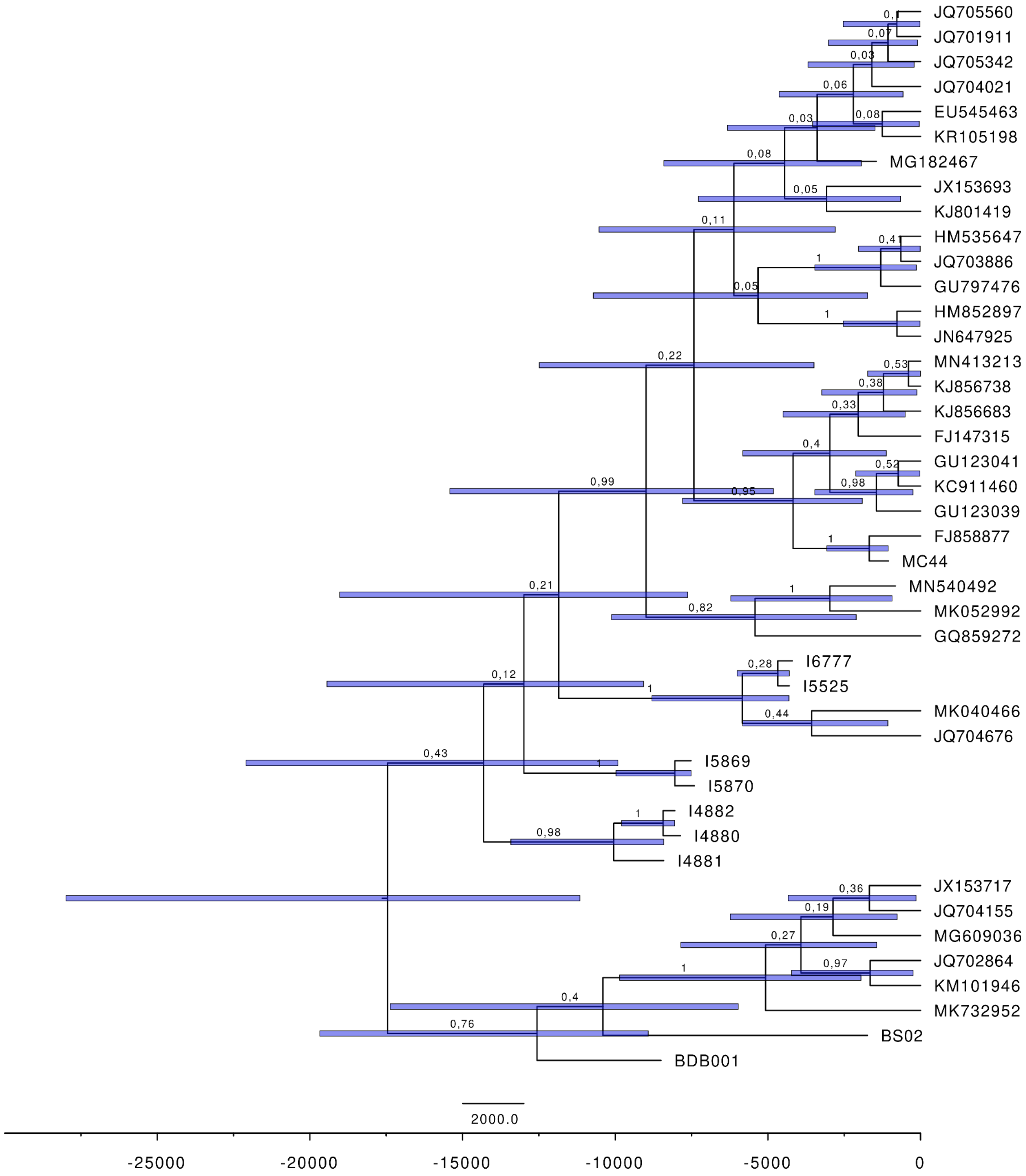


*Fig. S.2.1.11*

*Haplogroup U4b1b1 subtree, calculated by the BEAST software. Individual S15 diverged from the rest between ~4000-15000 BCE. Blue lines show the standard error of divergence dates.*

#### 2.2) Y chromosomal haplogroups and STR network analyses

For the Y chromosomal haplogroup determination we used a combination of shotgun, capture and 17 short tandem repeat (STR) markers from AmpFLSTR® Yfiler® PCR Amplification Kit (Applied Biosystems) analysed in 'NEVGEN' website (https://nevgen.org) (Supplementary Tables 1 and 3). Fine detailed classification was made by SNP set from capture and shotgun sequencing by using the 'Yleaf' v1 software (Ralf et al., 2018) with a custom set of SNP lists from ISOGG 14.255 ("ISOGG," 2019). We also made the Y STR network by using the 'Network' v10.1.0.0 and the 'Network publisher' v2.1.2.5 (Bandelt et al., 1999; “Network Software,” 2008) softwares, and a custom ‘R’ script that automatically pre-selected samples from available database data that are at either maximum 2 or 3 steps from the studied samples. We maximised 2 steps in cases where STR marker number is below 17, as these may include too many otherwise distantly related individuals to the network. Individual S9 from Bk-I has the R1a-Z280>>V2670 Y chromosome subgroup, which according to STR network analysis of modern samples is identical with a number of individuals from Northeastern Europe. Although the poor diversity pattern of this Y subgroup makes the results conditional. Contrary to maternal lineages, the paternal makeup for Bk-II shows high homogeneity with only a few STR marker differences observed, although a haplotype diversity test similar to mtDNA can not be performed due to the scarcity of data (Table 1 in the main text, Figures S.2.2.1/2/3). High Y homogeneity within and between the Bk groups shows Bk-II successiveness and population continuity in Bk-III, supported by qpAdm results as well (section 5.5.2.3) BK-III). It also suggests a strong patriarchal social network and – in combination with mtDNA data – likely a relatively smaller founder population. All lineages from BK-II and most lineages from Bk-III can be attested to I2a-M223>>L1229>>(Z2054), this subgroup was already present in Mesolithic/Neolithic France (Rivollat et al., 2020). I2a-L1229 is sparse in the database but group I2a-M223 were present in Megalithic cultures from the British Isles to today’s Czechia (Haak et al., 2015; Malmström et al., 2019; Sánchez-Quinto et al., 2019), reflecting the STR network analysis really well, and is also on par with the observations of (Rivollat et al., 2020). The qpAdm analyses (section 5.5.2.2) Bk-II) revealed that a major component of Bk-II is a Funnel Beaker culture (FBC) or related population. These groups frequently carried I2a-M223, pointing to a likely source for paternal lineages in Bk-II. The Jagodnjak group (Encrusted Pottery) from Croatia carried G2a2-Z31430 which can also be linked to these groups (Lipson et al., 2017), strengthening this theory. A two-step neighbour lineage from Early Mediaeval Hungary signals subsequent, probably limited, survival for these paternal lineages in Central Europe (Amorim et al., 2018). R1b-Z2103 is represented by two individuals in Bk-III, and it appears in contemporaneous populations such as in BBC period samples from Hungary (Olalde et al., 2018) or a Vučedol culture associated individual from Croatia (Mathieson et al., 2018). Sixteen marker STR network analysis shows more southeastern oriented connections for Bk-III R1b-Z2103 lineages, while the 17 STR marker network is more similar to I2a network distribution making results controversial (Figures S.2.2.1-3). Similarly to R1a, R1b is a relatively novel and widely distributed paternal lineage in the region which weakens its phylogeographic signal. Taking into consideration both previous appearances of this lineage in the region with the results of Network analysis, we can conclude that either it was present in Bk-II yet unsampled or came with Middle Bronze Age admixing groups. At this level both scenarios are plausible.

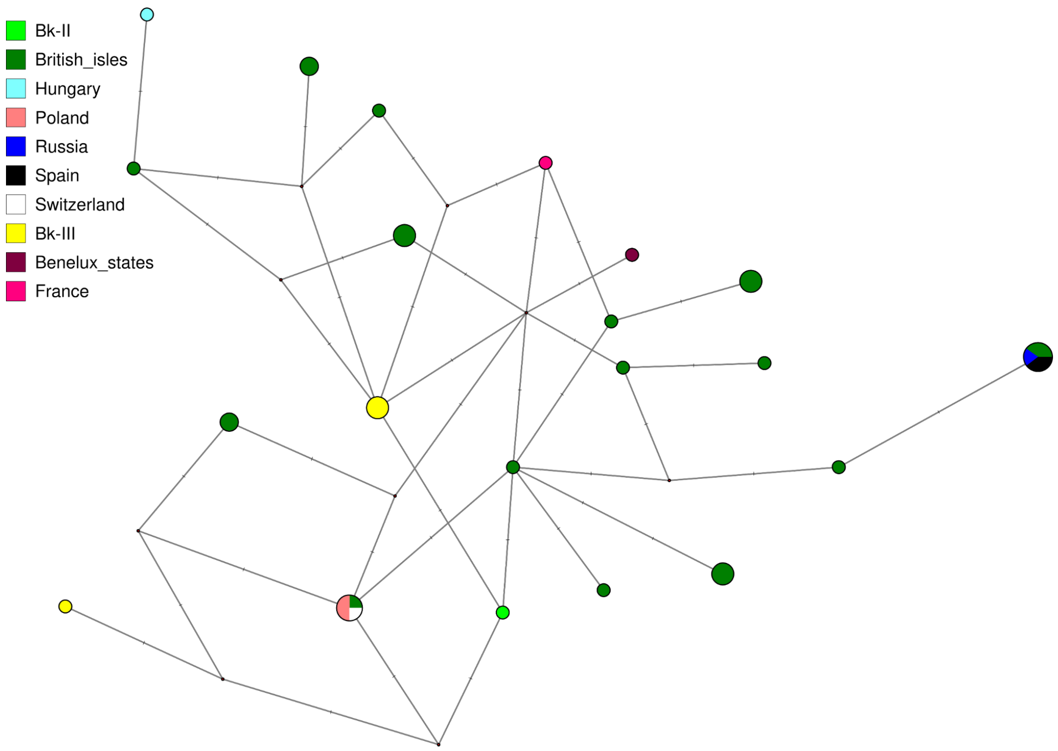


*Fig. S.2.2.1*

*Median Joining Network with 12 STR markers where sample subselection was maximised to two step distance among I2a-L1229 haplotypes. Affinities to the British Isles are apparent.*

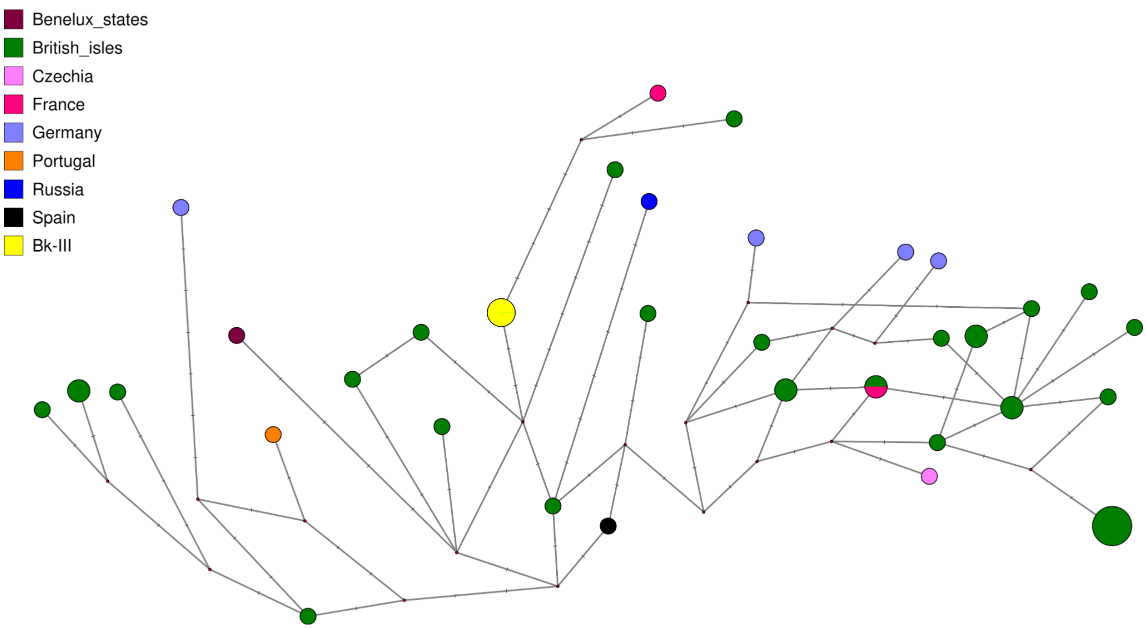


*Fig. S.2.2.2*

*Median Joining Network with 17 STR markers where sample subselection was maximised to three step distance among I2a-L1229 haplotypes. Population affinities are highly similar as seen on the 12 STR marker network.*

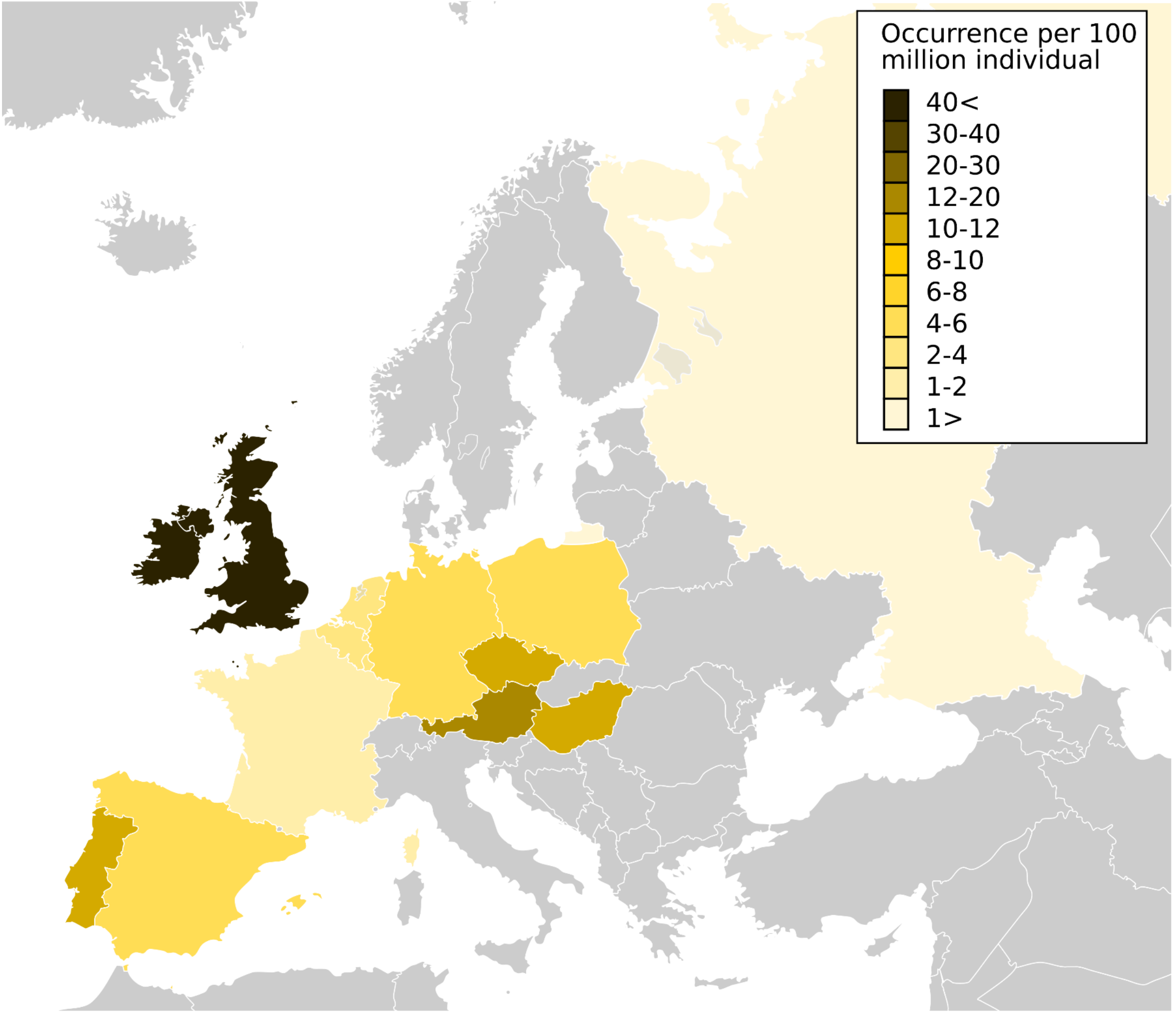


*Fig. S.2.2.3*

*I2a-L1229 Median Joining Network results from [Fig. S.2.2.1]() and [Fig. S.2.2.2]() are combined, from which we calculated sample number occurrences per 100 million individuals per country. This figure shows these proportional occurrences.*

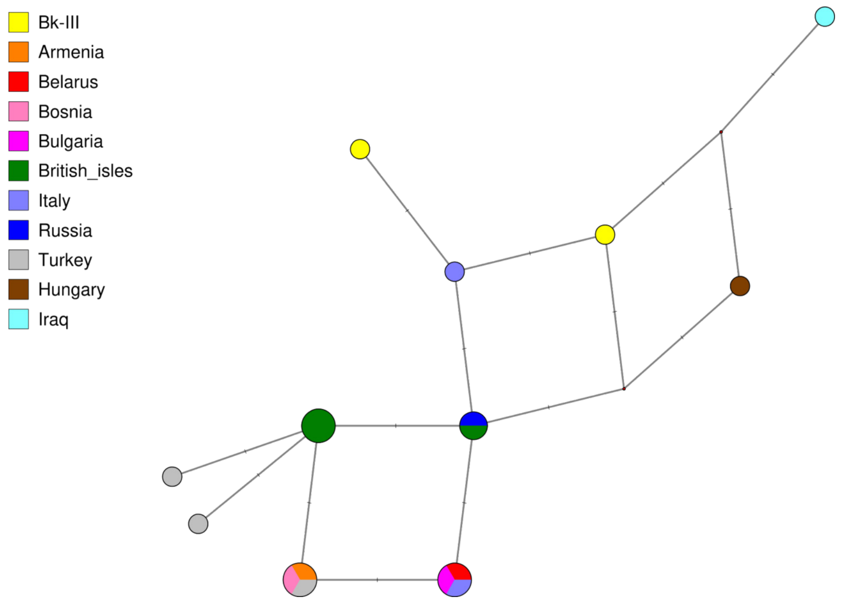


*Fig. S.2.2.4*

*Median Joining Network with 16 STR markers where sample subselection was maximised to two step distance among R1b-Z2103 haplotypes. More prominent southern affinities appear in line with modern and archaic geographical distribution of the lineage.*

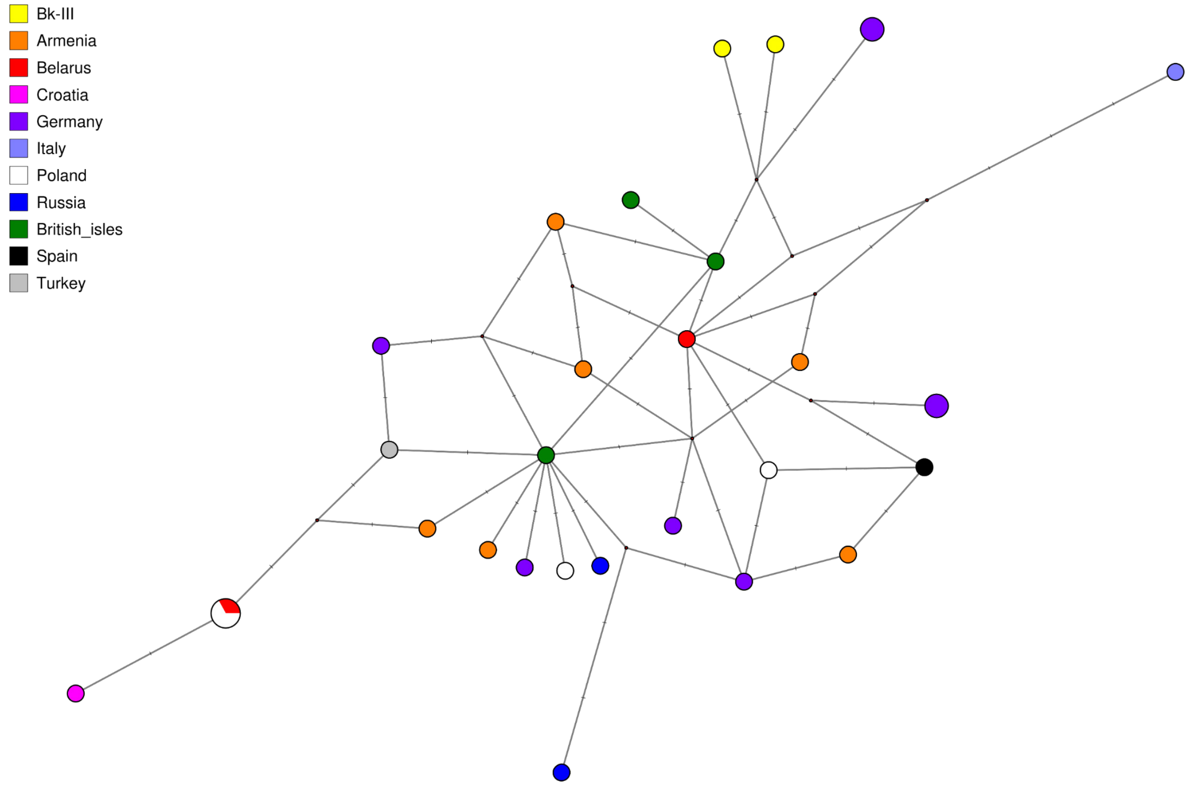


*Fig. S.2.2.5*

*Median Joining Network with 17 STR markers where sample subselection was maximised to three step distance among R1b-Z2103 haplotypes. Network shows more northern affinities for Bk-III, contradicting the 16 STR marker network.*

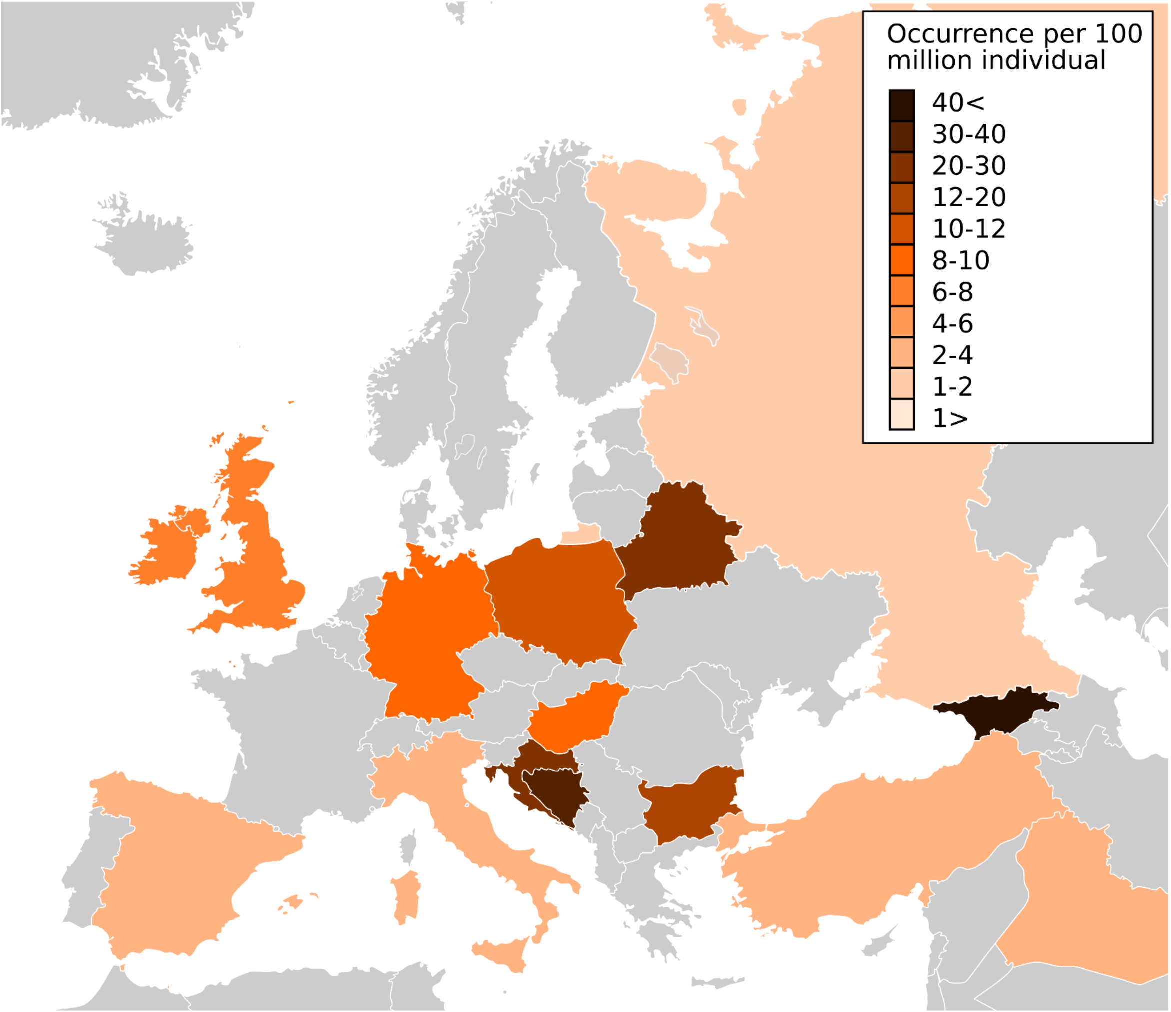


*Fig. S.2.2.6*

*R1b-Z2103 Median Joining Network results from Fig. S.2.2.4 and FIg. S.2.2.5 are combined, from which we calculated sample number occurrences per 100 million individuals per country. This figure shows these proportional occurrences.*

#### 2.3) Relatedness analysis

While it has been noted in recent publications that kinship relations are not necessarily determined through biological connection (Brück and Frieman, 2021; Mittnik et al., 2019), we still think that biological relatedness is an important source of information to be considered when reconstructing prehistoric social structures. Pairwise genetic relatedness assessments in this study were made by two independent methods due to low coverage genomes: (1) identity by descent (IBD) estimated by the 'READ' software (Monroy Kuhn et al., 2018) using mean values for normalisation, and (2) a new method called modified pairwise mismatch rate (MPMR), developed after (Allentoft et al., 2015). The formula for getting results per pair of individuals is the following:

$$MPMR_i = (M_i - Pop_{min}) \times (\frac{1}{1-Pop_{min}})$$

where $M_i$ is the number of matching SNPs between $i$ pairs of individual divided by the number of overlapping SNPs and where $Pop_{min}$ is the smallest $M$ value across the population of an arbitrary selection of individuals, in this case the whole set of burials from the Balatonkeresztúr site. This method theoretically would result in a ~40-50<% similarity rate for first degree, and ~35-40% similarity rate for second degree relationships. We ran this calculation with a bootstrap resampling of 1000 replicas, which we used to calculate the mean differences from the upper value of the standard error of the mean.

For each method we called genotypes from the 1240k SNP dataset (excluding mtDNA, X and Y chromosomes), and for the READ software run. We also added RISE479 (discovered at Érd, from the MBA period, Hungary) (Allentoft et al., 2015) to the pool, as this individual turned out to be highly similar to Bk-II and Bk-III, which provides sufficient background for IBD estimates. Supplementary Table 2a shows the results of the IBD and Table 2b the MPMR (Wang, 2017) patterns. Since we chose a rather conservative approach for the MPMR cutoff due to its higher sensitivity to first degree relations, we co-analysed relatedness network results of both methods (see Fig.1 in the main text), considering biological ages and uniparental genetic markers (see Table 1 in main text or Supplementary Table 1), for detailed results, see Supplementary Tables 2a and 2b.

As expected based on the chronological gap, individual S9 from Bk-I does not show genetic relationship to any individual from Bk-II nor Bk-III up to the second degree (uncle-nephew, half-sibling, grandparent-grandchild). Within Bk-II many relations were found: individual S4 is likely the son of S8; S11 is the father of S5 and S6; finally S1 and S2 are second degree relatives likely from the father's side. The structure of the grave groupings also suggest social links: the earliest members of the group are also the most genetically distant (*i.e.* highest HG component) as compared to others buried in Grave Group B within Bk-II, while their children and likely grandchildren and/or nephews are in Grave Group A. Bk-III shows extensive biological relatedness, however, it shall not be considered as one blood related family burial due to the structure of the relatedness network. In this case, individuals S17 and S19 are likely father-son relatives, while S15 and S17 are likely half brothers from the father's side. Additionally, individuals S20 and S21 are most likely a pair of dizygotic twins. Possible further connections may remain hidden due to low genomic coverages. Male dominance strongly influenced burial customs of the site both in Bk-II and Bk-III hinting at a special social structure compared to known Bronze Age groups, *e.g.* BA cemeteries from Germany (Mittnik et al., 2019) or from Serbia (Žegarac et al., 2021), where genetically confirmed male-female ratios are approximately even. Due to the chronological gap between Bk-II and Bk-III, second degree connections between them are improbable, thus we discarded these otherwise biologically possible connections in the final estimate, where we combined both 'READ' and 'MPMR' results. This is discussed in the main text.

### 3) Phenotype assessment

by Dániel Gerber

#### 3.1) Genetic sex and aneuploidy

Our new method that is implemented in the 'PAPline' pipeline is to determine genetic sex and the presence of any form of aneuploidies with low coverage data. First we obtained coverage data for all chromosomes for (1) the individuals in our dataset, (2) a selected ancient shotgun genome database (Allentoft et al., 2015), and (3) a mediaeval and modern day dataset (Csáky et al. *in prep*) by using the 'bedtools' v2.27.1 software (Quinlan and Hall, 2010). Then we analysed the results in a custom 'R' script. We used *Z*-score adjusted coverages – named as *ZAC* – for each chromosome by the following calculation:

$$ZAC_i = \frac{chr_{ij} - min_i}{max_i - min_i}$$

where *chr* is the coverage of the *i*-th chromosome of the *j*-th sample, *min* is the minimum coverage of the *i*-th chromosome across all samples, and *max* is the maximum coverage of the *i*-th chromosome across all samples. By providing the ratio of the Y and X chromosomes of these *AC*s, the presence of the Y chromosome can be confidently detected even at really low coverages. ZAC values are also eligible to test aneuploidy, as coverages correlate with ploidy. This method shrinks coverages to the same scale (except Y chromosome needs to be further divided by 2 as normally it exists in one or zero copy), from which we can easily infer ploidy even as low as 0.01x genomic coverage.

Accordingly, individual S10 possesses an extra copy of the Y chromosome, an aneuploidy known as Jacob's syndrome. It is important to note that data type (shotgun *vs.* capture), mapping method, and post filter processes may significantly alter output if one compares different datasets to each other, thus we recommend to analyse similarly processed data with this method.

#### 3.2) Variant discovery

##### 3.2.1) SNPs of pigmentation

For pigmentation assessment, we used a custom set of phenotypically relevant SNPs listed in the HIrisPlex-S system (Chaitanya et al., 2018), which we extended with variants from the SNPedia database (Cariaso and Lennon, 2012), for results, see Supplementary Table 5. SNPs were called with the following filters: trimmed read ends (2 bp), BaseQuality>30, and MappingQuality>25. According to the available data, BAD002, despite providing weak signals for pigmentation, likely blends to the average Neolithic European variation. S9 from Bk-I had European light skin that probably had no freckles and was likely a bit less sensitive to sunburn. The most likely colouration for his hair is blonde and for his eyes is blue, these traits are in accordance with previous studies of people with steppe origin (Reich, 2018). Bk-II shows almost uniform makeup. Even with sparse data, only a couple of variants show heterozygosity within the population. Accordingly, the overall pigmentation was dark, with skin colour probably darker than today's average European. However, light and blue eye colouration did occur, thus most individuals probably had lighter hazel or green eyes. The addition of other SNPs associated with light pigmentation, freckles and blonde hair, may have also played a role in their appearance. The individuals in the Bk-II population were probably more sensitive to sunburn and had low tan response. Surprisingly, Bk-III shows a high variability of pigmentation patterns. The colouration of the preceding population is present, but more and well-pronounced lighter pigmentation patterns are also observed. Two individuals (S14 and S17) from the mass grave probably had reddish blonde hair, green eyes and pale skin, maybe even freckles, likely as a result of admixture from populations of various origin.

##### 3.2.2) SNPs of clinical significance

We collected a custom set of variants from the ClinVar database (Landrum et al., 2018), which we tested on the new data. Genotype likelihood (GL) values and coverage data were calculated using the 'ANGSD' v0.931 software (Korneliussen et al., 2014). Since low coverage and deamination could highly bias the results, we only considered variants for discussion that are either (1) transversions, (2) have at least 3x coverage by MAPQ>25 and BQ>30 read positions, and (3) the skeletal remains show signs of symptom onset. The final estimates were summarised in Supplementary Table 6.

##### 3.2.2.1) Lig4 syndrome

The individual S15 carried a variant (rs104894421, GL>0.99) in heterozygous form which potentially causes Lig4 syndrome when carried in homozygous form. Therefore, in theory the genotype of individual S15 did not result in the actual onset of symptoms. On the other hand, in the Jagodnjak group the very same variant appears in individual JAG93 (1x coverage), while it is also present in individual JAG82 but on another locus (rs104894420) of the same gene. While it is merely speculative since individual S15 is not covered in the latter locus, there is a possibility that these two SNPs were both present in the Encrusted Pottery culture associated population, and individual S15 inherited two different loci both causing frameshift mutation resulting in an actual onset of symptoms, supported by some of his skeletal features, such as elongated face and hip dysplasia (Altmann and Gennery, 2016).

##### 3.2.2.2) Diabetes and related disease susceptibility

Individuals BAD002, S1, S8, S9, S10, S11 and S19, S20 and S45 carried variants for Type 2 diabetes susceptibility, diabetes 2 driven microvascular complications and obesity. Unsurprisingly, the discovered SNPs are also highly prevalent in today's European populations (~30-50%)("ClinVar," 2021).

##### 3.2.2.3) Hereditary spastic paraplegia

rs121434442 is an autosomal dominant SNP in the KIF5A gene that causes hereditary spastic paraplegia, a neurological disorder causing progressive muscle stiffness in the limbs, taking onset anytime between 2 to 40 years of age. Complex forms cause a high variety and number of symptoms, while the condition itself ranges between low-key to serious disability. Another SNP in SPG11 gene is also associated with the onset of symptoms. Although variants in both genes are transitions, S6 may have carried these, as individual S11, father of S6, shows signs of a limb condition that could be linked to the years-long onset of symptoms (Shribman et al., 2019).

##### 3.2.2.4) Autism 15 (AUTS15) susceptibility

Transversional mutation in rs7794745 can be observed in individuals S6 and S45, which SNP is known as a contributing factor for autism spectrum disorder (AUTS15), whereas the affected CNTNAP2 gene is linked to voice-specific frontotemporal activity (Koeda et al., 2015). S45 shows signs of severe bruxism (Figure S.3.2.2.4), a common phenomenon among autistic children (Muthu and Prathibha, 2008). While this dental condition could be the result of some profession-related abrasion, the co-occurrence of these features suggests the actual onset of symptoms. CNTNAP2 is also affected by rs2710102, found in individuals S9 and S19, but as a low coverage transitional mutation its actual prevalence is questionable, especially in the light of population discontinuity between Bk-I and Bk-II/III.

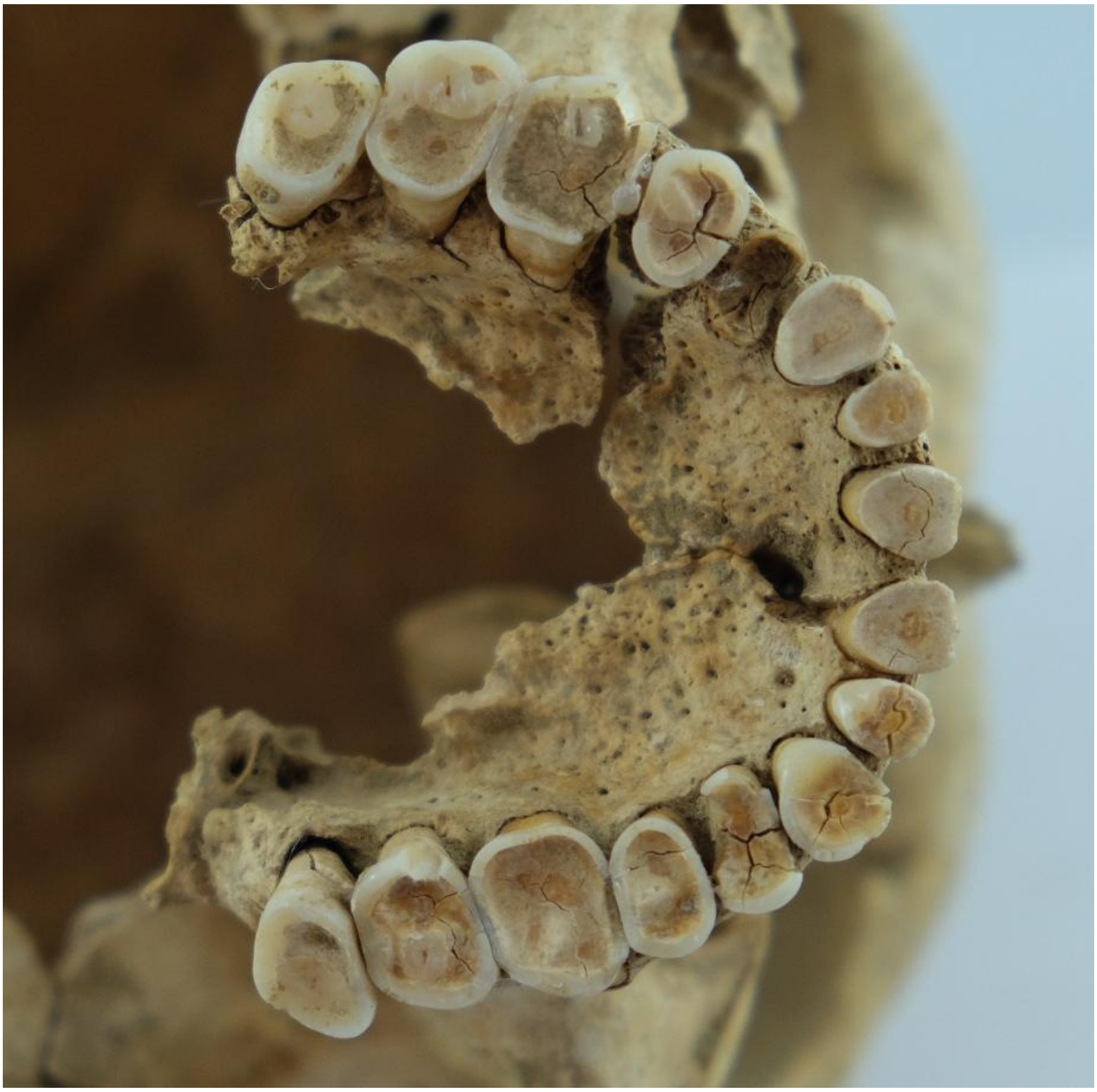

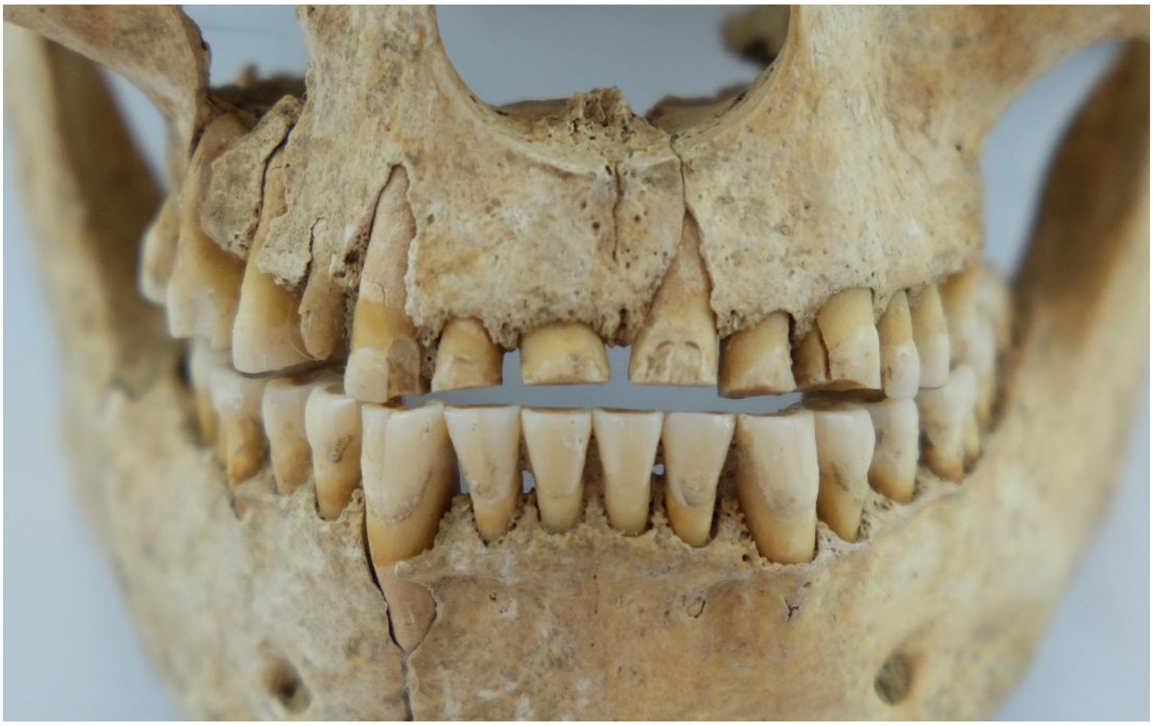

*Fig. S.3.2.2.4.*

*Upper photo is the occlusal view of the maxillary teeth of individual S45. Lower photo shows the frontal view of the teeth, the severe wear is only restricted to the upper incisors. Photo by Dániel Gerber.*

### 4) Facial reconstruction

by Ágnes Kustár

The method of facial reconstruction can be used to visualise the appearance of past individuals, which is also frequently used today for forensic identification. The skull of individual S13 is well preserved, except for the right zygomatic and temporal bones, which were slightly damaged. According to previous evaluation by Kitti Köhler, the skeletal remains suggest a gracile and short (~154 cm) physique without any traces of perimortem injuries. Secondary analysis by Ágnes Kustár confirmed these results, and also observed that there were no traces for - at least intensive - childbirth. There were two small and shallow perpendicular cuts on one of the ribs that could be perimortem damage, although neither the surrounding area nor the placing of the cuts suggest a life-threatening injury.

The physical appearance of the skull reflects the past facial features. Absolute size and form is small and gracile, overall highly feminine in characteristics. The shape of the skull is high and short, the forehead is narrow and bulging. The nape is curved, muscular joints (*linea nuchae superior et suprema*) are prominent, the external occipital protuberance is well developed, and while the mastoid is small the features indicate strong neck muscles. The nasal cavity is moderately wide (*mesorrhine*) and its lower margin is sharp (*anthropine*), both indicate moderately wide nasal wings. The nasal root is shallow and the nasal bridge is narrow. Distal parts of the nasal bones are broken and were reconstructed with resin. The anterior nasal spine is moderately developed and slightly turns upward. Each feature combined with moderately protruding nasal bones from the facial plane suggest a moderately prominent cartilaginous nasal structure (*nasus externus*). The orbit is high (*hypsikonch*), its shape is rounded, and the upper margin is slightly reclined. The distance between orbits is relatively small. The zygomatic bones are smooth, short and slim, the canine fossa is shallow. Due to the prominent protrusion of both jawbones, the upper and lower incisors are highly protruding. Accordingly, the lips may have been moderately full and also protruding. The mandible is small and short, and is moderately thick in its body. The *ramus mandibularis* is short, the bone joint is small, but the jaw corner is almost rectangular with moderately developed muscular joints. The chin bone is sharp and slightly protruding.

For facial reconstruction a detailed copy of the skull was made by rapid prototyping technique. First, the skull was scanned by Computer Tomography at Huniko Egészségügyi Szolgáltató Kft. (Hungary), then a resin cast was made by selective laser sintering at Varinex Zrt. (Hungary). Finally, a plaster cast was made after filling the gaps in the skull copy with beeswax (Figure S.4.1). During the process, the plaster cast was used as a base where soft tissues were built back for achieving facial features. The reconstruction was performed by well defined artistic and anatomical methodology (Gerasimov, 1949; Prag and Neave, 1997; Taylor, 2001). Plasticine clay was used to form muscles on the skull (Kustár and Skultéty, 1996; Sjøvold, 1981). The thickness of the muscles were estimated by the roughness of the bone surfaces (Röhrer-Ertl and Helmer, 1984) using 45 measuring points (Figure S.4.2). Markers of tissue thickness were placed on the plaster skull to measure points, lengths of the marker spikes were determined according to average values listed in SI Table 2. Long needles were used to mark regions and borders on the skull which became covered during the process to keep monitoring important morphological points (Figure S.4.3). Eyeballs made of resin were measured to be 25 mm in diameter fitting eye sockets. The cartilaginous nasal septum (*septum nasi cartilagineum*) was made of hardened beeswax to preserve shape, while the nasal spine and nasal wings were made of plasticine. The shape and dimensions of the nasal structure was estimated based on the shape of the nasal bones, the proportions of the nasal cavity and the direction of the anterior nasal spine (Rynn et al., 2010). Shape, size and length of mimic muscles were estimated according to the imprints of muscular joints on bone surface, anatomical regularities and individual characteristics. In the final artistic phase harmonisation of the features was performed (Figure S.4.4).

*SI Table 2*

*Grades according to bone relief: 1: Very gracile, smooth, 2: Less gracile, a little rough, 3: Rough, 4: Robust, very rough*

| Soft tissue thickness data on the face of individual S13 | | |
|---|---|---|
| **Measuring point** | **Degree** | **Thickness (mm)** |
| Bregma (b) | 1 | 4 |
| Metopion (m) | 1 | 4 |
| Glabella (g) | 1 | 5 |
| Nasion (n) | 1 | 4 |
| Rhinion (rhi) | 1 | 2 |
| Philtrum (ph) | 1 | 7 |
| Labiomentale (lab) | 1 | 7 |
| Pogonion (pog) | 1 | 8 |
| Gnathion (gn) | 1 | 7 |
| Arcus sup.medialis (acm) | 1 | 7 |
| Arcus sup.lateralis (acl) | 1 | 4 |
| Ectoconchion (ek) | 1 | 3 |
| Orbitale (or) | 1 | 3 |
| Dacryon (da) | 1 | 2 |
| Lacrimale (la) | 1 | 2 |
| Lat.apertura pir. (lat.ap) | 1 | 2 |
| Alare (al) | 1 | 3 |
| Subspinale lat. (ss lat) | 1 | 9 |
| Caput mandibulae (cap) | 1 | 3 |
| Gonion (go) | 2 | 4 |
| Zygion (zyg) | 1 | 2 |
| Facies zygomaticus (fac.zyg) | 1 | 4 |
| Zygomaxillare (zm) | 1 | 3 |
| Proc.mastoideus (mast) | 1 | 3 |
| Lambda (l) | 2 | 5 |
| Opisthocranion (op) | 2 | 5 |
| Subnasale (sn) (H11)* | NA | 13 |
| Labrale superius (ls)(H12)* | NA | 11 |
| Labrale inferius (li)(H13)* | NA | 12 |
| Mid mandibular border (mmb)(H28)* | NA | 11,5 |
| Euryon (eu)(H29)* | NA | 5,5 |

**H11-H29 measurements according to Helmer (Röhrer-Ertl and Helmer, 1984) (30-39 year old women)*

The reconstructed face reflects the shape of the skull: the head is wide and short, the forehead is narrow and convex, the *glabella* and eyebrows are less protruding and is slightly arched. The face itself is moderately wide and it narrows towards the chin, while her neck is relatively robust. The nasal root is moderately deep, the nasal bridge is also moderately protruding and narrow. The tip of the nose is sharp and is in a slight upward position. In the frontal view the nasal root is narrow, the nasal bridge and nasal wings are moderately wide. According to the small protuberances in the margins of the orbit (as indicators for the joint points of the eyelid hanging fibres), the eye openings are completely horizontal. The eyes are slightly sitting close to each other and the eyelid fold is moderate. The mouth is moderately wide and thick, the incisors are protruding and the protruding lips are not closing entirely. The jawline is not prominent and the chin is slightly protruding. The ears can not be confidently reconstructed, thus they were adjusted in harmony to other facial features, especially to the nose and eyebrows. Pictures were taken of the clay model in frontal (Figure S.4.5) and profile (Figure S.4.6) view.

Nutrition information can not be obtained from osteological features, thus we suggested average condition in facial reconstruction. Individual S13 died between 35-45 years of age, hence we created mimic wrinkles specific to 40-60 years olds. Colouration was created according to genetic results (Supplementary Table 5, section '3.2.1) SNPs of pigmentation') by Zsuzsa Herceg (Figure S.4.7). The overall pigmentation was transitional with intermediate skin tone and hazel eyes with blondish-brown hair accompanied by the presence of freckles. Hairstyle and hair structure was created according to genetic and artistic evidence, as Encrusted Pottery figurines show straight, braided hair (Figure S.4.6 (Hajdu et al., 2016)). Additionally, digital grave reconstruction was made by Fanni Gerber according to burial position, the assumed clothing and deduced jewellery (Figure S.4.7). Finally, an attempt for digital sculpting of the finished reconstruction was made by the newly developed Unreal Engine's Metahuman Creator ("Metahuman Creator," 2021) (Figure S.4.8), however, despite the life-like results, some important facial characteristics, such as maxillary prognathy, could not be implemented to this software, making this reconstruction not entirely plausible. On the other hand, by creating this digital version we showed the future opportunities for such reconstructions. Finally, a grave reconstruction (Main text Figure 1.d) is also created by Fanni Gerber, based on burial position and available data on clothing (Figure S.4.9).

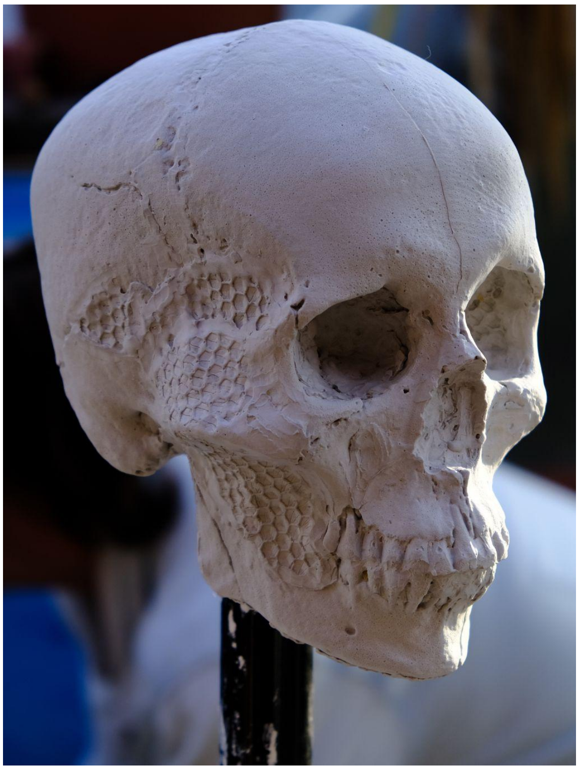

*Fig. S.4.1*

*Plaster cast of the skull by Ágnes Kustár. Photo by Dániel Gerber.*

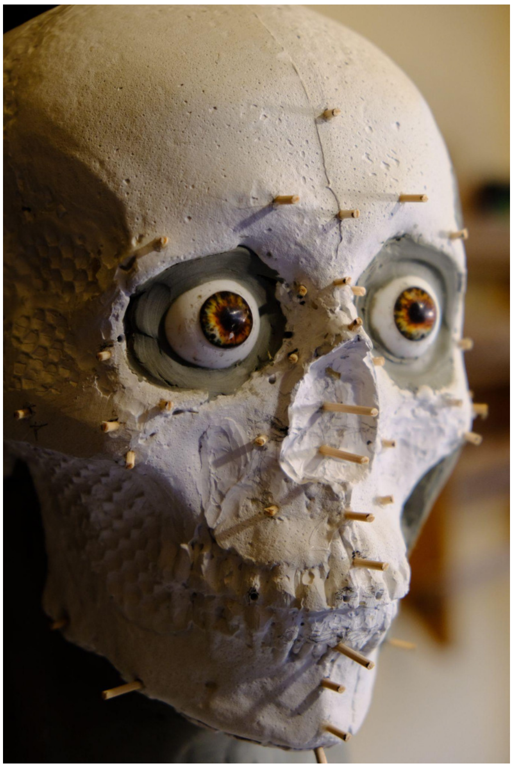

*Fig. S.4.2*

*Pins placed to measuring points and trimmed according to calculated tissue thickness (SI Table 2) by Ágnes Kustár. Photo by Dániel Gerber.*

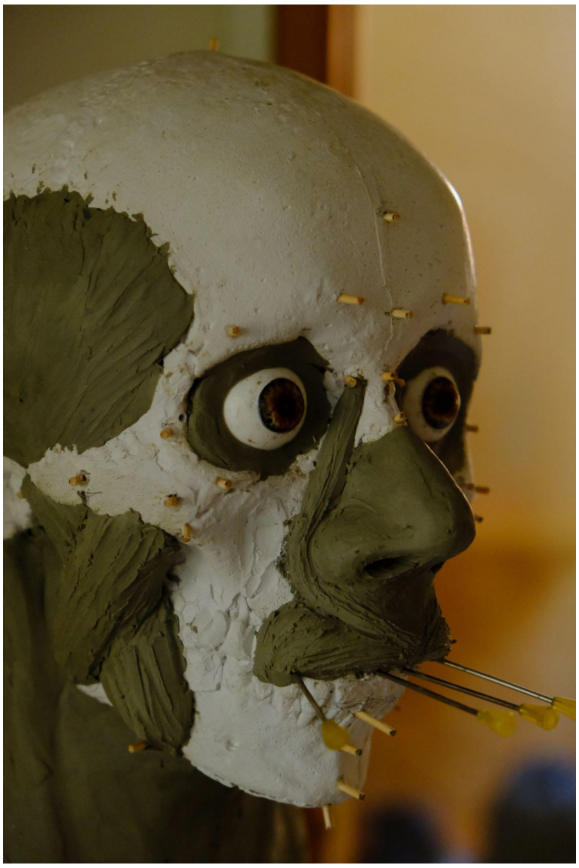

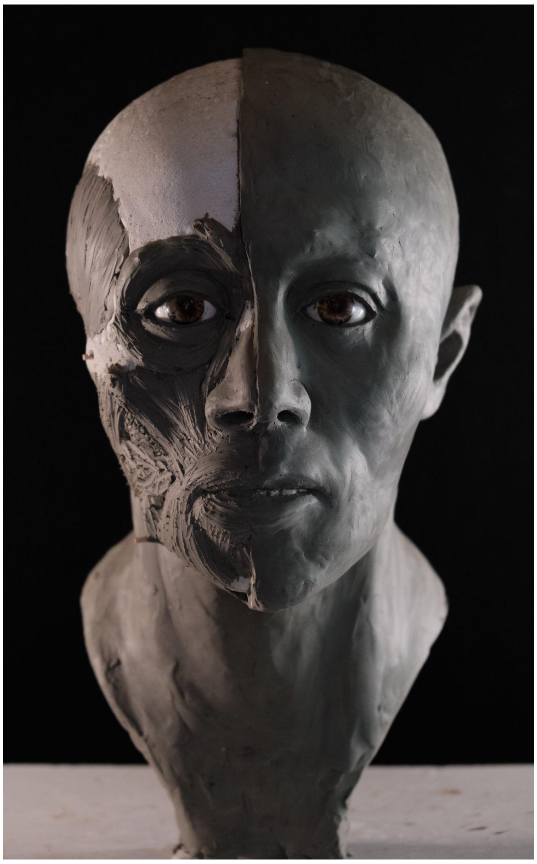

*Fig. S.4.3*

*During the tissue rebuilding process long needles were used to keep monitoring the important measurement points by Ágnes Kustár. Photo by Dániel Gerber.*

*Fig. S.4.4*

*Half of the skull is finished, the other half was kept halfway ready to monitor accuracy by Ágnes Kustár. Photo by Dániel Gerber.*

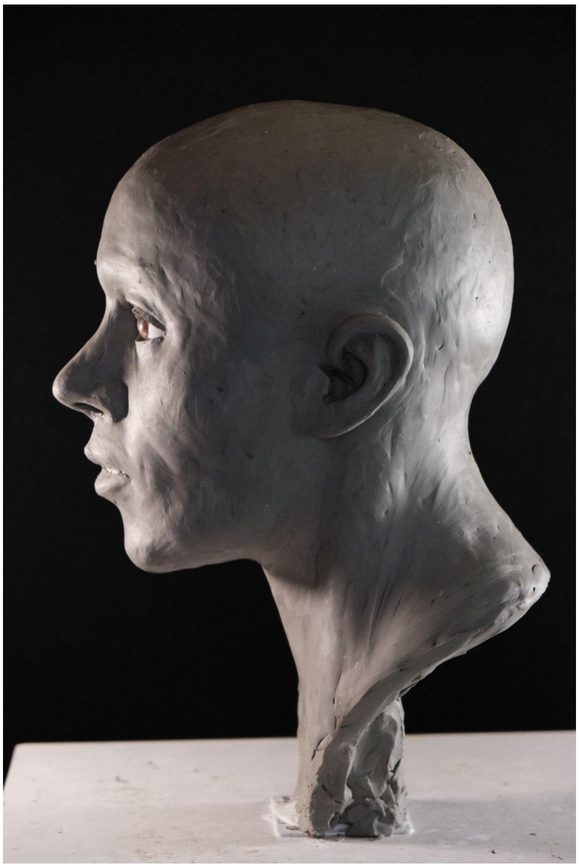

*Fig. S.4.5*

*Profile view of the final clay sculpture by Ágnes Kustár. Photo by Dániel Gerber.*

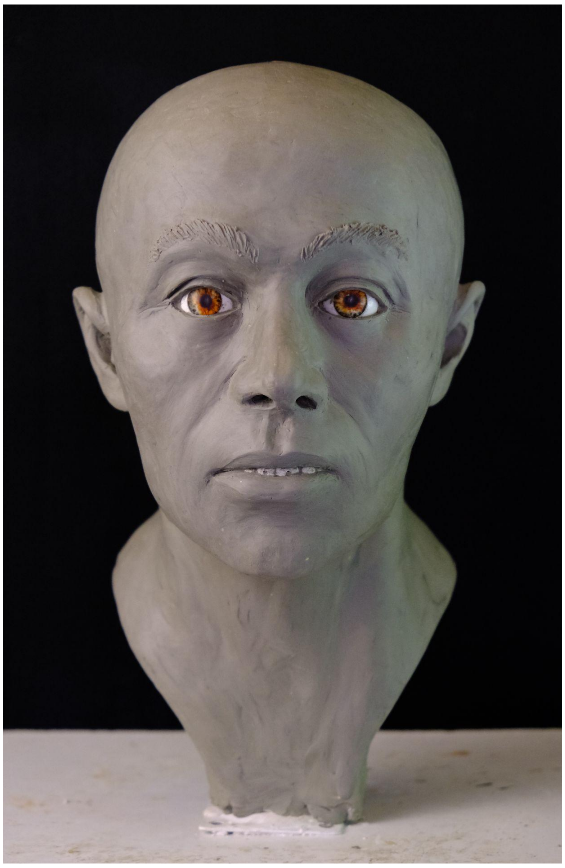

*Fig. S.4.6*

*Frontal view of the final clay sculpture by Ágnes Kustár. Photo by Dániel Gerber.*

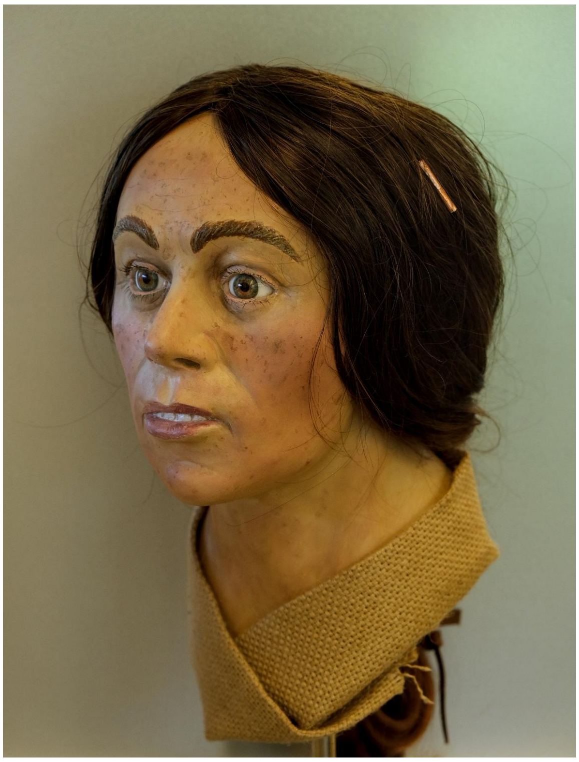

*Fig. S.4.7*

*Finished and coloured reconstruction of individual S13. A bronze bead was pinned to her hair as similar jewellery was found next to her skull. Photo by Dániel Gerber.*

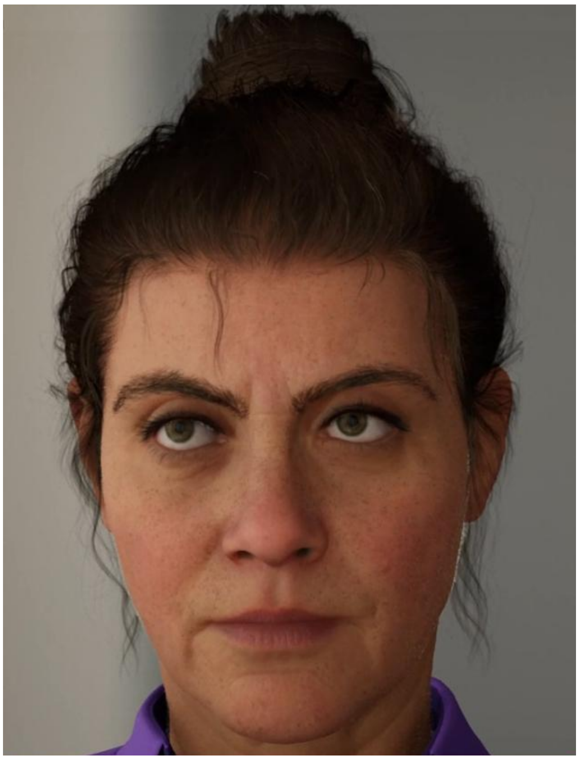

*Fig. S.4.8*

*Digital reconstruction made by Dániel Gerber with Unreal Engine's Metahuman Creator. This image is not accurate, due to the lack of distinguishing features, such as maxillary prognathy, and raw shaping of facial features.*

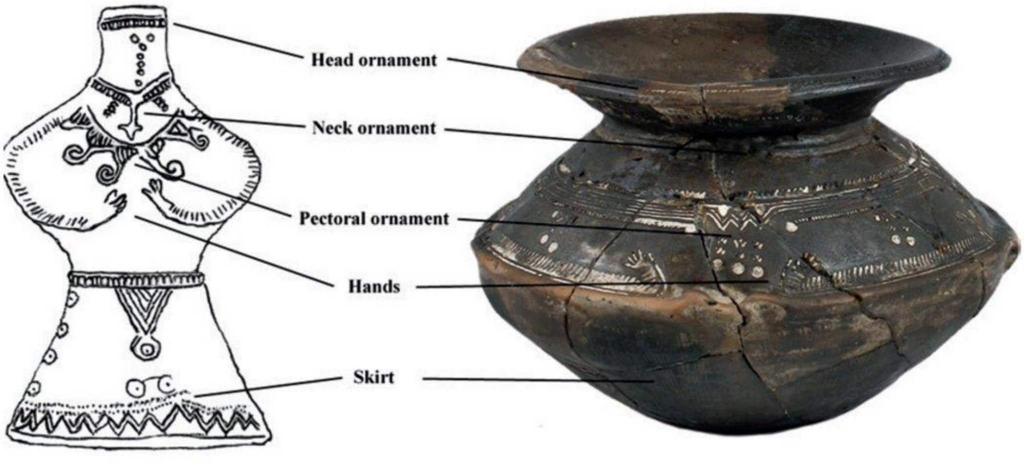


*Fig. S.4.9*

*Reconstruction of clothing style and jewellery based on Encrusted Pottery ornaments (Hajdu et al., 2016).*

### 5) Whole genome analyses

by Dániel Gerber, Balázs Gyuris, Anna Szécsényi-Nagy

#### 5.1) PCA

We used the 'smartpca' software (Patterson et al., 2006) and calculated principal components (PCs) on a set of present-day European, Near Eastern and Caucasian populations (here using their database (Reich, 2021) names: Abkhasian, Adygei, Albanian, Armenian, Assyrian, Balkar, Basque, BedouinA, BedouinB, Belarusian, Bulgarian, Canary_Islander, Chechen, Chuvash, Croatian, Cypriot, Czech, Druze, English, Estonian, Finnish, French, Georgian, German, Greek, Hungarian, Icelandic, Iranian, Irish, Irish_Ulster, Italian_North, Italian_South, Jew_Ashkenazi, Jew_Iranian, Jew_Iraqi, Jew_Libyan, Jew_Moroccan, Jew_Tunisian, Jew_Turkish, Jew_Yemenite, Jordanian, Kumyk, Lebanese, Lebanese_Christian, Lebanese_Muslim, Lezgin, Lithuanian, Maltese, Mordovian, North_Ossetian, Norwegian, Orcadian, Palestinian, Polish, Romanian, Russian, Sardinian, Saudi, Scottish, Shetlandic, Sicilian, Sorb, Spanish, Spanish_North, Syrian, Turkish, Ukrainian), using 590k SNPs in the calculation with an option of shrinkmode=YES, lsq=YES, and the PCAplot.r script from the 'PAPline' for plotting.

#### 5.2.1) Admixture

We made unsupervised and supervised admixture analysis for the whole genome (Figures S.5.2.1 and S.5.2.2) by using the 'ADMIXTURE' software (Alexander et al., 2009) with a fixed set of individuals as sources listed in the Supplementary Table 8, and results seen in Supplementary Table 9a (unsupervised) and 9b (supervised). For supervised admixture we selected sources based on the results of unsupervised analysis, which thus included HG (Antonio et al., 2019; Brace et al., 2019; Fu et al., 2016; Günther et al., 2018; Lazaridis et al., 2014; Mathieson et al., 2018, 2015; Mittnik et al., 2018; Rivollat et al., 2020), early farmer (Turkey_N, Germany_EN_LBK, Hungary_MN_LBK, Serbia_EN_Starcevo)(Feldman et al., 2019; Lipson et al., 2017; Mathieson et al., 2018, 2015; Rivollat et al., 2020) and steppe (Russia_Afanasievo, Russia_Kalmykia_EBA_Yamnaya.SG, Russia_Samara_EBA_Yamnaya, Ukraine_Ozera_EBA_Yamnaya)(Mathieson et al., 2018, 2015; Narasimhan et al., 2019) ancestry populations. We further divided HG sources to Western (WHG: France_Ranchot88, Germany_Mesolithic, Italy_Mesolithic.SG, Italy_North_Villabruna_HG, Luxembourg_Loschbour.DG, Switzerland_Bichon.SG, Wales_Mesolithic) and Eastern (EHG: Russia_EHG, Russia_HG_Karelia, Russia_HG_Samara, Ukraine_Mesolithic, Ukraine_N, Norway_Mesolithic.SG) components for their high genetic dissimilarity based on (Rivollat et al., 2020), but we excluded the GoyetQ2 type component because of their low coverage representative genomes in the database. For results of unsupervised analysis, see Supplementary Figure S.5.2.1, for supervised, see Fig. 2 in main text.

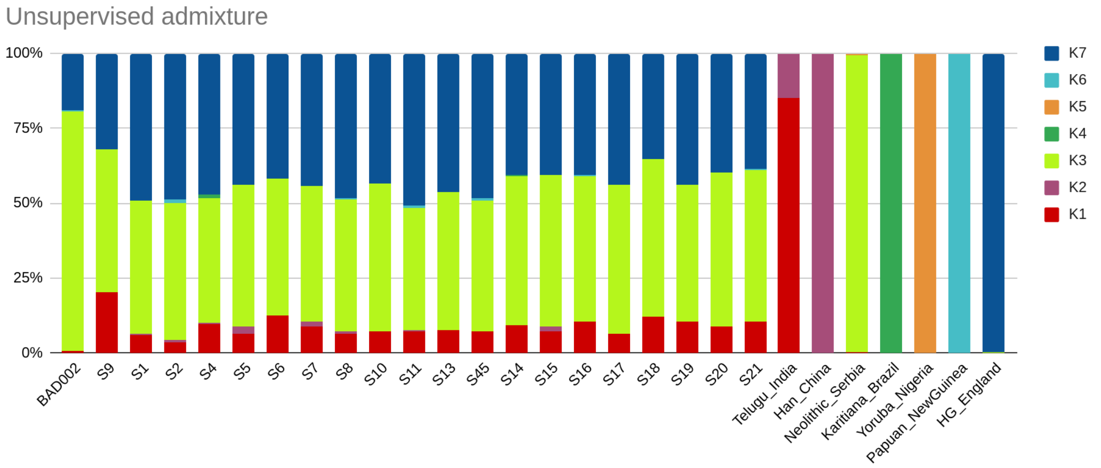


*Fig. S.5.2.1*

*Ancestry proportions of BAD002 and samples from the Balatonkeresztúr site in unsupervised admixture using seven source models. Results show exclusively European associated ancestry in all of the studied individuals.*

#### 5.2.2) Admixture based outlier detection

Outliers were detected using changes in difference quotient based on WHG + EHG component percentages. First, regrouping of the published samples was made according to the first column of Table 9b, then the combined HG component values were sorted within groups, where component differences between the lowest and highest values are more than an arbitrary number of 0.07. Then, difference quotients were set to see where the samples split into two groups, then we marked the ones with the higher HG component. This method resulted in a robust detection of (possible) HG component driven outliers, but in some cases revealed strong group separation, *e.g.* in the case of Serbia_Mokrin_EBA_Maros into low and high HG subgroups. In some cases low HG component outliers were detected, we treated these differently as well. Supplementary Table 8 shows the new groupings for additional population genetic analyses.

#### 5.3) *f4*-statistics

We used *R* packages 'admixr'(Petr et al., 2019) and the 'Admixtools' v7.0(Patterson et al., 2012) to calculate *f4*-statistics in the form of *f4*(W=test HG, X=Serbia Iron Gates, Y=target, Z=Mbuti.DG). In theory, if we consider Serbia Iron Gates as primary sources for HG ancestries in Central European Early Neolithic farmers, this test could reveal subsequent HG admixture regarding geographical origin of such events. Results of this test perfectly showed an additional WHG component in Western European Neolithic populations (*e.g.* in Scotland_N_mediumLowEEF, see Fig. S.5.3.1), and an additional EHG component in Eastern European populations (*e.g.* in Latvia_LN_CordedWare, see Figure S.5.3.2). We applied an average 50k SNP threshold, explained further in SI section 5.4. Despite low coverage, in the case of BAD002 the additional WHG component is apparent, and is significantly positive for Loschbour and Italian HG-s (Figure S.5.3.3). For Bk-II (Figure S.5.3.4) and Bk-III (Figure S.5.3.5) the results indicate extra WHG and EHG characteristic admixture (Figure S.5.3.4), however this seems less strong compared to the previous WHG or EHG driven admixtures. This could indicate several possibilities: (a) the extra HG component is more similar to Serbian Iron Gates HG, and the additional heterogeneous affinities only reflect subsequent mixtures with Bronze

Age populations of various genetic and geographical backgrounds, (b) the extra HG component is of multiple origin, thus it behaves like Iron Gates HG, but also shows extra affinities to particular EHG and WHG groups, (c) the extra HG is yet to be described, and while it is more similar to Serbian Iron Gates, it shows “random” affinities to various EHG and WHG groups, and (d) options “b” and “c” are both possible. To test these scenarios we ran different analyses which could disclose any of the listed possibilities.

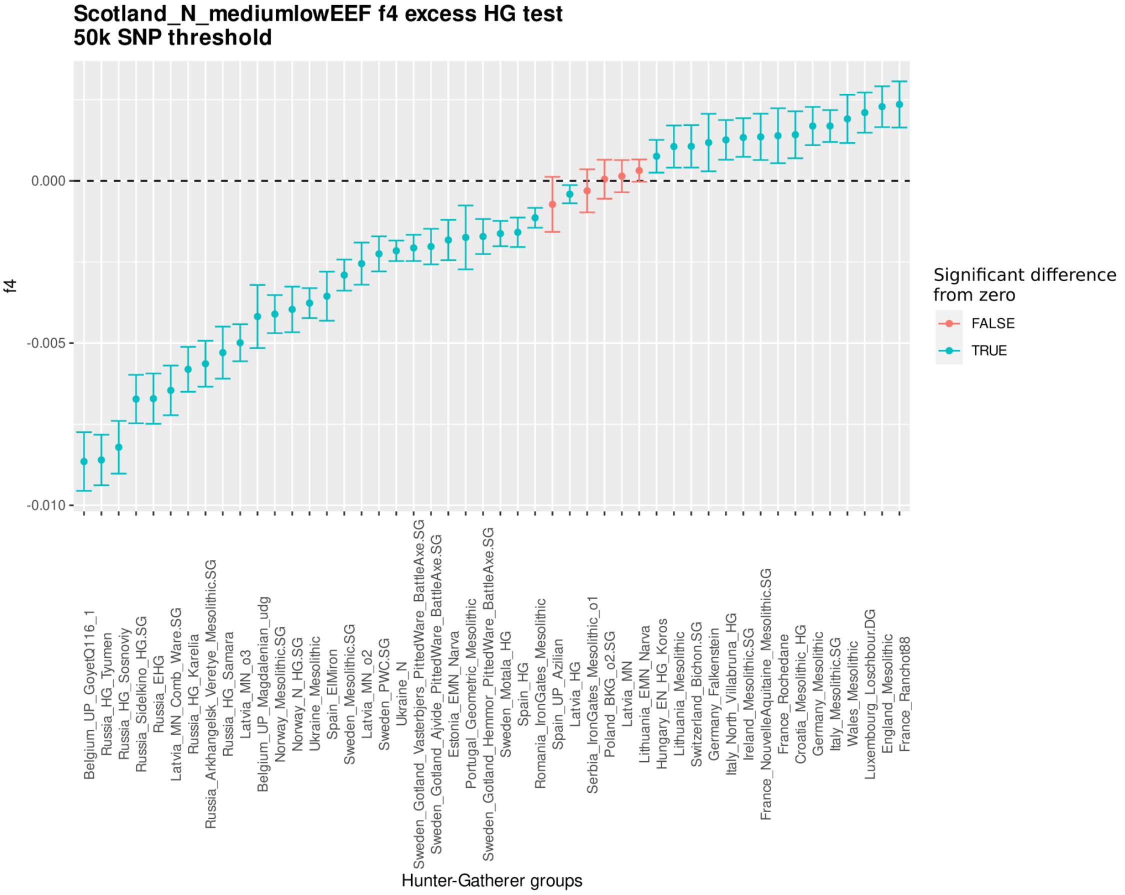


*Fig. S.5.3.1*

*The result of the f4-statistics of the Scotland_N_mediumlowEEF as target – in the form of (W=test HG, X=Serbia Iron Gates HG, Y=target, Z=Yoruba), where the HG component shows the most resemblance to Western HG-s.*

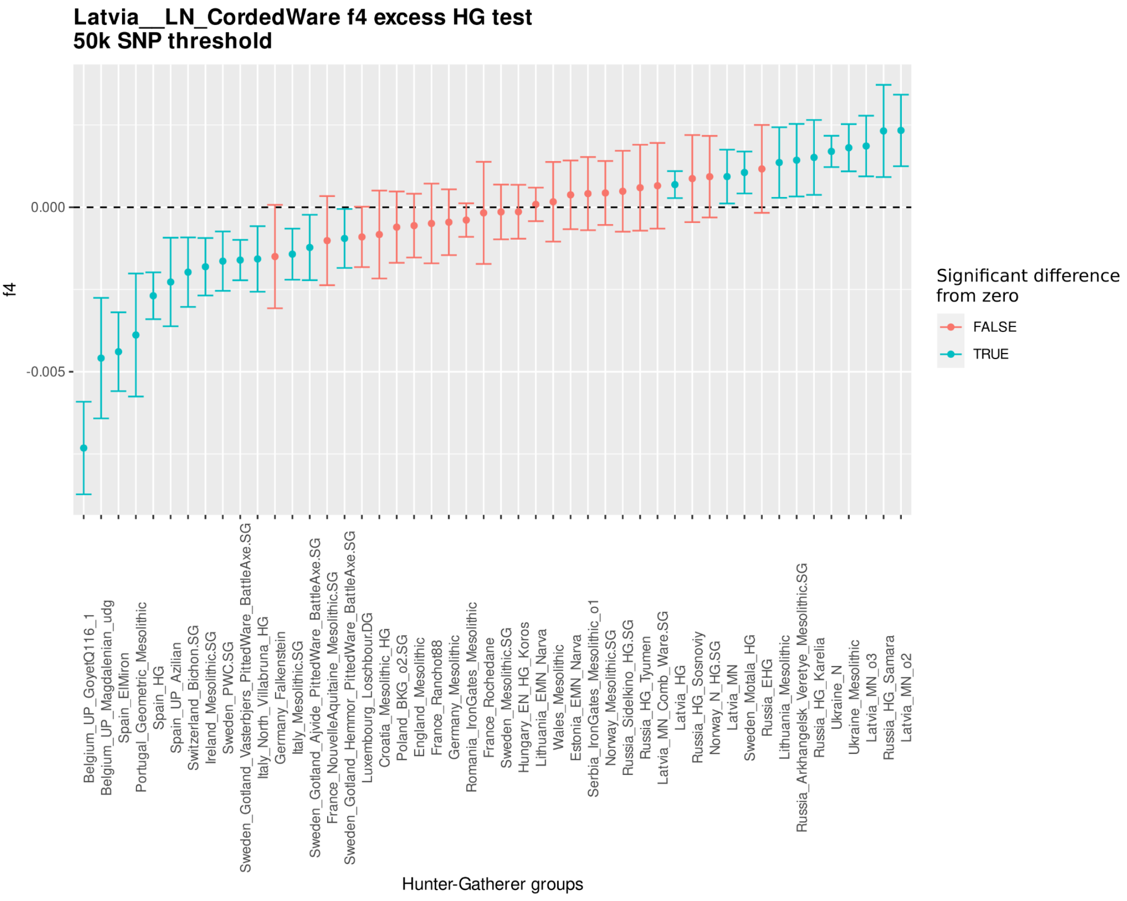


*Fig. S.5.3.2*

*The results of the f4-statistics of Latvia_LN_CordedWare as target – in the form of (W=test HG, X=Serbia Iron Gates HG, Y=target, Z=Yoruba). Results show EHG excess for this population.*

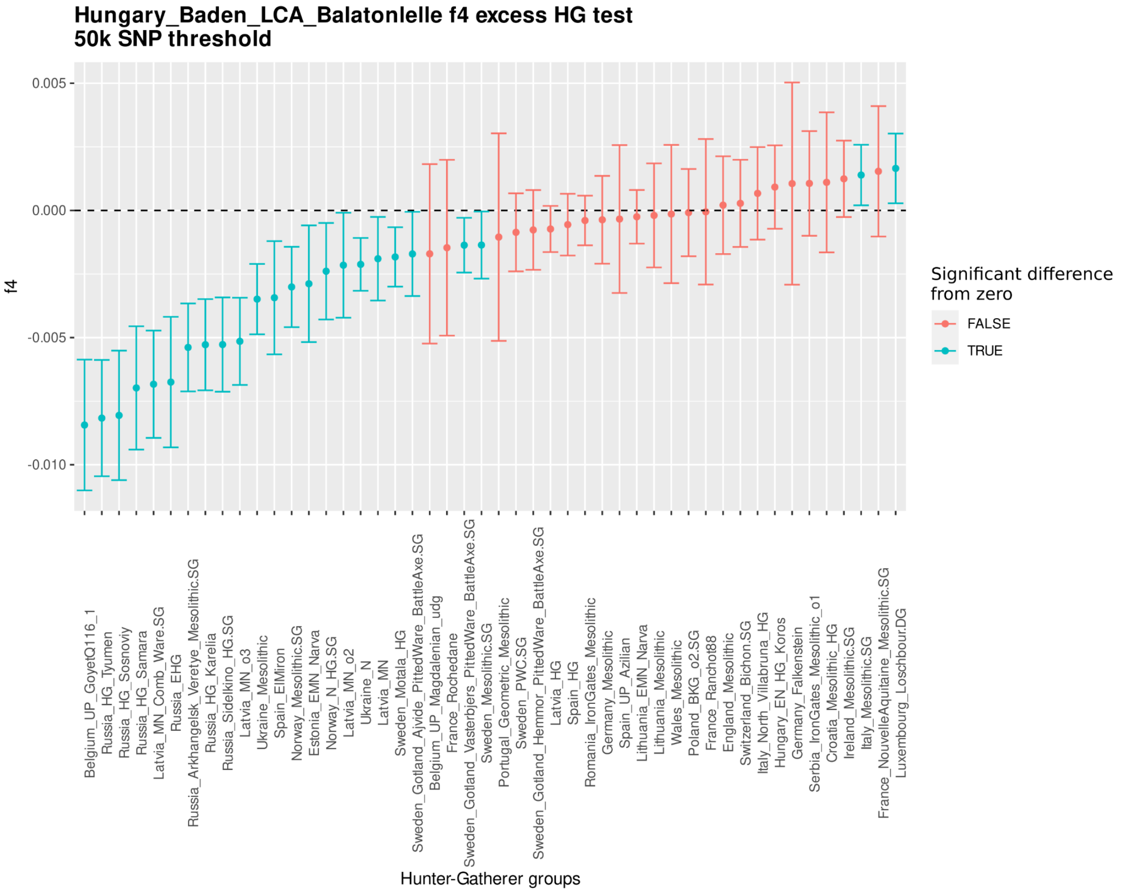


*Fig. S.5.3.3*

*The results of the f4-statistics of BAD002 individual as target – in the form of (W=test HG, X=Serbia Iron Gates HG, Y=target, Z=Yoruba). Despite low coverage and high standard error, the WHG excess is apparent for this individual (Malmström et al., 2019; Mittnik et al., 2018; Sánchez-Quinto et al., 2019).*

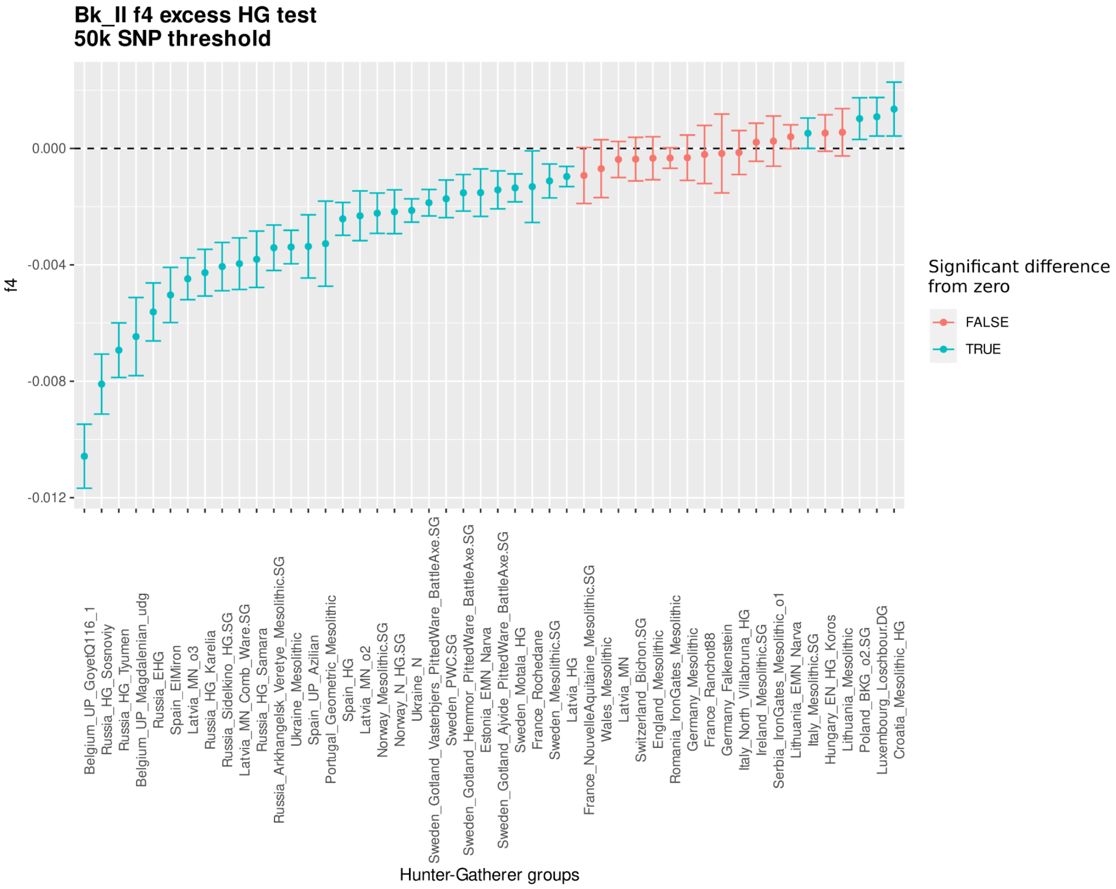


*Fig. S.5.3.4*

*The results of the f4-statistics of the Bk-II individuals as targets – in the form of (W=test HG, X=Serbia Iron Gates HG, Y=target, Z=Yoruba). While affinities to WHG populations seem to be stronger, excess of Southern Baltic HG ancestry also appears with insignificant f4 value in Bk-II.*

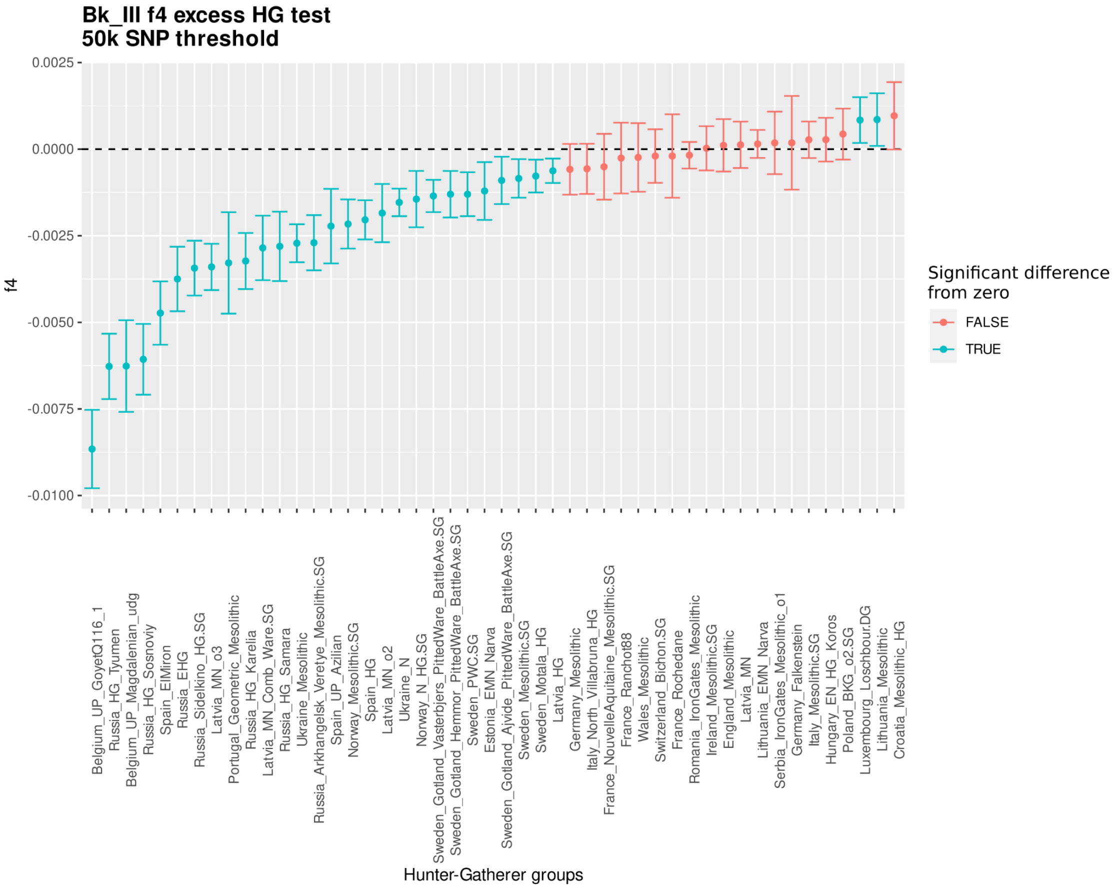


*Fig. S.5.3.5*

*The results of the f4-statistics of the Bk-III individuals as targets – in the form of (W=test HG, X=Serbia Iron Gates HG, Y=target, Z=Yoruba). Despite lower levels of HG ancestry a very similar pattern to Bk-II can be observed.*

#### 5.4) DATES analyses

We ran DATES (Chintalapati et al., 2022) analyses in order to see when HG and steppe ancestries mixed with Bk-I and Bk-II groups. Also, we calculated with a generation time of 25 years. Additionally, we chose 2425 BCE as the date of origin for Bk-I, and 2000 BCE for Bk-II according to radiocarbon estimates, see SI section 1.8. Sources for this analysis was based on ADMIXTURE (see SI section 5.2.1), qpAdm results (Bk-I: EEF ~53±3.7%, EHG ~23±2.8%, WHG ~6±3.7%, Caucasus HG ~18±3.2%, p=0.557; Bk-II: EEF ~40±2%, EHG ~39±3%, WHG ~13±2.7%, Caucasus HG ~8±2%, p=0.0917). For Bk-I we used Germany_EN_LBK (early Neolithic component) and Russia_Samara_EBA_Yamnaya (steppe component) as sources, which resulted in 21.201±11.398 generations prior admixture (between ~2675 and ~3225 BCE, mean ~2950 BCE), that is highly consistent with the timing of steppe ancestry arrival to Central-Eastern Europe. According to qpAdm and ADMIXTURE there are both EHG and WHG driven ancestries for Bk-II, thus we used Luxembourg_Loschbour.DG as WHG, Ukraine_N as EHG, Serbia_IronGates_Mesolithic as mixed HG characteristic component, Turkey_N/Germany_EN_LBK and Russia_Samara_EBA_Yamnaya as sources in pairs, respectively. Turkey_N and Serbia_IronGates_Mesolithic provided 34.754±15.092 generations prior admixture (between ~2500 and ~3250 BCE, mean ~2875 BCE), which indicates a relatively recent, but not an unpaired estimate for such event (Furtwängler et al., 2020). However, when we estimated WHG (consistent with estimates of (Freilich et al., 2021)) and EHG separately (see SI Table 5.4), the results showed high asymmetry, and a much more recent introgression of EHG ancestry to Bk-II. The admixture date of WHG and EHG components overlaps with the separate EHG admixture. The slightly deeper time estimate for the latter is either the result of the distortive effect of Serbia_IronGates_Mesolithic ancestry found in almost all Neolithic populations, or the EHG dominant component is previously admixed with some WHG-s (Chintalapati et al., 2022). Based on these results we conclude that three major HG admixture events occurred among the ancestors of Bk-II: the first was the Serbia_IronGates_Mesolithic associated admixture around the beginning of the spread of agriculture to Europe, commonly observable among most Neolithic populations. Secondly, there was a WHG characteristic admixture around the beginning of the third millennium BCE. A similar event was observed in Germany at the Blatterhohle site (Lipson et al., 2017) and in the UK (Scotland, Raschoille cave (Patterson et al., 2022). The third was an EHG characteristic admixture, parallel to the only known case from the Ukraine (Dereivka I (Mathieson et al., 2018)), although DATES provided controversial results for this individual (10.227±17.019, between ~2780 BCE and 3705 BCE, mean ~3000 BCE). The Germany_EN_LBK and Russia_Samara_EBA_Yamnaya admixture date for Bk-II also suggests relatively recent admixture between a population of farmers and of high steppe ancestry.

*SI Table 5.4*

*Tables of DATES results with calculated generations prior admixture. Upper cells show generation time with SE, while the lower cells show the corresponding estimated absolute chronological date with SE considering 25 years of generation time.*

| **Bk-I** | **IronGatesHG** | **WHG** | **EHG** | **EEF** | **Steppe** |
|---|---|---|---|---|---|
| **IronGatesHG** | | | | | |
| **WHG** | | | | | |
| **EHG** | | | | | |
| **EEF** | | | | | 21.201±11.39 |
| **Steppe** | | | | 2950±275 BCE | |

| **Bk-II** | **IronGatesHG** | **WHG** | **EHG** | **EEF** | **Steppe** |
|---|---|---|---|---|---|
| **IronGatesHG** | | | | 34.754±15.09 | |
| **WHG** | | | 21.145±12.95 | 30.261±17.03 | |
| **EHG** | | 2525±325 BCE | | 17.049±9.48 | |
| **EEF** | 2875±375 BCE | 2750±425 BCE | 2425±225 BCE | | 16.581±4.33 |
| **Steppe** | | | | 2400±100 BCE | |

#### 5.5) *f3*-statistics

The outgroup *f3*-statistics and clustering analyses were based on (Patterson et al., 2012) and made by Admixtools v7.0 and 'R' packages 'admixr', 'admixtools'. We checked HG affinities by f3-statistics. Since f4, preliminary qpAdm and DATES results suggested that the increased HG component in Bk-II is (1) of at least double origin, (2) has both EHG and WHG characteristics, (3) is independent from mixed characteristic HG groups, such as Iron Gates HG, and (4) is dissimilar to “pure” EHG groups from Scandinavia, Estonia, Ukraine and Russia and shows some affinity to certain Latvian, Lithuanian, Polish, Iron Gates and Croatian HG-s (all having EHG ancestry). All of this suggests that the source of this particular genetic makeup is yet to be described and probably originated somewhere from unsampled regions of the Balkans or Eastern Europe. We wanted to trace back this specific ancestry to see whether it appeared before or after Bk-II among Neolithic and Bronze Age groups, thus we designed a simple experiment to unravel similar HG ancestry patterns among database data: we ran pairwise *f3* between target and HG populations, using Mbuti outgroup (Supplementary Table 11). In theory, *f3* distances between the target and a certain HG would only depend on the target’s HG component, as other components would roughly be equally distant to all HG groups. Accordingly, all targets can be clustered by their distance pattern to various HG groups. To avoid noise, we excluded HG groups that had less than a median of 100k overlapping SNPs with all the targets. The final run covers HG groups all over Europe, from Iberia to the Ural mountains (see Table 8), thus the geographical and genetic resolution is deep enough for a delicate analysis (for results, see Supplementary Table 11). We calculated euclidean distances between targets based on these *f3* results, and used “complete” clustering method in ‘R’ by ‘hclust’ (Figure S.5.5.1). This resulted in grouping together Bk-II and Bk-III with a number of outliers and groups, but none of them had an equal or higher amount of HG component than Bk-II, excluding them as possible actual sources.

We also made a 10,000 bootstrap resampling, where we bootstrapped HG components from the distance matrix, and then clustered targets in each replica similarly as described above. This resulted in additional groups to consider, compared to the original clustering, but most of them had high replication numbers (Figure S.5.5.2, SI Table 5.5.1). Notably, a few western Neolithic groups also appear between 3000 and 4000 bootstrap replicas, from which the occurrence of other groups and outliers drops significantly. Most of these low occurring groups are quite similar to mid occurring (between 6000-8000 replicas, also they are a sister branch in the original cluster to the branch of Bk-II) newly appearing groups/individuals by geographical distribution, age and genomic makeup. These results are consistent with previous observations from DATES and uniparental markers: most probable WHG admixture and origin of paternal lineages likely trace back to those regions represented by these groups (Figure S.5.5.2), suggesting potential sources for Bk-II.

Also, a number of contemporary Bronze Age groups appear in close proximity to Bk-II besides Bk-III. They are mostly outliers from proximate regions, most likely as a result of the regional expansion of the Kisapostag culture, which we discuss further in the main text. On the other hand, not only individuals but groups as well from much farther regions and/or chronological layers appear to carry a Bk-II like HG component, pointing to possible origin of this ancestry. Interestingly, many of the connections appeared here are supported by indirect or even direct uniparental connections as well (e.g. Tollense, Mokrin group, Poland_EBA, etc.).

Finally, we also made outgroup *f3* with our groups against all of the reference populations, for results, see Supplementary Table 10. The aim of this experiment was to guide us during qpAdm analysis, where we applied the same SNP threshold filter, further described in SI section 5.6.

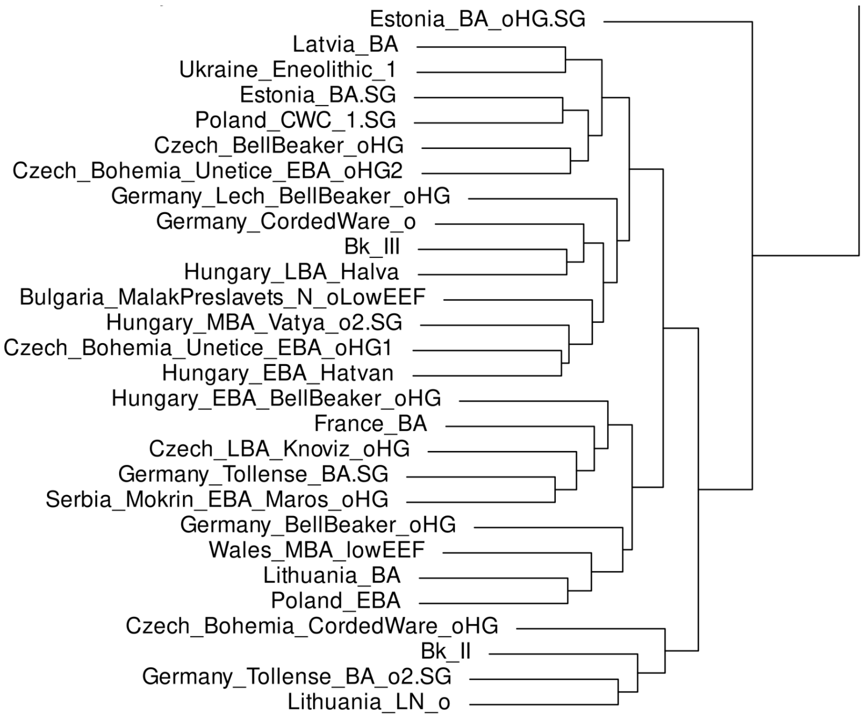


*Fig. S.5.5.1*

*Outgroup f3 with euclidean distance based clustering, only the subcluster of Bk-II is shown. Most of these groups/samples postdate Bk-II and are restricted to the Carpathian Basin and vicinity regions, except for groups from the Baltics and the British Isles.*

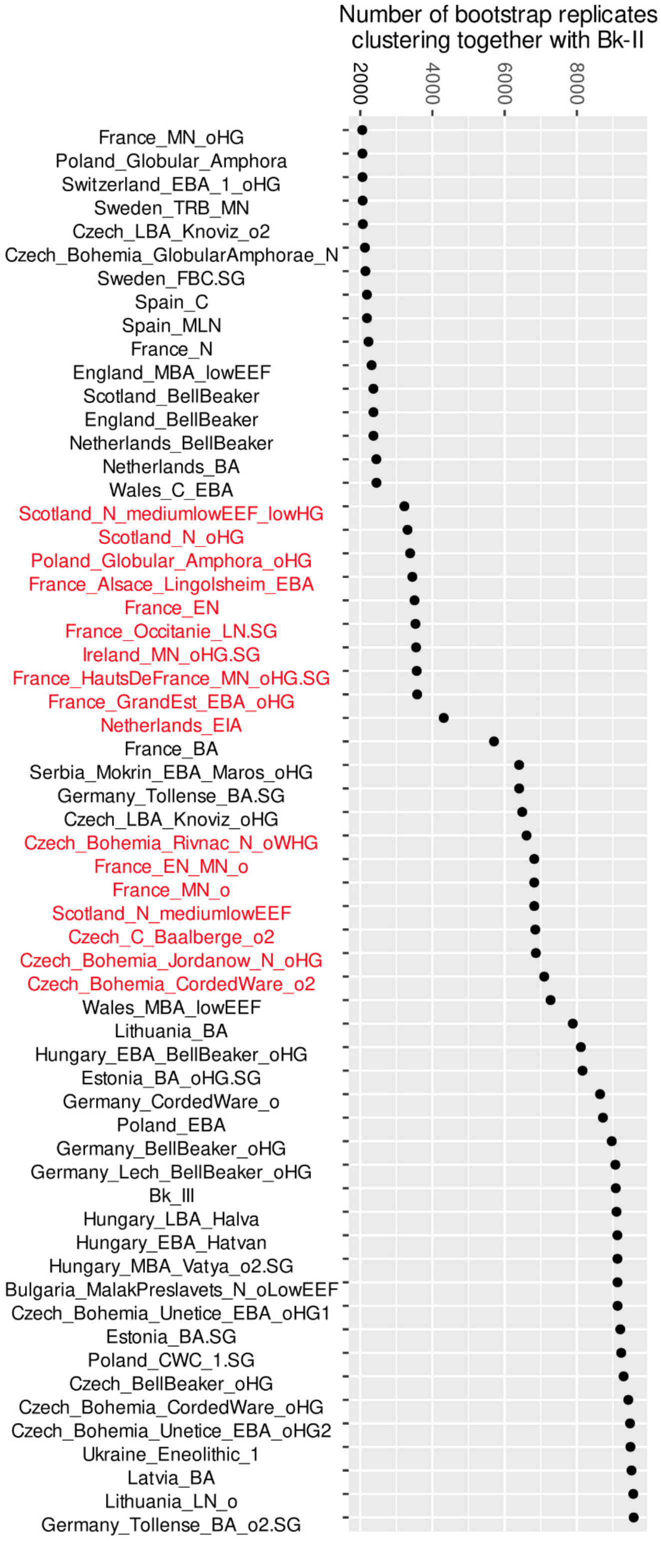


*Fig. S.5.5.2*

*Bootstrapped f3 clustering provided high statistical support for the original clustering, and also showed increased co-clustering of a neighbouring sub-cluster, containing mostly Western Neolithic individuals and groups of high HG components.*

*SI Table 5.5.1*

*Sample/group data of co-clustering individuals with 5000+ positive bootstrap replicas.*

| **Populations/individuals** | **Bootstrap** | **Date (radiocarbon or archaeological)** | **Average HG ancestry (%)** |
|---|---|---|---|
| Germany_Tollense_BA_o2.SG (WEZ56.SG) | 9574 | 1300-1200 BCE | 24.91 |
| Lithuania_LN_o (Spiginas2) | 9563 | 2130-1750 calBCE | 38.19 |
| Latvia_BA | 9514 | 810-230 calBCE | 29.62 |
| Ukraine_Eneolithic_1 (I4110) | 9484 | 3630-3380 calBCE | 36.2 |
| Czech_Bohemia_Unetice_EBA_oHG2 (VLI051) | 9473 | 1880-1690 calBCE | 31.28 |
| Czech_Bohemia_CordedWare_oHG (TRM001) | 9425 | 2900-2500 BCE | 25.19 |
| Czech_BellBeaker_oHG (I7286) | 9295 | 2400-2200 calBCE | 26.43 |
| Poland_CWC_1.SG | 9228 | 2600-2250 BCE | 34.77 |
| Estonia_BA.SG | 9203 | 1270-400 calBCE | 31.48 |
| Czech_Bohemia_Unetice_EBA_oHG1 (VLI050) | 9127 | 1950-1750 calBCE | 30.23 |
| Bulgaria_MalakPreslavets_N_oLowEEF (I1108) | 9125 | 5800-5400 BCE | 37.71 |
| Hungary_MBA_Vatya_o2.SG (RISE479) | 9124 | 2000-1500 BCE | 35.68 |
| Hungary_EBA_Hatvan (I1502) | 9122 | 2200-1970 calBCE | 32.67 |
| Hungary_LBA_Halva (I20749) | 9099 | 1700-800 BCE | 25.03 |
| Bk_III | 9078 | 1870-1620 calBCE | 30.04 |
| Germany_Lech_BellBeaker_oHG (HUGO_180sk1_d) | 9067 | 2460-2210 calBCE | 27.52 |
| Germany_BellBeaker_oHG (I5659) | 8967 | 2500-2000 BCE | 24.42 |
| Poland_EBA (I6579) | 8722 | 2340-2040 calBCE | 27.4 |
| Germany_CordedWare_o (I1540) | 8642 | 2500-2050 BCE | 25.13 |
| Estonia_BA_oHG.SG (S19_X12_1.SG) | 8157 | 900-790 calBCE | 39.46 |
| Hungary_EBA_BellBeaker_oHG (I3528) | 8112 | 2560-2300 calBCE | 34.68 |
| Lithuania_BA | 7888 | 1257-797 calBCE | 23.94 |
| Wales_MBA_lowEEF (I5440) | 7272 | 1500-1322 calBCE | 17.75 |
| Czech_Bohemia_CordedWare_o2 | 7095 | 2848-2495 calBCE | 29.1 |

| | | | |
|---|---|---|---|
| (VLI009) | | | |
| Czech_Bohemia_Jordanow_N_oHG (NER001) | 6865 | 4235-3995 calBCE | 36.89 |
| Czech_C_Baalberge_o2 (I14168) | 6849 | 4300-3500 BCE | 27.63 |
| France_EN_MN_o (LBR005) | 6820 | 5216-4953 calBCE | 33.76 |
| France_MN_o | 6820 | 4777-4548 calBCE | 37.92 |
| Scotland_N_mediumlowEEF | 6820 | 3942-3037 calBCE | 30.03 |
| Czech_Bohemia_Rivnac_N_oWHG (TUC003) | 6606 | 3090-2890 calBCE | 32.82 |
| Czech_LBA_Knoviz_oHG (I15959) | 6486 | 1300-800 BCE | 28.26 |
| Germany_Tollense_BA.SG | 6402 | 1300-1200 BCE | 28.4 |
| Serbia_Mokrin_EBA_Maros_oHG | 6400 | 2100-1800 BCE | 28.25 |
| France_BA (I16791) | 5705 | 1300-900 BCE | 17.46 |

#### 5.6) qpAdm

qpAdm calculations were performed for each of the studied groups for which we used 'Admixtools' v7.0 and 'R' package 'admixtools' (Patterson et al., 2012). Potential sources of ancestries were selected based primarily on principal component analysis (PCA), ADMIXTURE, and uniparental analyses results to avoid unnecessary model testing and reduce computational time. For outgroups we chose 'Mbuti.DG', 'Russia_Ust_Ishim.DG', 'Russia_MA1_HG.SG',, 'Russia_Kostenki14', 'Iran_GanjDareh_N', 'Luxembourg_Loschbour.DG', 'Russia_HG_Tyumen, 'Israel_Natufian', 'Turkey_Boncuklu_N', 'Russia_Afanasievo', 'Russia_Steppe_Maikop', 'Belgium_UP_GoyetQ116_1'. For BAD002 we added 'Serbia_IronGates_Mesolithic' and 'Germany_BellBeaker' and excluded 'Luxembourg_Loschbour.DG' and 'Russia_HG_Tyumen'. We created a custom 'R' script (provided in the 'PAPline' package) for running all possible combinations of single, dual and triple ancestry sources. Since qpAdm results tend to give high *false* positive rates in case of low coverage, we restricted analyses to source populations that have at least 50,000 SNPs overlap with BAD002 and 100,000 SNPs overlap with the Bk groups. The script automatically rejects combinations where negative components appear or standard errors reach a limit (we restricted acceptable scenarios where SE is lower than 0.1 and does not exceed the weight of the corresponding component) while it provides a list of best combination candidates (*i.e.* models that were not rejected in an ascending order of *p*-values).

##### 5.6.1) BAD002

We restricted qpAdm analysis to European Neolithic populations without steppe ancestry, as BAD002 lacks this component. Most models required minor extra WHG (either modelled by France_MontAime_MLN_oHG1.SG or Luxembourg_Loschbour.DG) next to Neolithic populations mostly from Western Europe, however, there were also fitted one way models using groups from Iberia, France, Poland, Bohemia, British Isles and Switzerland. Following Occam's theorem, if we hypothesise a single source for BAD002, outgroup f3 results would unravel best affinities to populations in question. Outgroup f3 shows close affinities to the aforementioned populations, where France_MontAime_MLN_oHG1.SG is the closest to BAD002, followed by

France_MontAime_MLN.SG and Spain_EN. We ran f4 in the form of f4(BAD002, Spain_EN, France_MontAime_MLN.SG, Mbuti) to see whether an excess of allele sharing towards these sources exists for BAD002, however, it resulted in no significant divergence from zero. Since Spain_EN always requires additional WHG in qpAdm models, as well it is much more distinct chronologically from BAD002, we rejected this group as the direct source. On the other hand, France_MontAime_MLN.SG is sufficient as a single source (*p*=0.27) as well as it chronologically overlaps BAD002, and while Ireland and Spain represented most frequently as regions of origin, we ultimately selected this as the most probable source. However, genetic links to the aforementioned regions can not be excluded, especially in the light of archaeological context (Bondár, 2020; Bondár et al., 2021; Bondár and Szécsényi-Nagy, 2020). For results, see Supplementary Table 12.

#### 5.6.2) Balatonkeresztúr site

Since none of the samples/groups of similar HG composition from outgroup f3 clustering possess enough portion of such ancestry to explain Bk-II, we concluded that these are likely other descendants of the actual source or directly of Bk-II. Accordingly, we excluded these from qpAdm analyses. Many probable models were calculated during the qpAdm loop, thus we applied the following method to find the most plausible model: considering the most parsimonious scenario as only two sources explain the target, which combinations were scored according to (1) standard errors, i.e. the lowest standard error gets the best rank, (2) weight balance, *i.e.* the closest the proportions to 50-50%, the better rank it gets, and (3) outgroup f3 distance. We refer to this dual ancestry ranking system as DARS in further descriptions.

### 5.6.2.1) Bk-I

Croatia_EBA_Vucedol_3 (N=1) appears in the closest proximity to individual S9 by far to others (Supplementary Table 11b) in outgroup f3, thus we included this individual to all of our models for the first run of qpAdm rotation. After the initial results, we ran an independent qpAdm with the most probable populations based on occurrence frequency, where we added Croatia_EBA_Vucedol_3 again without setting it as a fixed source. After the second run, Croatia_EBA_Vucedol_3 appeared again in almost all acceptable models, thus we conclude that this individual truly represents one of the sources for S9. On the other hand, Croatia_EBA_Vucedol_3 lacks steppe ancestry, which is prominent in S9, meaning that at least two sources need to explain Bk-I. Most models (492 out of 535) suggest a three-way admixture of Croatia_EBA_Vucedol_3, another Neolithic source of no steppe ancestry and a high steppe ancestry group. However, if we consider the most parsimonious scenario, i.e. the two-way admixture model, then we end up with 42 possible sources for S9 besides Croatia_EBA_Vucedol_3. Most of these are equally probable, thus we applied DARS to see which explains Bk-I the best. According to the combined results Russia_Srubnaya_Alakul_lowHG.SG and Russia_Srubnaya_Alakul.SG are the two best ranked groups, followed by Lithuania_LN and Poland_Southeast_CordedWare.SG, Russia_Andronovo.SG, Russia_Fatyanovo_BA.SG, etc. Since the Srubnaya group is geographically distant and chronologically postdating Bk-I, we can hypothesise that this population is only the best proximate to describe Bk-I, related to 3rd mill. BCE steppe populations. Interestingly, Lithuania_LN also possesses the best weight and standard error ranks and the same specific R1a paternal lineage as S9, meaning that in composition (and also geographically and chronologically) this group is more similar to Bk-I. In summary we conclude that the actual source for Bk-I is yet to be sampled, but can be described as a group with high steppe ancestry, maybe in the vicinity of the Baltics. For results, see Supplementary Table 13.

### 5.6.2.2) Bk-II

We fixed Ukraine_EBA, and Serbia_IronGates_Mesolithic for qpAdm loop, as the former has sufficiently high HG ancestry that can potentially explain Bk-II, otherwise we expect a “pure” HG source represented by the latter. Following the most parsimonious way of having only two sources, we first examined the two-way admixture models (summing up to 19), which all contained Ukraine_EBA. Since Ukraine_EBA contains mainly EHG characteristic HG ancestry and also contains some steppe ancestry, mostly Neolithic groups appeared as the second source with no Steppe and mostly WHG characteristic HG ancestry. By applying DARS, Sweden_FBC and Poland_GAC.SG appeared in first place with equal ranks, in second Sweden_Ansarve_Megalithic.SG and Czech_Bohemia_GlobularAmphorae_N with equal ranks, and in third France_LaClape_LN_EBA_Veraza.SG, which groups have similar characteristics of having higher WHG characteristic ancestry. None of these groups appear above 3000 bootstrap replicas in *f3* clustering, suggesting that the actual source for Bk-II is not represented in the database. However, the geographical, archaeological and genetic proximity of these candidates to good hits in *f3* clustering, the origin of paternal lineages and DATES results all indicate that the actual source of Bk-II does belong to this milieu. On the other hand, Ukraine_EBA remains a bit controversial, as *f3* clustering placed this individual far from the Bk-II subcluster. Despite this candidate having the same R1b haplogroup found in Bk-III, chronologically overlapping with Bk-II and explaining it well by composition, we wanted to analyse the three component models further. Here we fixed Sweden_FBC as the Neolithic component, as this group is closer to Bk-II in outgroup *f3* than Poland_GAC.SG. We also fixed Ukraine_N, as we expect an independent EHG characteristic mixture in the three-way scenario. In this run, no plausible models were calculated, thus we kept the two-way admixture model with Ukraine_EBA with the notion that this individual may only represent the true source for Bk-II only by composition, despite having fairly similar makeup in f4 statistics to Bk-II (Figure S.5.6.2.2). For results, see Supplementary Table 14.

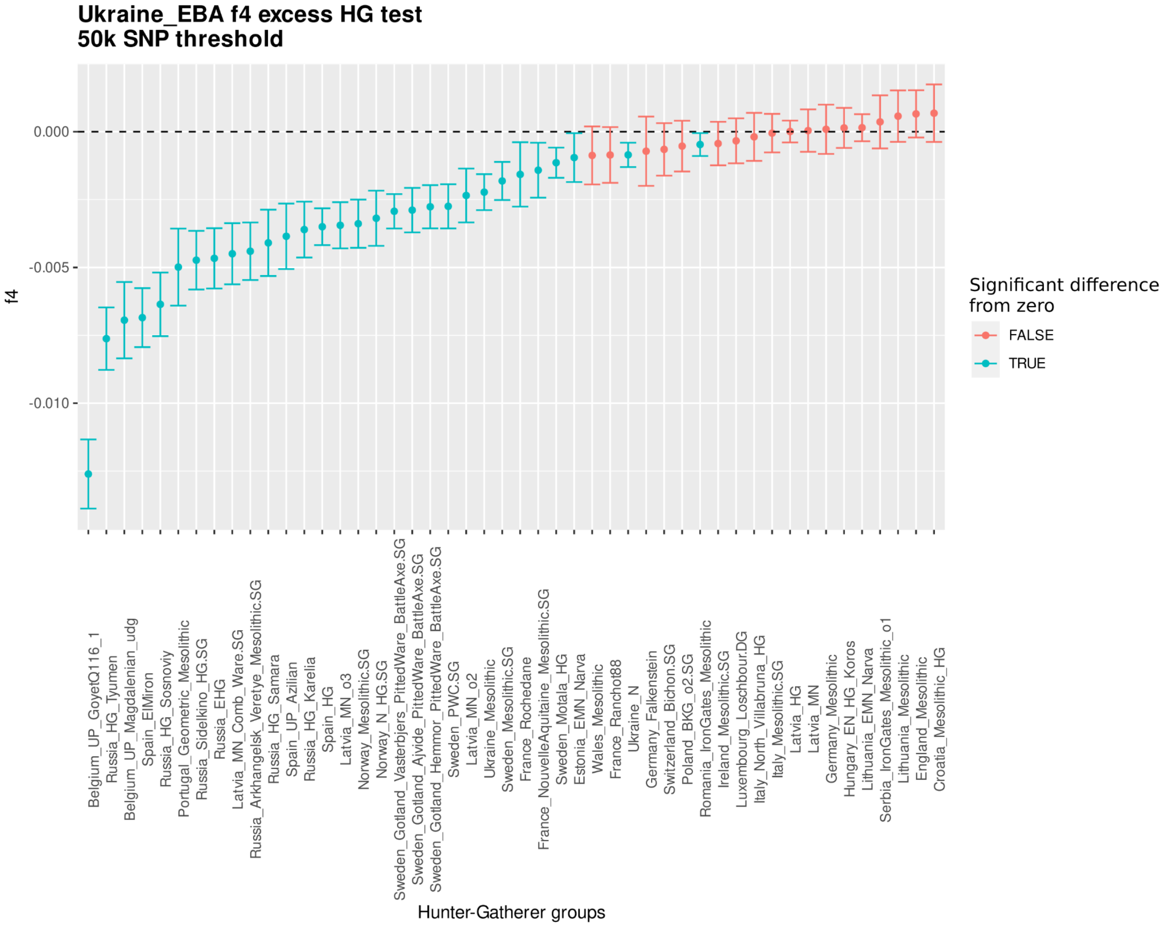

*Fig. S.5.6.2.2*

*F4-statistics of Ukraine_EBA in the form of (W=test HG, X=Serbia Iron Gates HG, Y=target, Z=Yoruba). Similar makeup to Bk-II can be observed, but the overall HG composition loosely resembles Bk-II's, not only by the lack of WHG ancestry, but also by weaker connection to Baltic HG groups.*

##### 5.6.2.3) Bk-III

For the qpAdm analysis for Bk-III we considered Bk-II as a fixed source for: (1) chronological, geographical and archaeological continuity, and (2) possessing highly similar genetic features to Bk-II and having the closest distance from Bk-III in outgroup *f3*. In this run 8283 possible models were fitted, for which we used the most parsimonious scenario: Bk-II admixture with a second source. For the 47 possible sources we applied DARS. Hungary_LBA got the best rank among all sources, which geographically perfectly fits the scenario and also has the best *f3* value, where we suggest Bk-II admixture with local groups resulted in Bk-III. Serbia_Mokrin_EBA_Maros on the other hand has only a slightly lower rank, which, according to our previous results, did mix with Bk-II (or with a closely related group), and fits chronologically better to Bk-III. But also checking the following groups we can observe more possibilities, where outliers of Bk-II like admixture appeared. Nevertheless, Hungary_LBA likely is still the best representation of the admixing population, as this group represents the heterogeneous medium of Hungary during the Bronze Age, and does not restrict Bk-II mixture with one, but with multiple groups of local origin, which is a much more plausible scenario in our view. For results, see Supplementary Table 15.

Antonio, M.L., Gao, Z., Moots, H.M., Lucci, M., Candilio, F., Sawyer, S., Oberreiter, V., Calderon, D., Devitofranceschi, K., Aikens, R.C., Aneli, S., Bartoli, F., Bedini, A., Cheronet, O., Cotter, D.J.,

Fernandes, D.M., Gasperetti, G., Grifoni, R., Guidi, A., La Pastina, F., Loreti, E., Manacorda, D., Matullo, G., Morretta, S., Nava, A., Fiocchi Nicolai, V., Nomi, F., Pavolini, C., Pentiricci, M., Pergola, P., Piranomonte, M., Schmidt, R., Spinola, G., Sperduti, A., Rubini, M., Bondioli, L., Coppa, A., Pinhasi, R., Pritchard, J.K., 2019. Ancient Rome: A genetic crossroads of Europe and the Mediterranean. Science 366, 708–714. https://doi.org/10.1126/science.aay6826
Aufderheide, A.C., Rodríguez-Martín, C., 1998. The Cambridge encyclopedia of human paleopathology. Cambridge University Press, Cambridge.
Bandelt, H.J., Forster, P., Rohl, A., 1999. Median-joining networks for inferring intraspecific phylogenies. Mol. Biol. Evol. 16, 37–48. https://doi.org/10.1093/oxfordjournals.molbev.a026036
Bernert, Z., Évinger, S., Hajdu, T., 2007. New data on the biological age estimation of children using bone measurements based on historical populations from the Carpathian Basin. Ann. Hist.-Nat. MUSEI Natl. Hung. 99, 199–206.
Bóna, I., 1975. Die mittlere Bronzezeit Ungarns und ihre südöstlichen Beziehungen. Archaeol. Hung. 49, 73–76, 229–230.
Bóna, I., 1961. Geschichte der frühen und mittleren Bronzezeit in Ungarn und im mittleren Donauraum, in: Annales Universitatis Scientiarum Budapestinensis de Rolando Eötvös Nominatae III-IV. ELTE, Budapest, pp. 3–22.
Bondár M., 2020. Késő rézkori különleges temetkezések Balatonlelléről (Somogy megye). Kaposvári Rippl-Rónai Múz. Közleményei 89–108. https://doi.org/10.26080/krrmkozl.2020.7.89
Bondár, M., 2018. COMPLEX ANALYSES OF THE LATE COPPER AGE BURIALS IN THE CARPATHIAN BASIN 11.
Bondár, M., 2012. Prehistoric wagon models in the Carpathian basin (3500-1500 BC), Series minor. Archaeolingua, Budapest.
Bondár M., Demény A., Németh P., Karlik M., Fintor K., Tóth M., 2021. KÜLÖNLEGES „GAGÁT" GYÖNGY EGY KÜLÖNLEGES KÉSŐ RÉZKORI SÍRBÓL. Archeometriai Műh. 18, 143–156.
Bondár, M., Szécsényi-Nagy, A., 2020. Skull cult in the Late Copper Age. Ziridava 34, 91–104.
Brace, S., Diekmann, Y., Booth, T.J., van Dorp, L., Faltyskova, Z., Rohland, N., Mallick, S., Olalde, I., Ferry, M., Michel, M., Oppenheimer, J., Broomandkhoshbacht, N., Stewardson, K., Martiniano, R., Walsh, S., Kayser, M., Charlton, S., Hellenthal, G., Armit, I., Schulting, R., Craig, O.E., Sheridan, A., Parker Pearson, M., Stringer, C., Reich, D., Thomas, M.G., Barnes, I., 2019. Ancient genomes indicate population replacement in Early Neolithic Britain. Nat. Ecol. Evol. 3, 765–771. https://doi.org/10.1038/s41559-019-0871-9
Bronk Ramsey, C., 2009. Bayesian Analysis of Radiocarbon Dates. Radiocarbon 51, 337–360. https://doi.org/10.1017/S0033822200033865
Brück, J., Frieman, C.J., 2021. Making kin: The archaeology and genetics of human relationships. TATuP 30, 47–52. https://doi.org/10.14512/tatup.30.2.47
Cariaso, M., Lennon, G., 2012. SNPedia: a wiki supporting personal genome annotation, interpretation and analysis. Nucleic Acids Res. 40, D1308–D1312. https://doi.org/10.1093/nar/gkr798
Chaitanya, L., Breslin, K., Zuñiga, S., Wirken, L., Pośpiech, E., Kukla-Bartoszek, M., Sijen, T., Knijff, P. de, Liu, F., Branicki, W., Kayser, M., Walsh, S., 2018. The HIrisPlex-S system for eye, hair and skin colour prediction from DNA: Introduction and forensic developmental validation. Forensic Sci. Int. Genet. 35, 123–135. https://doi.org/10.1016/j.fsigen.2018.04.004
Chintalapati, M., Patterson, N., Moorjani, P., 2022. Reconstructing the spatiotemporal patterns of admixture during the European Holocene using a novel genomic dating method (preprint). Evolutionary Biology. https://doi.org/10.1101/2022.01.18.476710
ClinVar [WWW Document], 2021. . NCBI. URL https://www.ncbi.nlm.nih.gov/clinvar/
Dani, J., 2013. The Significance of Metallurgy at the Beginning of the Third Millennium BC in the Carpathian Basin., in: Transitions to the Bronze Age. Interregional Interaction and Socio-Cultural Change in the Third Millennium BC Carpathian Basin and Neighbouring Regions. Archaeolingua, pp. 203–231.
Dani, J., Fischl, K.P., Kulcsár, G., Szeverényi, V., Kiss, V., 2016. Visible and invisible inequality in Early and Middle Bronze Age Hungary., in: Arm Und Reich – Zur Ressourcenverteilung in Prähistorischen Gesellschaften. Rich and Poor – Competing for Resources in Prehistoric Societies., Mitteldeutscher Archäologentag. Halle (Saale), pp. 219–242.
Earle, T., Kulcsár, G., Kiss, V., Serlegi, G., Szeverényi, V., 2014. RECENT RESULTS FROM THE BRONZE AGE RESEARCHINTO THE BENTA VALLEY. Hung. Archaeol. E-J. 2014.
Fábián S., 2006. Õskori tömegsír Balatonkeresztúr-Réti-dûlõ lelõhelyrõl. Somogyi Múzeumok Közleményei 17, 79–88.

Fábián, S., Serlegi, G., 2009. Settlement and environment in the Late Copper Age along the southern shore of Lake Balaton in Hungary., in: Reimagining Regional Analyses: The Archaeology of Spatial and Social Dynamics. Cambridge Scholars Publishing, Newcastle, pp. 199–231.
Fazekas, I.G., Kósa, F., 1978. Forensic fetal osteology. Akadémiai Kiadó, Budapest.
Feldman, M., Fernández-Domínguez, E., Reynolds, L., Baird, D., Pearson, J., Hershkovitz, I., May, H., Goring-Morris, N., Benz, M., Gresky, J., Bianco, R.A., Fairbairn, A., Mustafaoğlu, G., Stockhammer, P.W., Posth, C., Haak, W., Jeong, C., Krause, J., 2019. Late Pleistocene human genome suggests a local origin for the first farmers of central Anatolia. Nat. Commun. 10, 1218. https://doi.org/10.1038/s41467-019-09209-7
Ferembach, D., Schwidetzky, I., Stloukal, M., 1979. Empfehlungen für die Alters- und Geschlechtsdiagnose am Skelett. Homo 30, 1–32.
Fischl, K.P., Kiss, V., Szeverényi, V., 2013. Transformations in the Carpathian Basin around 16oo B. C., in: Kultureller Umbruch Im Schatten Des Thera-Ausbruchs?(16oo – Cultural Change in the Shadow of TheThera-Eruption?)., Mitteldeutscher Archäologentag. Tagungen des Landesmuseums für Vorgeschichte Halle, Halle (Saale), pp. 355–371.
Freilich, S., Ringbauer, H., Los, D., Novak, M., Pavičić, D.T., Schiffels, S., Pinhasi, R., 2021. Reconstructing genetic histories and social organisation in Neolithic and Bronze Age Croatia. Sci. Rep. 11, 16729. https://doi.org/10.1038/s41598-021-94932-9
Fu, Q., Posth, C., Hajdinjak, M., Petr, M., Mallick, S., Fernandes, D., Furtwängler, A., Haak, W., Meyer, M., Mittnik, A., Nickel, B., Peltzer, A., Rohland, N., Slon, V., Talamo, S., Lazaridis, I., Lipson, M., Mathieson, I., Schiffels, S., Skoglund, P., Derevianko, A.P., Drozdov, N., Slavinsky, V., Tsybankov, A., Cremonesi, R.G., Mallegni, F., Gély, B., Vacca, E., Morales, M.R.G., Straus, L.G., Neugebauer-Maresch, C., Teschler-Nicola, M., Constantin, S., Moldovan, O.T., Benazzi, S., Peresani, M., Coppola, D., Lari, M., Ricci, S., Ronchitelli, A., Valentin, F., Thevenet, C., Wehrberger, K., Grigorescu, D., Rougier, H., Crevecoeur, I., Flas, D., Semal, P., Mannino, M.A., Cupillard, C., Bocherens, H., Conard, N.J., Harvati, K., Moiseyev, V., Drucker, D.G., Svoboda, J., Richards, M.P., Caramelli, D., Pinhasi, R., Kelso, J., Patterson, N., Krause, J., Pääbo, S., Reich, D., 2016. The genetic history of Ice Age Europe. Nature 534, 200–205. https://doi.org/10.1038/nature17993
Furtwängler, A., Rohrlach, A.B., Lamnidis, T.C., Papac, L., Neumann, G.U., Siebke, I., Reiter, E., Steuri, N., Hald, J., Denaire, A., Schnitzler, B., Wahl, J., Ramstein, M., Schuenemann, V.J., Stockhammer, P.W., Hafner, A., Lösch, S., Haak, W., Schiffels, S., Krause, J., 2020. Ancient genomes reveal social and genetic structure of Late Neolithic Switzerland. Nat. Commun. 11, 1915. https://doi.org/10.1038/s41467-020-15560-x
Gerasimov, M.M., 1949. Osnovy vosstanovleniya litsa po cherepu. [Bases of the craniofacial reconstruction.]. Nauka, Moskva, p. 185 pp.
Gouy, M., Guindon, S., Gascuel, O., 2010. SeaView Version 4: A Multiplatform Graphical User Interface for Sequence Alignment and Phylogenetic Tree Building. Mol. Biol. Evol. 27, 221–224. https://doi.org/10.1093/molbev/msp259
Günther, T., Malmström, H., Svensson, E.M., Omrak, A., Sánchez-Quinto, F., Kılınç, G.M., Krzewińska, M., Eriksson, G., Fraser, M., Edlund, H., Munters, A.R., Coutinho, A., Simões, L.G., Vicente, M., Sjölander, A., Jansen Sellevold, B., Jørgensen, R., Claes, P., Shriver, M.D., Valdiosera, C., Netea, M.G., Apel, J., Lidén, K., Skar, B., Storå, J., Götherström, A., Jakobsson, M., 2018. Population genomics of Mesolithic Scandinavia: Investigating early postglacial migration routes and high-latitude adaptation. PLOS Biol. 16, e2003703. https://doi.org/10.1371/journal.pbio.2003703
Haak, W., Lazaridis, I., Patterson, N., Rohland, N., Mallick, S., Llamas, B., Brandt, G., Nordenfelt, S., Harney, E., Stewardson, K., Fu, Q., Mittnik, A., Bánffy, E., Economou, C., Francken, M., Friederich, S., Pena, R.G., Hallgren, F., Khartanovich, V., Khokhlov, A., Kunst, M., Kuznetsov, P., Meller, H., Mochalov, O., Moiseyev, V., Nicklisch, N., Pichler, S.L., Risch, R., Rojo Guerra, M.A., Roth, C., Szécsényi-Nagy, A., Wahl, J., Meyer, M., Krause, J., Brown, D., Anthony, D., Cooper, A., Alt, K.W., Reich, D., 2015. Massive migration from the steppe was a source for Indo-European languages in Europe. Nature 522, 207–211. https://doi.org/10.1038/nature14317
Hajdu, T., György-Toronyi, A., Pap, I., Rosendahl, W., Szabó, G., 2016. The chronology and meaning of the Transdanubian encrusted pottery decoration. Praehistorische Z. 91, 353–368. https://doi.org/10.1515/pz-2016-0024
Honti S., Belényesy K., Fábián S., Gallina Z., Hajdú D.Á., Hansel B., Horváth T., Kiss V., Koós I., Marton T., Németh G.P., Oross K., Osztás A., Polgár P., Pásztókai-Szeőke J., Serlegi G., Siklósi Z., Sófalvi A., Virágos G., Sófalvi A., Virágos, 2004. A tervezett M7-esautópálya

Somogy megyei szakaszának megelőző régészeti feltárása (2002–2003) Előzetes jelentés III. Somogyi Múzeumok Közleményei 16, 3–70.
Honti S., Fábián S., Gallina Z., Hajdú D.Á., Hornok P., Koós I., Mersdorf Z., Molnár I., Németh G.P., Polgár P., Pásztókai-Szeőke J., Serlegi G., Siklósi Z., Sipos C., Somogyi K., 2006. Régészeti kutatások az M7-es autópálya Somogy megyei szakaszán és a 67-es úton (2004–2005) Elõzetes jelentés IV. Somogyi Múzeumok Közleményei 17, 64.
Işcan, M., Loth, S., Wright, R., 1985. Age estimation from the rib by phase analysis: white females. J. Forensic Sci. 30, 853–63.
Işcan, M., Loth, S., Wright, R., 1984. Age estimation from the rib by phase analysis: white males. J. Forensic Sci. 29, 1094–104.
K. Zoffmann, Z., 2000. Anthropological sketch of the prehistoric population of the Carpathian Basin. Acta Biol. Szeged. 44, 75–79.
Katoh, K., Standley, D.M., 2013. MAFFT Multiple Sequence Alignment Software Version 7: Improvements in Performance and Usability. Mol. Biol. Evol. 30, 772–780. https://doi.org/10.1093/molbev/mst010
Kiss, V., 2020a. The Bronze Age burial from Balatonakali revisited., in: Objects, Ideas and Travelers. Contacts between the Balkans, the Aegean and Western Anatolia during the Bronze Age and Early Iron Age. Conference to the Memory of Alexandru Vulpe. Deutsche Nationalbibliothek, Tulcea, pp. 553–568.
Kiss, V., 2020b. Transformations of Metal Supply during the Bronze Age in the Carpathian Basin. Hung. Hist. Rev. 9, 315–330. https://doi.org/10.38145/2020.2.315
Kiss, V., 2015. Recent data on chronology, distribution, and connections of Kisapostag, Transdanubian Encrusted Pottery and Litzenkeramik, in: KEĎ BRONZ VYSTRIEDAL MEĎ. Archaeologica Slovaca Monographiae, Slovakia, pp. 27–38.
Kiss, V., 2012. Middle Bronze Age encrusted pottery in western Hungary, Varia archaeologica Hungarica. Archaeolingua, Budapest.
Kiss, V., 2009. The Life Cycle of Middle Bronze Age Bronze Artefacts from the Western Part of the Carpathian Basin., in: Metals and Societies. Studies in Honour of Barbara S. Ottaway., Universitätsforschungen Zur Prähistorischen Archäologie. Bonn, pp. 328–335.
Kiss, V., Csányi, M., Dani, J., Fischl, K.P., Kulcsár, G., Szatmári, I., 2019. Chronology of the Early and Middle Bronze Age in Hungary: New results., in: Reinecke´s Heritage. Terminology, Chronology and Identity in Central Europe, 2300 and 1600 BC., Studia Hercynia. pp. 173–197.
Koeda, M., Watanabe, A., Tsuda, K., Matsumoto, M., Ikeda, Y., Kim, W., Tateno, A., Naing, B.T., Karibe, H., Shimada, T., Suzuki, H., Matsuura, M., Okubo, Y., 2015. Interaction effect between handedness and CNTNAP2 polymorphism (rs7794745 genotype) on voice-specific frontotemporal activity in healthy individuals: an fMRI study. Front. Behav. Neurosci. 9. https://doi.org/10.3389/fnbeh.2015.00087
Köhler, K., 2011. A harang alakú edények népe Szigetszentmiklós-Felső Ürge hegyi dűlő lelőhelyének embertani vizsgálati eredményei (Anthropological examination of the Bell Beaker cemetery at Szigetszentmiklós-Felső-Ürge-hegyi dűlő). Anthr. Közlemények 52, 55–76.
Köhler, K., 2009. The Anthropological Remains from the Budakalász Cemetery, in: The Copper Age Cemetery of Budakalász. Pytheas, Budapest, pp. 303–364.
Köhler K., 2006. Õskori tömegsír embertani leletei Balatonkeresztúrról. Somogyi Múzeumok Közleményei 17, 8.
Köhler, K., Marcsik, A., Zádori, P., Biro, G., Szeniczey, T., Fábián, S., Serlegi, G., Marton, T., Donoghue, H.D., Hajdu, T., 2017. Possible cases of leprosy from the Late Copper Age (3780-3650 cal BC) in Hungary. PLOS ONE 12, e0185966. https://doi.org/10.1371/journal.pone.0185966
Korneliussen, T.S., Albrechtsen, A., Nielsen, R., 2014. ANGSD: Analysis of Next Generation Sequencing Data. BMC Bioinformatics 15, 356. https://doi.org/10.1186/s12859-014-0356-4
Kulcsár, G., 2009. The beginnings of the Bronze Age in the Carpathian Basin: the Makó-Kosihy-Čaka and the Somogyvár-Vinkovci cultures in Hungary, Varia archaeologica Hungarica. Archaeolingua, Budapest.
Kustár, Á., Skultéty, G., 1996. A benepusztai honfoglaláskori férfi arcrekonstrukciója [Craniofacial reconstruction of a Hungarian conqueror male from the site of Benepuszta]. Savaria 22, 179–190.
Landrum, M.J., Lee, J.M., Benson, M., Brown, G.R., Chao, C., Chitipiralla, S., Gu, B., Hart, J., Hoffman, D., Jang, W., Karapetyan, K., Katz, K., Liu, C., Maddipatla, Z., Malheiro, A., McDaniel, K., Ovetsky, M., Riley, G., Zhou, G., Holmes, J.B., Kattman, B.L., Maglott, D.R.,

2018. ClinVar: improving access to variant interpretations and supporting evidence. Nucleic Acids Res. 46, D1062–D1067. https://doi.org/10.1093/nar/gkx1153
Lazaridis, I., Patterson, N., Mittnik, A., Renaud, G., Mallick, S., Kirsanow, K., Sudmant, P.H., Schraiber, J.G., Castellano, S., Lipson, M., Berger, B., Economou, C., Bollongino, R., Fu, Q., Bos, K.I., Nordenfelt, S., Li, H., de Filippo, C., Prüfer, K., Sawyer, S., Posth, C., Haak, W., Hallgren, F., Fornander, E., Rohland, N., Delsate, D., Francken, M., Guinet, J.-M., Wahl, J., Ayodo, G., Babiker, H.A., Bailliet, G., Balanovska, E., Balanovsky, O., Barrantes, R., Bedoya, G., Ben-Ami, H., Bene, J., Berrada, F., Bravi, C.M., Brisighelli, F., Busby, G.B.J., Cali, F., Churnosov, M., Cole, D.E.C., Corach, D., Damba, L., van Driem, G., Dryomov, S., Dugoujon, J.-M., Fedorova, S.A., Gallego Romero, I., Gubina, M., Hammer, M., Henn, B.M., Hervig, T., Hodoglugil, U., Jha, A.R., Karachanak-Yankova, S., Khusainova, R., Khusnutdinova, E., Kittles, R., Kivisild, T., Klitz, W., Kučinskas, V., Kushniarevich, A., Laredj, L., Litvinov, S., Loukidis, T., Mahley, R.W., Melegh, B., Metspalu, E., Molina, J., Mountain, J., Näkkäläjärvi, K., Nesheva, D., Nyambo, T., Osipova, L., Parik, J., Platonov, F., Posukh, O., Romano, V., Rothhammer, F., Rudan, I., Ruizbakiev, R., Sahakyan, H., Sajantila, A., Salas, A., Starikovskaya, E.B., Tarekegn, A., Toncheva, D., Turdikulova, S., Uktveryte, I., Utevska, O., Vasquez, R., Villena, M., Voevoda, M., Winkler, C.A., Yepiskoposyan, L., Zalloua, P., Zemunik, T., Cooper, A., Capelli, C., Thomas, M.G., Ruiz-Linares, A., Tishkoff, S.A., Singh, L., Thangaraj, K., Villems, R., Comas, D., Sukernik, R., Metspalu, M., Meyer, M., Eichler, E.E., Burger, J., Slatkin, M., Pääbo, S., Kelso, J., Reich, D., Krause, J., 2014. Ancient human genomes suggest three ancestral populations for present-day Europeans. Nature 513, 409–413. https://doi.org/10.1038/nature13673
Li, H., 2011. A statistical framework for SNP calling, mutation discovery, association mapping and population genetical parameter estimation from sequencing data. Bioinformatics 27, 2987–2993. https://doi.org/10.1093/bioinformatics/btr509
Li, H., Handsaker, B., Wysoker, A., Fennell, T., Ruan, J., Homer, N., Marth, G., Abecasis, G., Durbin, R., 1000 Genome Project Data Processing Subgroup, 2009. The Sequence Alignment/Map format and SAMtools. Bioinformatics 25, 2078–2079. https://doi.org/10.1093/bioinformatics/btp352
Lipson, M., Szécsényi-Nagy, A., Mallick, S., Pósa, A., Stégmár, B., Keerl, V., Rohland, N., Stewardson, K., Ferry, M., Michel, M., Oppenheimer, J., Broomandkhoshbacht, N., Harney, E., Nordenfelt, S., Llamas, B., Gusztáv Mende, B., Köhler, K., Oross, K., Bondár, M., Marton, T., Osztás, A., Jakucs, J., Paluch, T., Horváth, F., Csengeri, P., Koós, J., Sebők, K., Anders, A., Raczky, P., Regenye, J., Barna, J.P., Fábián, S., Serlegi, G., Toldi, Z., Gyöngyvér Nagy, E., Dani, J., Molnár, E., Pálfi, G., Márk, L., Melegh, B., Bánfai, Z., Domboróczki, L., Fernández-Eraso, J., Antonio Mujika-Alustiza, J., Alonso Fernández, C., Jiménez Echevarría, J., Bollongino, R., Orschiedt, J., Schierhold, K., Meller, H., Cooper, A., Burger, J., Bánffy, E., Alt, K.W., Lalueza-Fox, C., Haak, W., Reich, D., 2017. Parallel palaeogenomic transects reveal complex genetic history of early European farmers. Nature 551, 368–372. https://doi.org/10.1038/nature24476
Malmström, H., Günther, T., Svensson, E.M., Juras, A., Fraser, M., Munters, A.R., Pospieszny, Ł., Tõrv, M., Lindström, J., Götherström, A., Storå, J., Jakobsson, M., 2019. The genomic ancestry of the Scandinavian Battle Axe Culture people and their relation to the broader Corded Ware horizon. Proc. R. Soc. B Biol. Sci. 286, 20191528. https://doi.org/10.1098/rspb.2019.1528
Manchester, K., 1983. The archaeology of disease. University of Bradford Press.
Marciniak, S., Bergey, C.M., Silva, A.M., Hałuszko, A., Furmanek, M., Veselka, B., Velemínský, P., Vercellotti, G., Wahl, J., Zariņa, G., Longhi, C., Kolář, J., Garrido-Pena, R., Flores Fernández, R., Herrero-Corral, A.M., Simalcsik, A., Müller, W., Sheridan, A., Miliauskienė, Ž., Jankauskas, R., Moiseyev, V., Köhler, K., Király, Á., Gamarra, B., Cheronet, O., Szeverényi, V., Kiss, V., Szeniczey, T., Kiss, K., K. Zoffmann, Z., Koós, J., Hellebrandt, M., Domboróczki, L., Virag, C., Novak, M., Reich, D., Hajdu, T., von Cramon-Taubadel, N., Pinhasi, R., Perry, G.H., 2021. An integrative skeletal and paleogenomic analysis of prehistoric stature variation suggests relatively reduced health for early European farmers. BioRxiv Prepr. https://doi.org/10.1101/2021.03.31.437881
Massy, K., Friedrich, R., Mittnik, A., Stockhammer, P.W., 2022. Pedigree-based Bayesian modelling of radiocarbon dates. PLOS ONE 17, e0270374. https://doi.org/10.1371/journal.pone.0270374
Mathieson, I., Alpaslan-Roodenberg, S., Posth, C., Szécsényi-Nagy, A., Rohland, N., Mallick, S., Olalde, I., Broomandkhoshbacht, N., Candilio, F., Cheronet, O., Fernandes, D., Ferry, M., Gamarra, B., Fortes, G.G., Haak, W., Harney, E., Jones, E., Keating, D., Krause-Kyora, B.,

Kucukkalipci, I., Michel, M., Mittnik, A., Nägele, K., Novak, M., Oppenheimer, J., Patterson, N., Pfrengle, S., Sirak, K., Stewardson, K., Vai, S., Alexandrov, S., Alt, K.W., Andreescu, R., Antonović, D., Ash, A., Atanassova, N., Bacvarov, K., Gusztáv, M.B., Bocherens, H., Bolus, M., Boroneanţ, A., Boyadzhiev, Y., Budnik, A., Burmaz, J., Chohadzhiev, S., Conard, N.J., Cottiaux, R., Čuka, M., Cupillard, C., Drucker, D.G., Elenski, N., Francken, M., Galabova, B., Ganetsovski, G., Gély, B., Hajdu, T., Handzhyiska, V., Harvati, K., Higham, T., Iliev, S., Janković, I., Karavanić, I., Kennett, D.J., Komšo, D., Kozak, A., Labuda, D., Lari, M., Lazar, C., Leppek, M., Leshtakov, K., Vetro, D.L., Los, D., Lozanov, I., Malina, M., Martini, F., McSweeney, K., Meller, H., Menđušić, M., Mirea, P., Moiseyev, V., Petrova, V., Price, T.D., Simalcsik, A., Sineo, L., Šlaus, M., Slavchev, V., Stanev, P., Starović, A., Szeniczey, T., Talamo, S., Teschler-Nicola, M., Thevenet, C., Valchev, I., Valentin, F., Vasilyev, S., Veljanovska, F., Venelinova, S., Veselovskaya, E., Viola, B., Virag, C., Zaninović, J., Zäuner, S., Stockhammer, P.W., Catalano, G., Krauß, R., Caramelli, D., Zariņa, G., Gaydarska, B., Lillie, M., Nikitin, A.G., Potekhina, I., Papathanasiou, A., Borić, D., Bonsall, C., Krause, J., Pinhasi, R., Reich, D., 2018. The genomic history of southeastern Europe. Nature 555, 197–203. https://doi.org/10.1038/nature25778

Mathieson, I., Lazaridis, I., Rohland, N., Mallick, S., Patterson, N., Roodenberg, S.A., Harney, E., Stewardson, K., Fernandes, D., Novak, M., Sirak, K., Gamba, C., Jones, E.R., Llamas, B., Dryomov, S., Pickrell, J., Arsuaga, J.L., de Castro, J.M.B., Carbonell, E., Gerritsen, F., Khokhlov, A., Kuznetsov, P., Lozano, M., Meller, H., Mochalov, O., Moiseyev, V., Guerra, M.A.R., Roodenberg, J., Vergès, J.M., Krause, J., Cooper, A., Alt, K.W., Brown, D., Anthony, D., Lalueza-Fox, C., Haak, W., Pinhasi, R., Reich, D., 2015. Genome-wide patterns of selection in 230 ancient Eurasians. Nature 528, 499–503. https://doi.org/10.1038/nature16152

Metahuman Creator, 2021.

Mittnik, A., Massy, K., Knipper, C., Wittenborn, F., Friedrich, R., Pfrengle, S., Burri, M., Carlichi-Witjes, N., Deeg, H., Furtwängler, A., Harbeck, M., von Heyking, K., Kociumaka, C., Kucukkalipci, I., Lindauer, S., Metz, S., Staskiewicz, A., Thiel, A., Wahl, J., Haak, W., Pernicka, E., Schiffels, S., Stockhammer, P.W., Krause, J., 2019. Kinship-based social inequality in Bronze Age Europe. Science 366, 731–734. https://doi.org/10.1126/science.aax6219

Mittnik, A., Wang, C.-C., Pfrengle, S., Daubaras, M., Zariņa, G., Hallgren, F., Allmäe, R., Khartanovich, V., Moiseyev, V., Tõrv, M., Furtwängler, A., Andrades Valtueña, A., Feldman, M., Economou, C., Oinonen, M., Vasks, A., Balanovska, E., Reich, D., Jankauskas, R., Haak, W., Schiffels, S., Krause, J., 2018. The genetic prehistory of the Baltic Sea region. Nat. Commun. 9, 442. https://doi.org/10.1038/s41467-018-02825-9

Molnár, M., Janovics, R., Major, I., Orsovszki, J., Gönczi, R., Veres, M., Leonard, A.G., Castle, S.M., Lange, T.E., Wacker, L., Hajdas, I., Jull, A.J.T., 2013a. Status Report of the New AMS $^{14}$C Sample Preparation Lab of the Hertelendi Laboratory of Environmental Studies (Debrecen, Hungary). Radiocarbon 55, 665–676. https://doi.org/10.1017/S0033822200057829

Molnár, M., Rinyu, L., Veres, M., Seiler, M., Wacker, L., Synal, H.-A., 2013b. EnvironMICADAS: A Mini $^{14}$C AMS with Enhanced Gas Ion Source Interface in the Hertelendi Laboratory of Environmental Studies (HEKAL), Hungary. Radiocarbon 55, 338–344. https://doi.org/10.1017/S0033822200057453

Monroy Kuhn, J.M., Jakobsson, M., Günther, T., 2018. Estimating genetic kin relationships in prehistoric populations. PLOS ONE 13, e0195491. https://doi.org/10.1371/journal.pone.0195491

Müller-Scheeßel, N., 2013. ‚Irreguläre' Bestattungen in der Urgeschichte: Norm, Ritual, Strafe …? / Akten der Internationalen Tagung in Frankfurt a. M. vom 3. bis 5. Februar 2012., Kolloquien zur Vor- und Frühgeschichte. Deutsche Nationalbibliothek, Bonn.

Muthu, M., Prathibha, K., 2008. Management of a child with autism and severe bruxism: a case report. J Indian Soc Pedod Prev Dent 26(2):82–4. https://doi.org/10.4103/0970-4388.41623

Narasimhan, V.M., Patterson, N., Moorjani, P., Rohland, N., Bernardos, R., Mallick, S., Lazaridis, I., Nakatsuka, N., Olalde, I., Lipson, M., Kim, A.M., Olivieri, L.M., Coppa, A., Vidale, M., Mallory, J., Moiseyev, V., Kitov, E., Monge, J., Adamski, N., Alex, N., Broomandkhoshbacht, N., Candilio, F., Callan, K., Cheronet, O., Culleton, B.J., Ferry, M., Fernandes, D., Freilich, S., Gamarra, B., Gaudio, D., Hajdinjak, M., Harney, É., Harper, T.K., Keating, D., Lawson, A.M., Mah, M., Mandl, K., Michel, M., Novak, M., Oppenheimer, J., Rai, N., Sirak, K., Slon, V., Stewardson, K., Zalzala, F., Zhang, Z., Akhatov, G., Bagashev, A.N., Bagnera, A., Baitanayev, B., Bendezu-Sarmiento, J., Bissembaev, A.A., Bonora, G.L., Chargynov, T.T., Chikisheva, T., Dashkovskiy, P.K., Derevianko, A., Dobeš, M., Douka, K., Dubova, N., Duisengali, M.N., Enshin, D., Epimakhov, A., Fribus, A.V., Fuller, D., Goryachev, A., Gromov, A., Grushin, S.P.,

Hanks, B., Judd, M., Kazizov, E., Khokhlov, A., Krygin, A.P., Kupriyanova, E., Kuznetsov, P., Luiselli, D., Maksudov, F., Mamedov, A.M., Mamirov, T.B., Meiklejohn, C., Merrett, D.C., Micheli, R., Mochalov, O., Mustafokulov, S., Nayak, A., Pettener, D., Potts, R., Razhev, D., Rykun, M., Sarno, S., Savenkova, T.M., Sikhymbaeva, K., Slepchenko, S.M., Soltobaev, O.A., Stepanova, N., Svyatko, S., Tabaldiev, K., Teschler-Nicola, M., Tishkin, A.A., Tkachev, V.V., Vasilyev, S., Velemínský, P., Voyakin, D., Yermolayeva, A., Zahir, M., Zubkov, V.S., Zubova, A., Shinde, V.S., Lalueza-Fox, C., Meyer, M., Anthony, D., Boivin, N., Thangaraj, K., Kennett, D.J., Frachetti, M., Pinhasi, R., Reich, D., 2019. The formation of human populations in South and Central Asia. Science 365, eaat7487. https://doi.org/10.1126/science.aat7487

Network Software [WWW Document], 2008. . Fluxus-Eng. URL https://www.fluxus-engineering.com/sharepub.htm#a1

Olalde, I., Brace, S., Allentoft, M.E., Armit, I., Kristiansen, K., Booth, T., Rohland, N., Mallick, S., Szécsényi-Nagy, A., Mittnik, A., Altena, E., Lipson, M., Lazaridis, I., Harper, T.K., Patterson, N., Broomandkhoshbacht, N., Diekmann, Y., Faltyskova, Z., Fernandes, D., Ferry, M., Harney, E., de Knijff, P., Michel, M., Oppenheimer, J., Stewardson, K., Barclay, A., Alt, K.W., Liesau, C., Ríos, P., Blasco, C., Miguel, J.V., García, R.M., Fernández, A.A., Bánffy, E., Bernabò-Brea, M., Billoin, D., Bonsall, C., Bonsall, L., Allen, T., Büster, L., Carver, S., Navarro, L.C., Craig, O.E., Cook, G.T., Cunliffe, B., Denaire, A., Dinwiddy, K.E., Dodwell, N., Ernée, M., Evans, C., Kuchařík, M., Farré, J.F., Fowler, C., Gazenbeek, M., Pena, R.G., Haber-Uriarte, M., Haduch, E., Hey, G., Jowett, N., Knowles, T., Massy, K., Pfrengle, S., Lefranc, P., Lemercier, O., Lefebvre, A., Martínez, C.H., Olmo, V.G., Ramírez, A.B., Maurandi, J.L., Majó, T., McKinley, J.I., McSweeney, K., Mende, B.G., Modi, A., Kulcsár, G., Kiss, V., Czene, A., Patay, R., Endrődi, A., Köhler, K., Hajdu, T., Szeniczey, T., Dani, J., Bernert, Z., Hoole, M., Cheronet, O., Keating, D., Velemínský, P., Dobeš, M., Candilio, F., Brown, F., Fernández, R.F., Herrero-Corral, A.-M., Tusa, S., Carnieri, E., Lentini, L., Valenti, A., Zanini, A., Waddington, C., Delibes, G., Guerra-Doce, E., Neil, B., Brittain, M., Luke, M., Mortimer, R., Desideri, J., Besse, M., Brücken, G., Furmanek, M., Hałuszko, A., Mackiewicz, M., Rapiński, A., Leach, S., Soriano, I., Lillios, K.T., Cardoso, J.L., Pearson, M.P., Włodarczak, P., Price, T.D., Prieto, P., Rey, P.-J., Risch, R., Rojo Guerra, M.A., Schmitt, A., Serralongue, J., Silva, A.M., Smrčka, V., Vergnaud, L., Zilhão, J., Caramelli, D., Higham, T., Thomas, M.G., Kennett, D.J., Fokkens, H., Heyd, V., Sheridan, A., Sjögren, K.-G., Stockhammer, P.W., Krause, J., Pinhasi, R., Haak, W., Barnes, I., Lalueza-Fox, C., Reich, D., 2018. The Beaker phenomenon and the genomic transformation of northwest Europe. Nature 555, 190–196. https://doi.org/10.1038/nature25738

Ortner, D., 2003. Identification of Pathological Conditions in Human Skeletal Remains. London-New York Academic Press.

Patterson, N., Isakov, M., Booth, T., Büster, L., Fischer, C.-E., Olalde, I., Ringbauer, H., Akbari, A., Cheronet, O., Bleasdale, M., Adamski, N., Altena, E., Bernardos, R., Brace, S., Broomandkhoshbacht, N., Callan, K., Candilio, F., Culleton, B., Curtis, E., Demetz, L., Carlson, K.S.D., Edwards, C.J., Fernandes, D.M., Foody, M.G.B., Freilich, S., Goodchild, H., Kearns, A., Lawson, A.M., Lazaridis, I., Mah, M., Mallick, S., Mandl, K., Micco, A., Michel, M., Morante, G.B., Oppenheimer, J., Özdoğan, K.T., Qiu, L., Schattke, C., Stewardson, K., Workman, J.N., Zalzala, F., Zhang, Z., Agustí, B., Allen, T., Almássy, K., Amkreutz, L., Ash, A., Baillif-Ducros, C., Barclay, A., Bartosiewicz, L., Baxter, K., Bernert, Z., Blažek, J., Bodružić, M., Boissinot, P., Bonsall, C., Bradley, P., Brittain, M., Brookes, A., Brown, F., Brown, L., Brunning, R., Budd, C., Burmaz, J., Canet, S., Carnicero-Cáceres, S., Čaušević-Bully, M., Chamberlain, A., Chauvin, S., Clough, S., Čondić, N., Coppa, A., Craig, O., Črešnar, M., Cummings, V., Czifra, S., Danielisová, A., Daniels, R., Davies, A., de Jersey, P., Deacon, J., Deminger, C., Ditchfield, P.W., Dizdar, M., Dobeš, M., Dobisíková, M., Domboróczki, L., Drinkall, G., Đukić, A., Ernée, M., Evans, C., Evans, J., Fernández-Götz, M., Filipović, S., Fitzpatrick, A., Fokkens, H., Fowler, C., Fox, A., Gallina, Z., Gamble, M., González Morales, M.R., González-Rabanal, B., Green, A., Gyenesei, K., Habermehl, D., Hajdu, T., Hamilton, D., Harris, J., Hayden, C., Hendriks, J., Hernu, B., Hey, G., Horňák, M., Ilon, G., Istvánovits, E., Jones, A.M., Kavur, M.B., Kazek, K., Kenyon, R.A., Khreisheh, A., Kiss, V., Kleijne, J., Knight, M., Kootker, L.M., Kovács, P.F., Kozubová, A., Kulcsár, G., Kulcsár, V., Le Pennec, C., Legge, M., Leivers, M., Loe, L., López-Costas, O., Lord, T., Los, D., Lyall, J., Marín-Arroyo, A.B., Mason, P., Matošević, D., Maxted, A., McIntyre, L., McKinley, J., McSweeney, K., Meijlink, B., Mende, B.G., Menđušić, M., Metlička, M., Meyer, S., Mihovilić, K., Milasinovic, L., Minnitt, S., Moore, J., Morley, G., Mullan, G., Musilová, M., Neil, B., Nicholls, R., Novak, M., Pala, M., Papworth, M., Paresys, C., Patten, R., Perkić, D., Pesti, K., Petit, A., Petriščáková, K., Pichon,

C., Pickard, C., Pilling, Z., Price, T.D., Radović, S., Redfern, R., Resutík, B., Rhodes, D.T., Richards, M.B., Roberts, A., Roefstra, J., Sankot, P., Šefčáková, A., Sheridan, A., Skae, S., Šmolíková, M., Somogyi, K., Somogyvári, Á., Stephens, M., Szabó, G., Szécsényi-Nagy, A., Szeniczey, T., Tabor, J., Tankó, K., Maria, C.T., Terry, R., Teržan, B., Teschler-Nicola, M., Torres-Martínez, J.F., Trapp, J., Turle, R., Ujvári, F., van der Heiden, M., Veleminsky, P., Veselka, B., Vytlačil, Z., Waddington, C., Ware, P., Wilkinson, P., Wilson, L., Wiseman, R., Young, E., Zaninović, J., Žitňan, A., Lalueza-Fox, C., de Knijff, P., Barnes, I., Halkon, P., Thomas, M.G., Kennett, D.J., Cunliffe, B., Lillie, M., Rohland, N., Pinhasi, R., Armit, I., Reich, D., 2022. Large-scale migration into Britain during the Middle to Late Bronze Age. Nature 601, 588–594. https://doi.org/10.1038/s41586-021-04287-4

Patterson, N., Moorjani, P., Luo, Y., Mallick, S., Rohland, N., Zhan, Y., Genschoreck, T., Webster, T., Reich, D., 2012. Ancient Admixture in Human History. Genetics 192, 1065–1093. https://doi.org/10.1534/genetics.112.145037

Patterson, N., Price, A.L., Reich, D., 2006. Population Structure and Eigenanalysis. PLoS Genet. 2, e190. https://doi.org/10.1371/journal.pgen.0020190

Petr, M., Vernot, B., Kelso, J., 2019. admixr—R package for reproducible analyses using ADMIXTOOLS. Bioinformatics 35, 3194–3195. https://doi.org/10.1093/bioinformatics/btz030

Pfeifer, B., Wittelsbürger, U., Ramos-Onsins, S.E., Lercher, M.J., 2014. PopGenome: An Efficient Swiss Army Knife for Population Genomic Analyses in R. Mol. Biol. Evol. 31, 1929–1936. https://doi.org/10.1093/molbev/msu136

Prag, J., Neave, R., 1997. Making faces: Using forensic and archaeological evidence, in: British Museum Press. London, p. 256 pp.

Preda-Balanica, B., Frinculeasa, A., Heyd, V., 2020. The Yamnaya Impact North of the Lower Danube - A Tale of Newcomers and Locals. Bull. Société Préhistorique Francaise 117, 85–101.

Quinlan, A.R., Hall, I.M., 2010. BEDTools: a flexible suite of utilities for comparing genomic features. Bioinformatics 26, 841–842. https://doi.org/10.1093/bioinformatics/btq033

Ralf, A., Montiel González, D., Zhong, K., Kayser, M., 2018. Yleaf: Software for Human Y-Chromosomal Haplogroup Inference from Next-Generation Sequencing Data. Mol. Biol. Evol. 35, 1291–1294. https://doi.org/10.1093/molbev/msy032

Reich, D., 2021. AADR - Allen Ancient DNA Resource [WWW Document]. URL https://reich.hms.harvard.edu/allen-ancient-dna-resource-aadr-downloadable-genotypes-present-day-and-ancient-dna-data

Reich, D., 2018. Who we are and how we got there: Ancient DNA and the new science of the human past. Oxford University Press, Oxford.

Reimer, P.J., Austin, W.E.N., Bard, E., Bayliss, A., Blackwell, P.G., Bronk Ramsey, C., Butzin, M., Cheng, H., Edwards, R.L., Friedrich, M., Grootes, P.M., Guilderson, T.P., Hajdas, I., Heaton, T.J., Hogg, A.G., Hughen, K.A., Kromer, B., Manning, S.W., Muscheler, R., Palmer, J.G., Pearson, C., van der Plicht, J., Reimer, R.W., Richards, D.A., Scott, E.M., Southon, J.R., Turney, C.S.M., Wacker, L., Adolphi, F., Büntgen, U., Capano, M., Fahrni, S.M., Fogtmann-Schulz, A., Friedrich, R., Köhler, P., Kudsk, S., Miyake, F., Olsen, J., Reinig, F., Sakamoto, M., Sookdeo, A., Talamo, S., 2020. The IntCal20 Northern Hemisphere Radiocarbon Age Calibration Curve (0–55 cal kBP). Radiocarbon 62, 725–757. https://doi.org/10.1017/RDC.2020.41

Rivollat, M., Jeong, C., Schiffels, S., Küçükkalıpçı, İ., Pemonge, M.-H., Rohrlach, A.B., Alt, K.W., Binder, D., Friederich, S., Ghesquière, E., Gronenborn, D., Laporte, L., Lefranc, P., Meller, H., Réveillas, H., Rosenstock, E., Rottier, S., Scarre, C., Soler, L., Wahl, J., Krause, J., Deguilloux, M.-F., Haak, W., 2020. Ancient genome-wide DNA from France highlights the complexity of interactions between Mesolithic hunter-gatherers and Neolithic farmers. Sci. Adv. 6, eaaz5344. https://doi.org/10.1126/sciadv.aaz5344

Röhrer-Ertl, O., Helmer, R., 1984. Zu Stand und Möglichkeiten der Erneut modifizierten Kollmann–Methode. (Gesichtsrekonstruktion aufgrund des Schädels.), in: Gegenbaurs Morphologisches Jahrbuch. pp. 369–373.

Ronquist, F., Huelsenbeck, J.P., 2003. MrBayes 3: Bayesian phylogenetic inference under mixed models. Bioinformatics 19, 1572–1574. https://doi.org/10.1093/bioinformatics/btg180

Rynn, C., Wilkinson, C.M., Peters, H.L., 2010. Prediction of nasal morphology from the skull. Forensic Sci. Med. Pathol. 6, 20–34. https://doi.org/10.1007/s12024-009-9124-6

Sánchez-Quinto, F., Malmström, H., Fraser, M., Girdland-Flink, L., Svensson, E.M., Simões, L.G., George, R., Hollfelder, N., Burenhult, G., Noble, G., Britton, K., Talamo, S., Curtis, N., Brzobohata, H., Sumberova, R., Götherström, A., Storå, J., Jakobsson, M., 2019. Megalithic tombs in western and northern Neolithic Europe were linked to a kindred society. Proc. Natl.

Acad. Sci. 116, 9469–9474. https://doi.org/10.1073/pnas.1818037116
Sedig, J.W., Olalde, I., Patterson, N., Harney, É., Reich, D., 2021. Combining ancient DNA and radiocarbon dating data to increase chronological accuracy. J. Archaeol. Sci. 133, 105452. https://doi.org/10.1016/j.jas.2021.105452
Shribman, S., Reid, E., Crosby, A.H., Houlden, H., Warner, T.T., 2019. Hereditary spastic paraplegia: from diagnosis to emerging therapeutic approaches. Lancet Neurol. 18, 1136–1146. https://doi.org/10.1016/S1474-4422(19)30235-2
Sjøvold, T., 1981. Árpás anatomical method for face reconstruction, in: Ossa. pp. 203–204.
Somogyi, K., 2004. A kisapostagi kultúra birituális temetője Ordacsehi-Csereföldön – Das birituelle Gräberfeld der Kisapostag-Kultur on Ordacsehi-Csereföld, in: Őskoros Kutatók III. Összejövetelének Konferenciakötete. Szombathely, pp. 349–381.
Stloukal, M., Hanáková, H., 1978. Die Lange der Langsknochen altslawischer Bevölkerungen unter besonderer Berücksichtigung von Wachstumsfragen. Homo 29, 53–69.
Suchard, M.A., Lemey, P., Baele, G., Ayres, D.L., Drummond, A.J., Rambaut, A., 2018. Bayesian phylogenetic and phylodynamic data integration using BEAST 1.10. Virus Evol. 4. https://doi.org/10.1093/ve/vey016
Taylor, K.T., 2001. Forensic art and illustration. CRC Press.
Torma, I., 1978. A balatonakali bronzkori sír (Das bronzezeitliche Grab inBalatonakali). Veszprémi Megyei Múzeumok Közleményei 13, 15–24.
Ubelaker, D.H., 1989. Human skeletal remains, excavation, analysis, interpretation. Taraxacum, Washington DC.
Wang, J., 2017. Estimating pairwise relatedness in a small sample of individuals. Heredity 119, 302–313. https://doi.org/10.1038/hdy.2017.52
Y-DNA Haplogroup Tree 2019-2020 [WWW Document], 2019. . Int. Soc. Genet. Geneal. URL https://isogg.org/tree/
Žegarac, A., Winkelbach, L., Blöcher, J., Diekmann, Y., Krečković Gavrilović, M., Porčić, M., Stojković, B., Milašinović, L., Schreiber, M., Wegmann, D., Veeramah, K.R., Stefanović, S., Burger, J., 2021. Ancient genomes provide insights into family structure and the heredity of social status in the early Bronze Age of southeastern Europe. Sci. Rep. 11, 10072. https://doi.org/10.1038/s41598-021-89090-x
